# Kinetics and mechanisms of catalyzed dual-E (antithetic) controllers

**DOI:** 10.1101/2021.12.26.474198

**Authors:** Qaiser Waheed, Huimin Zhou, Peter Ruoff

## Abstract

Homeostasis plays a central role in our understanding how cells and organisms are able to oppose environmental disturbances and thereby maintain an internal stability. During the last two decades there has been an increased interest in using control engineering methods, especially integral control, in the analysis and design of homeostatic networks. Several reaction kinetic mechanisms have been discovered which lead to integral control. In two of them integral control is achieved, either by the removal of a single control species E by zero-order kinetics (”single-E controllers”), or by the removal of two control species by second-order kinetics (”antithetic or dual-E control”). In this paper we show results when the control species E_1_ and E_2_ in antithetic control are removed enzymatically by ping-pong or ternary-complex mechanisms. Our findings show that enzyme-catalyzed dual-E controllers can work in two control modes. In one mode, one of the two control species is active, but requires zero-order kinetics in its removal. In the other mode, both controller species are active and both are removed enzymatically. Conditions for the two control modes are put forward and biochemical examples with the structure of enzyme-catalyzed dual-E controllers are discussed.

## Introduction

During the last twenty years there has been an increasing interest in the design of molecular models that can exhibit integral control and show robust homeostasis/perfect adaptation. [1–11]. Integral control, which is part of many industrial regulation processes works in the following way (Fig 1): the controlled variable A, outlined in blue, is compared with the controller’s set-point A_set_ (shown in red). The difference (or error) between A_set_ and the actual value of A, *ϵ*=*A_set_ − A*, is calculated and integrated in time. The time integral of *ϵ*, described as the variable E, is then used to correct for perturbations acting on A. It can be shown that for step-wise perturbations an integral feedback will move A precisely to A_set_ [3].

**Fig 1.**
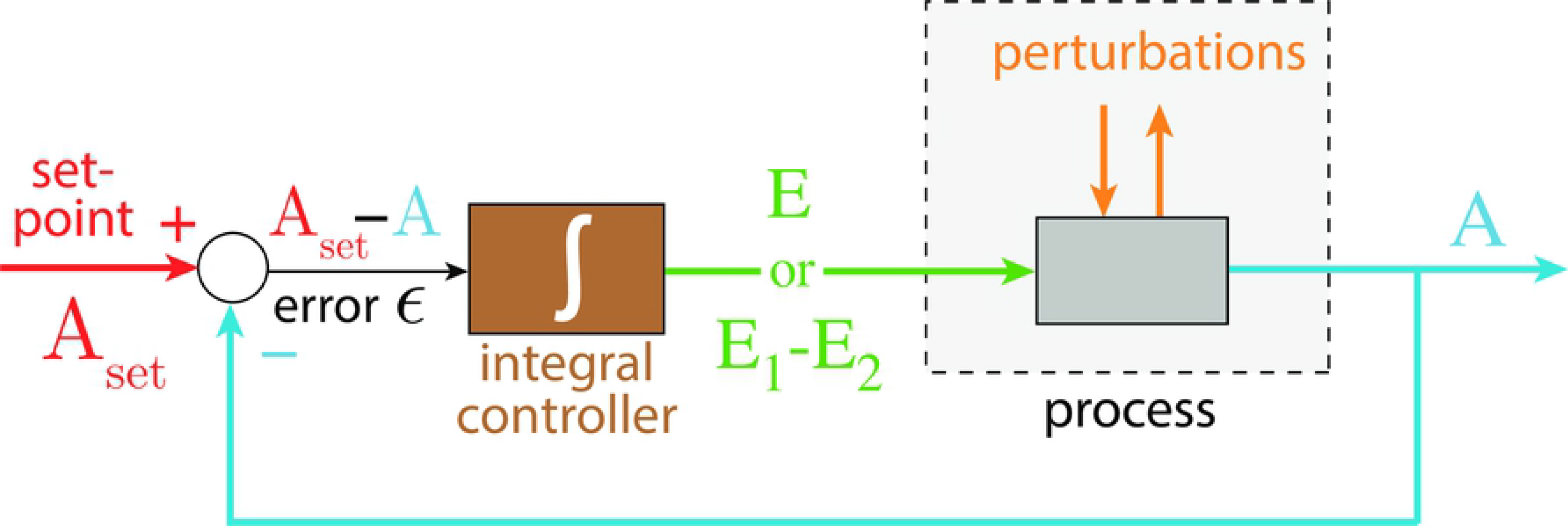
The concept of integral control. In single-E controllers the variable E is proportional to the integrated error *ϵ*, *∫ ϵdt*, which is used to correct for perturbations in A. In dual-E (antithetic) controllers the difference between variables E_1_ and E_2_ is proportional to the integrated error (see text). In both cases integral control will move A precisely to its set-point A_set_ when A is perturbed by step-wise perturbations [3].

Mustafa Khammash’s group recently suggested an interesting alternative approach, termed antithetic control, where instead of one controller molecule E there are two (E_1_ and E_2_) [7, 8, 10, 11]. In the single-E control case the condition of integral control is given by

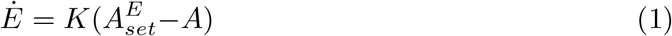

where K is a constant.

In the antithetic/dual-E case integral control is achieved by

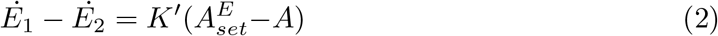

with *K*′ being a constant. Fig 2 shows, as an example, how integral control in single- and dual-E controllers can be achieved in a negative feedback structure termed motif 5. Motif 5, an outflow controller, is one of eight basic negative feedback structures, which divide equally into two sets of inflow and outflow controllers [6]. Briefly, in inflow controllers the compensatory flux opposes an uncontrolled removal of the controlled variable (here A), while in outflow controllers an uncontrolled inflow of the controlled variables is compensated.

**Fig 2.**
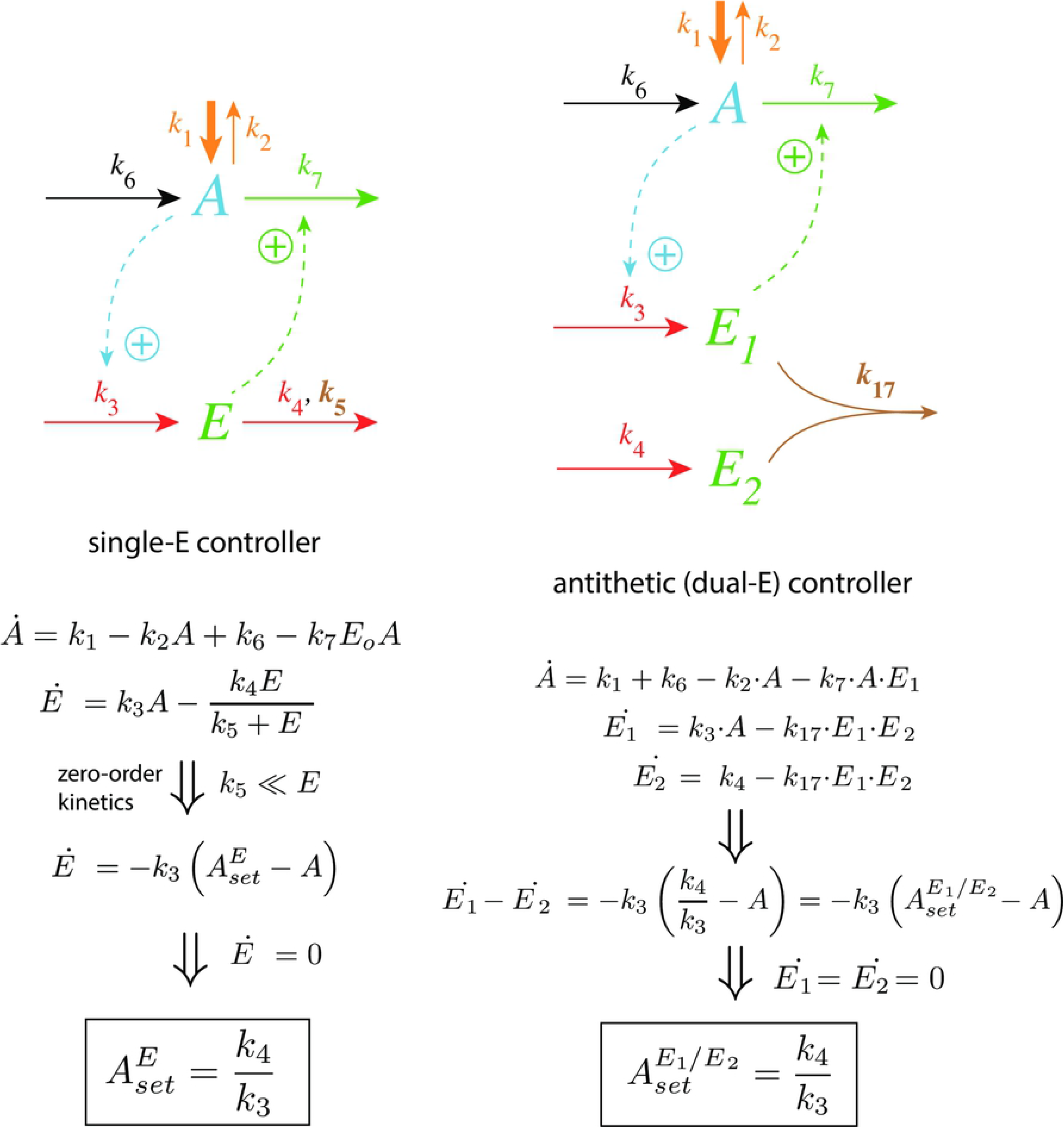
Single-E and dual-E (antithetic) representations of integral control using a motif 5 negative feedback structure. Left panel: Single-E controller where error integration occurs by zero-order kinetics (low *k*_5_) removing *E* [4, 6]. Right panel: Dual-E controller [7, 8, 10] with controller pairs E_1_ and E_2_. Error integration occurs by the (here second-order) reaction between *E*_1_ and *E*_2_. In the single-E controller the concentration of E is proportional to the integrated error 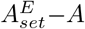. In the antithetic (dual-E) controller, the difference *E*_1_−*E*_2_ is proportional to the integrated error 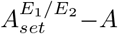. The colorings of the reaction schemes relate to the different parts in the general control loop shown in Fig 1.

As indicated in Fig 2, left panel, and by Eq 1 the steady state condition of E (*Ė* = 0) for a single-E controller determines its set-point. Since the antithetic controller is based on a reaction between *E*_1_ and *E*_2_ with speed *v* and rate constant *k*_17_ (Fig 2, right panel), i.e.,

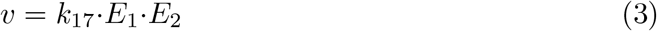

the set-point for this controller is determined by the difference of the steady state conditions between *E*_1_ and *E*_2_ (Eq 2).

## Aim of this work

As practically all processes within a cell are catalyzed by enzymes, we asked the question what influence enzymes may have on dual-E controllers, specifically when the reaction between controller species *E*_1_ and *E*_2_ is catalyzed. We here show the behaviors of a set of catalyzed antithetic/dual-E controllers. The enzymatic mechanisms for the removal of *E*_1_ and *E*_2_ include ping-pong, as well as random-order and compulsory order ternary-complex mechanisms [12, 13]. The role of total enzyme concentration is investigated and how the negative feedback structure of the motifs influence controller performance. Fig 3 shows the incorporation of dual-E integral control into the eight negative feedback motifs [6] with enzyme *Ez* catalyzing the reaction between *E*_1_ and *E*_2_.

**Fig 3.**
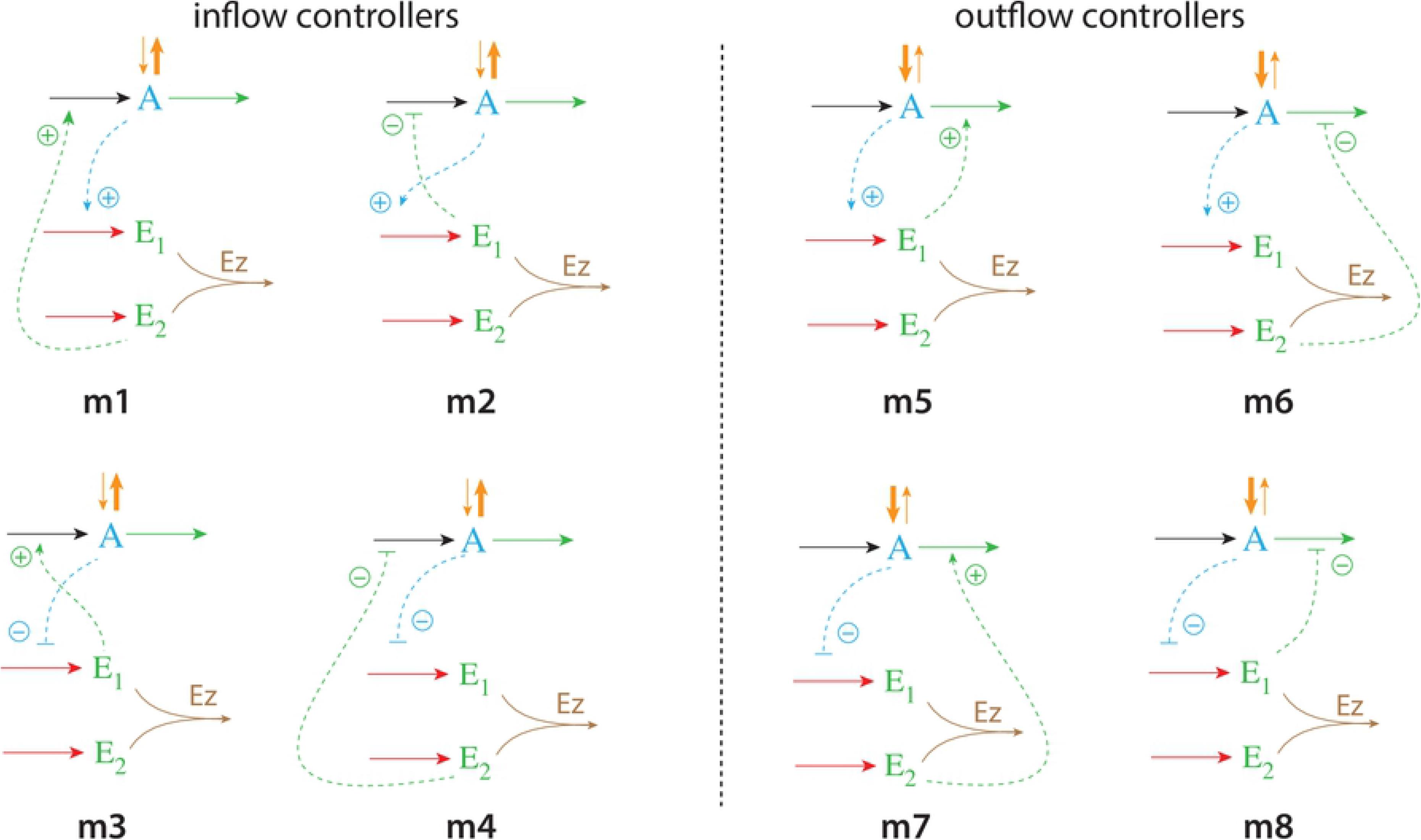
Dual-E (antithetic) integral control in combination with the eight negative feedback structures m1-m8. In the calculations the removal of *E*_1_ and *E*_2_ is catalyzed by enzyme Ez using different mechanisms. The signaling between *A* and the manipulated variables *E*_1_/*E*_2_ occurs either by an “inner loop” between *A* and *E*_1_ (motifs m2, m3, m5, and m8), or by an “outer-loop” signaling between *A* and *E*_2_ (motifs m1, m4, m6, and m7).

We will show that the performance of the catalyzed dual-E controllers, like response time, depends to a certain degree on the feedback structure/motif and on the enzymatic processing mechanism of *E*_1_ and *E*_2_. In comparison with single-E control [4, 6] the enzymatic dual-E controllers have the advantage that robust homeostasis is not bound to the requirement of zero-order kinetics, but can also work in its presence.

## Materials and methods

Computations were performed by using the Fortran subroutine LSODE [14]. Plots were generated with gnuplot (www.gnuplot.info) and edited with Adobe Illustrator (adobe.com). To make notations simpler, concentrations of compounds are generally denoted by compound names without square brackets. Time derivatives are indicated by the ‘dot’ notation. Concentrations and rate parameter values are given in arbitrary units (au).

### Enzymatic mechanisms considered

There are two major mechanisms [12, 13] when *E*_1_ and *E*_2_ are processed by an enzyme Ez, i.e.,

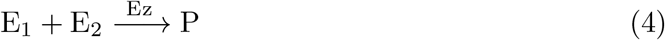

In one of them, a ternary complex E_1_·Ez·E_2_ between enzyme and substrates E_1_ and E_2_ is formed, either via a random binding order (Fig 4a) or by a compulsory binding order (Fig 4b). The other mechanism, termed “ping-pong”, contains two compulsory order binding events. During the first step one of the substrates E_1_ or E_2_ binds to the enzyme Ez, releases a possible first product and creates an alternative enzymatic form Ez^*^, which is able to bind the second substrate. In the final step the enzymatic species Ez is regenerated and a possible second product is released. A new enzymatic cycle can start again (Fig 4c).

**Fig 4.**
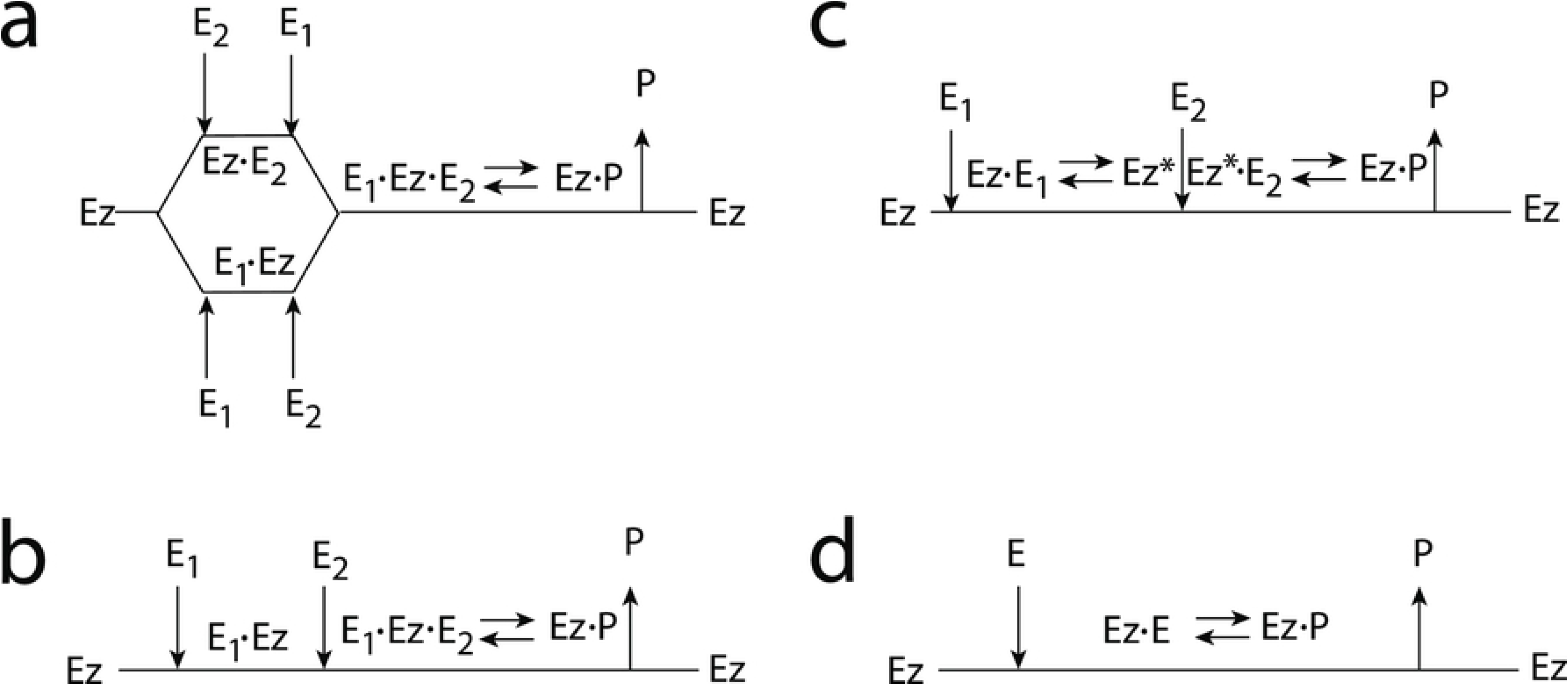
Overview (Cleland notation [16]) of the enzymatic mechanisms removing *E*_1_ and *E*_2_ (Eq 4). (a) Ternary complex mechanism with random binding of *E*_1_ and *E*_2_ to the enzyme. (b) Ternary complex mechanism with compulsory binding order. Here *E*_1_ binds first to free enzyme *Ez* then *E*_2_ binds to the *E*_1_·*Ez* complex. Alternatively, *E*_2_ can bind first to *Ez* and then *E*_1_ to form the ternary complex. (c) Ping-pong mechanism. *E*_1_ (or *E*_2_) bind first to *Ez* leading to the alternate enzyme form *Ez*^*^, which then can bind *E*_2_ (or *E*_1_). (d) Single-substrate Michaelis-Menten mechanism used in single-E controllers.

In the case of single-E controllers E is removed by enzyme Ez

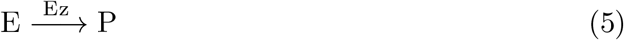

by using (single-substrate) Michaelis-Menten kinetics (Fig 4d). Although single-E controllers have already been analyzed to a large extent before [4, 6, 15], we will encounter their catalyzed versions also here, because some of the dual-E controllers can switch between single-E and dual-E control mode.

We noted that a necessary condition for robust homeostasis to occur is that the involved negative feedback loops need to be described as irreversible processes. Therefore, the enzymatic reactions in Eqs 4 and 5 need to be irreversible. Already in 1925 Lotka [17] investigated whether certain biological phenomena, such as oscillations and homeostasis, could be based on Le Chatelier’s principle, since at that time biologists attempted to apply the principle to biological systems [18]. Lotka concluded in the negative. Today we regard life as an overall irreversible process, a “dissipative structure” being far from chemical equilibrium [19, 20] and which allows for self-maintenance [21].

For each of the three mechanisms in Fig 4a-c steady state expressions for *v* of reaction 4 have been found numerically with LSODE and by using the King-Altman method [22] (S1 Text). The King-Altman method has the advantage that *v* can be expressed as an analytical function of the concentrations of *E*_1_ and *E*_2_ and the other rate parameters. Our calculations showed that the steady state expressions of *v* were always in excellent agreement with the corresponding numerical results.

### Feedback motifs considered

From the eight feedback structures of Fig 3 we have analyzed four of them: two of the four “inner-loop” motifs m2 and m5 and the two “outer-loop” motifs m4 and m7. The remaining four motifs have similar feedback symmetries and we do not expect significant differences to those considered here.

## Results

For each of the motifs m2, m4, m5, and m7 we describe how the controllers perform under step-wise perturbations when the above mentioned ternary-complex and ping-pong mechanisms are applied to remove *E*_1_ and *E*_2_.

### Controllers based on motif 2

This motif’s performance has been found to be remarkably good, especially with respect to perturbations which increase their strength with time [23, 24]. Motif 2 is an inflow type of controller which opposes outflow perturbations in the controlled variable.

#### Motif 2 dual-E controller removing *E*_1_ and *E*_2_ by an enzymatic random-order ternary-complez mechanism

Fig 5 shows the reaction scheme when *E*_1_ and *E*_2_ are removed enzymatically by using a ternary-complex mechanism with random binding order and E_1_ as the derepressing agent.

**Fig 5.**
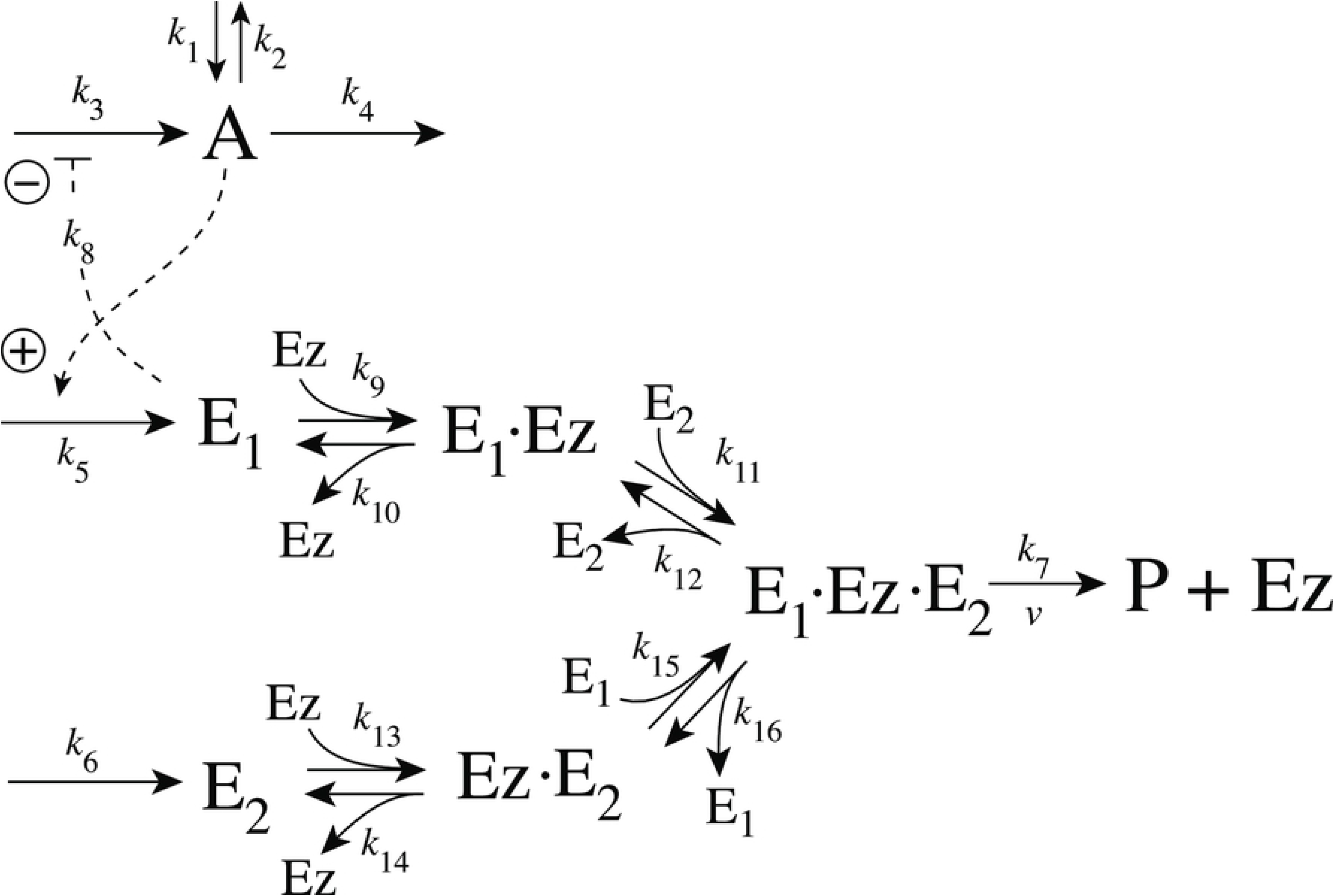
Motif 2 antithetic controller: removal of E_1_ and E_2_ by enzyme Ez using a ternary-complex mechanism with random binding order.

The rate equations are:

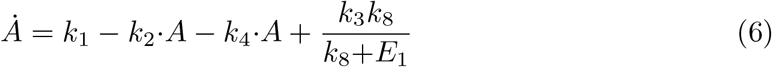

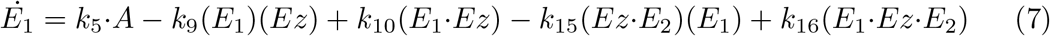

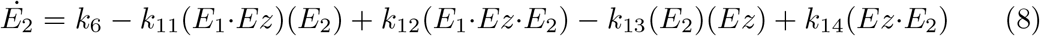

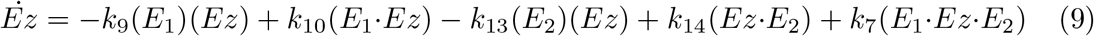

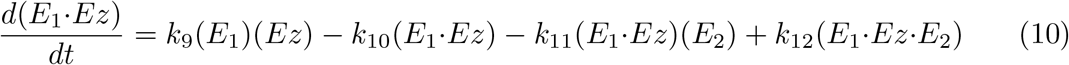

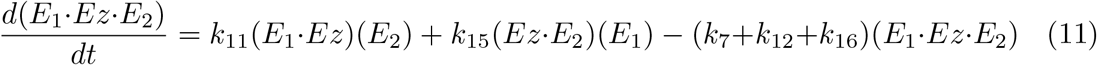

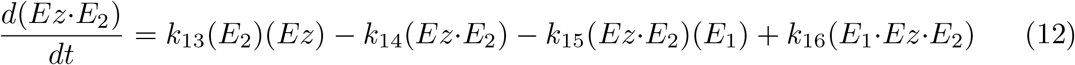

An analytical expression for the reaction velocity

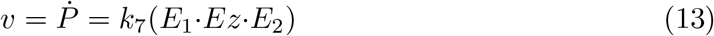

can be obtained by the steady state approximation (S1 Text), which has been found (see below) to be in excellent agreement with the numerical results.

We observed that the enzymatic controller in Fig 5 can show two set-points of A. One is given by

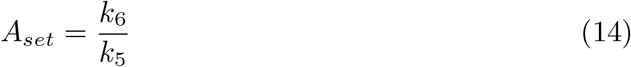

when both E_1_ and E_2_ participate in the regulation of A (dual-E control).

The other set-point is given by

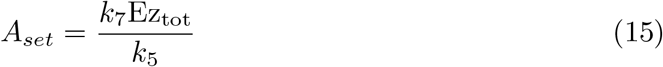

In this case only E_1_ participates in the control of A. (single-E control). The switching between the two control modes is described in more detail below.

#### Motif 2 single-E controller with Michaelis-Menten removal of E

Due to the above indicated switch between catalyzed dual-E and single-E control mode we here show the catalyzed single-E m2 controller (Fig 6), which will also be compared with the catalyzed m2 dual-E controller.

**Fig 6.**
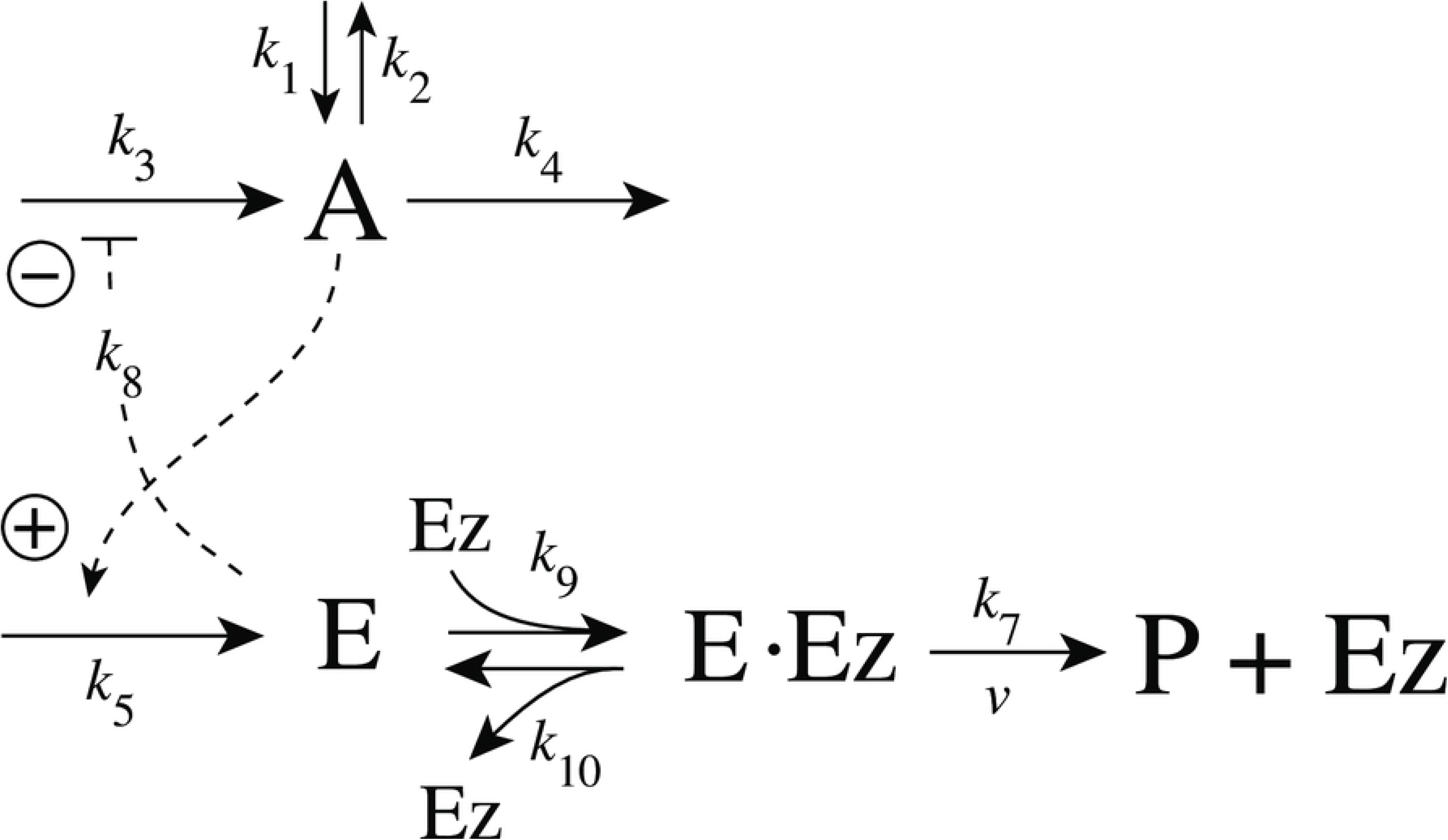
Motif 2 single-E controller: removal of E by enzyme Ez using a Michaelis-Menten mechanism.

The rate equations for the scheme in Fig 6 are:

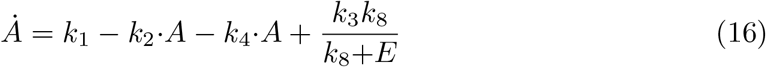

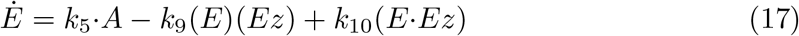

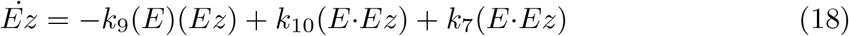

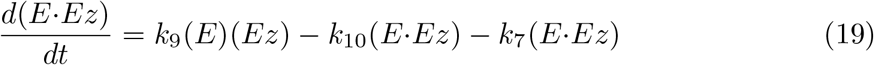

In this case the set-point of the controller is described by Eq 15.

#### The catalyzed m2-controllers: Failure at larger perturbation strengths and enzyme limitation

Fig 7 shows a comparison of the single-E controller of Fig 5 and the dual-E controller of Fig 6 for step-wise perturbations in *k*_2_. While rate constants have been more or less arbitrarily set, for comparison reasons the set-points of the controllers are both put at 2.0. Due to the two different set-point expressions for the dual-E and the single-E controllers (Eqs 14 and 15) *k*_5_ and *k*_7_ values differ slightly as indicated in the legend of Fig 7. Perturbations are applied as follows: During phase 1 (0-10 time units) A is at the controllers’ set-points (2.0) with a *k*_2_-value of 10.0. During phase 2 *k*_2_ is increased step-wise using the three perturbations: 1, *k*_2_ = 1×10^2^; 2, *k*_2_ = 1×10^3^; 3, *k*_2_ = 2×10^4^. By comparing the left panels of Fig 7 it is seen that one of the advantages of the dual-E controller is that it can maintain irs set-point even under enzymatic non-zero conditions, which means that Ez is not saturated by its substrates E_1_ and E_2_. The single-E controller, however, has problems to defend its set-point as with increasing *k*_2_ values the E·Ez complex shows increased dissociation (Fig 7, lower right panel) leading to an increasingly poorer performance and thereby increased offsets in *A* from *A_set_*.

**Fig 7.**
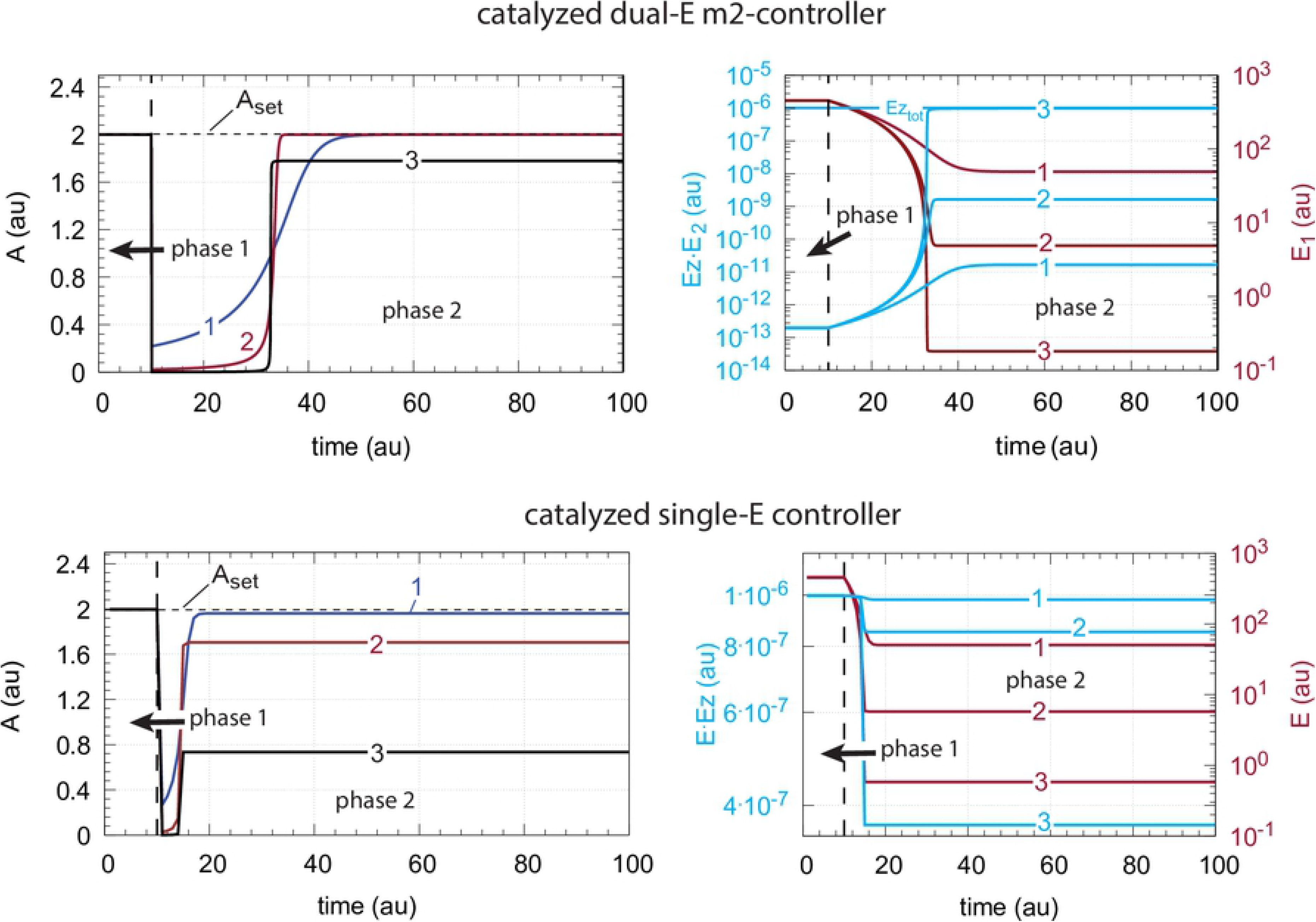
Behavior of the catalyzed dual-E controller (Fig 5) and the single-E controller (Fig 6) towards step-wise perturbations in *k*_2_. Total enzyme concentration Ez_tot_=1×10^−6^. Upper left panel: Behavior of controlled variable A of the dual-E controller. Phase 1: *k*_2_=10.0; phase 2: **1**, *k*_2_ = 1×10^2^; **2**, *k*_2_ = 1×10^3^; **3**, *k*_2_ = 2×10^4^, note the offset in A from A_set_. Upper right panel: Behavior of E_1_ and Ez·E_2_ as a function of *k*_2_-perturbations 1-3. Note that for perturbation 3 the enzyme is saturated with E_2_. Rate constants: *k*_1_=0.0, *k*_3_=1×10^5^, *k*_4_=1.0, *k*_5_=10.0, *k*_6_=20.0, *k*_7_=1×10^9^, *k*_8_=0.1, *k*_9_=1×10^8^, *k*_10_=1×10^3^, *k*_11_=1×10^8^, *k*_12_=1×10^3^, *k*_13_=1×10^8^, *k*_14_=1×10^3^, *k*_15_=1×10^8^, *k*_16_=1×10^3^. Initial concentrations: A_0_=2.0, E_1,0_=454.4, E_2,0_=0.204, Ez_0_=4.4×10^−10^, (E_1_·Ez)_0_=9.796×10^−7^, (E_1_·Ez·E_2_)_0_=2.0×10^−8^, (Ez·E_2_)_0_=1.98×10^−13^. Lower left panel: Behavior of controlled variable A for the single-E controller. Same step-wise *k*_2_ perturbations 1-3 as for the dual-E controller. Lower right panel: Behavior of E as a function of *k*_2_-perturbations. Rate constant values are the same as for the dual-E controller, except that *k*_5_=50.0, and *k*_7_=1×10^8^. Initial concentrations: A_0_=1.995, E_0_=455.5, Ez_0_=2.19×10^−9^, (Ez·E)_0_=9.976×10^−7^.

However, with perturbation 3 also the enzyme-catalyzed dual-E controller starts to break down. The reason for the breakdown is related to the total amount of enzyme, *Ez_tot_*, and the values of *k*_5_ and *k*_6_. In the present settings *k*_5_ and *k*_6_ are relatively high, which leads in the dual-E controller to a saturation of *Ez* by *E*_2_, i.e. the concentration of *EzE*_2_ approaches that of the total enzyme concentration *Ez_tot_*. Under these conditions, however, the relationship

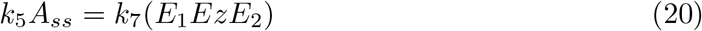

is still obeyed leading to the *A* steady state

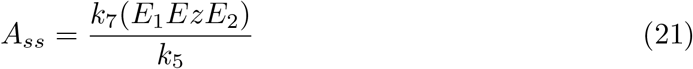

Thus, *E*_1_ still exerts control over *A* in the dual-E controller, but now in form of a *single-E (i.e. E*_1_*) control mode*. In case *Ez* works under zero-order conditions ((*E*_1_*EzE*_2_)≈*Ez_tot_*), Eq 21 becomes Eq 15. In this mode *E*_2_ shows wind-up: *E*_2_ increases linearly in time with slope *Ė*_2_ increasing with increasing *k*_2_ values. It is the increase of *E*_2_ which leads to the saturation (poisoning) of *Ez* by *E*_2_ (see Fig 7, upper right panel).

The above described limitation of the the dual-E controller can still be circumvented by either decreasing *k*_5_ and *k*_6_, or by increasing *Ez_tot_* (see next section). It should however be pointed out that there is another way of a (dual-E) controller breakdown which cannot be opposed by either increasing *Ez_tot_*) or by decreasing *k*_5_ and *k*_6_. This type of breakdown occurs when *E*_1_ is driven by *k*_2_ to such a low concentration that the compensatory flux *j_comp_* approaches its maximum value *k*_3_, i.e.

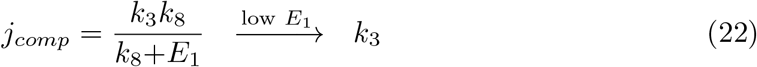

By setting in Eq 6 the term *k*_3_*k*_8_/(*k*_8_+*E*_1_) to *k*_3_ and *A* to *A_set_* we can calculate the upper limit of *k*_2_, 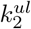,

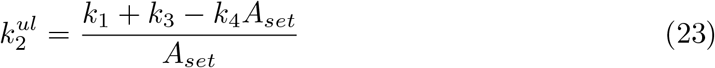

Whenever 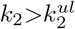 the controller breaks down irrespective of the values of *k*_5_, *k*_6_, and *Ez_tot_*. Note, that in curve 3 of the upper right panel of Fig 7 we have that

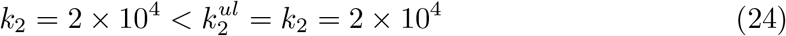

which is the reason why the dual-E’s homeostatic behavior can be restored as described in the next section.

A more detailed description of this type of breakdown is given in the chapter *Dual-E controllers based on motif 4*.

#### Avoiding enzyme limitations

Enzyme overload can be avoided by two means, either by increasing the total amount of enzyme, or by decreasing the reaction rates *k*_5_ and *k*_6_ by which *E*, *E*_1_, and *E*_2_ are formed.

Fig 8 illustrates the behavior of the controlled variable A for the antithetic controller (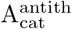, outlined in orange) in Fig 7 when perturbation 3 is applied. In panel a the total amount of enzyme has been increased from 10^−6^ to 10^−5^. In panel b the enzyme concentration is kept at 10^−6^, but *k*_5_ and *k*_6_ are in phase 2 decreased by one order of magnitude to respectively 1.0 and 2.0. In comparison, the behavior of the controlled variable A for the zero-order controller (A_zo_, outlined in black) is also shown. For the higher total enzyme concentrations both controllers behave identical, while for the decreased values of *k*_5_ and *k*_6_ the antithetic controller is less aggressive, but eventually moves A to the controller’s set-point.

**Fig 8.**
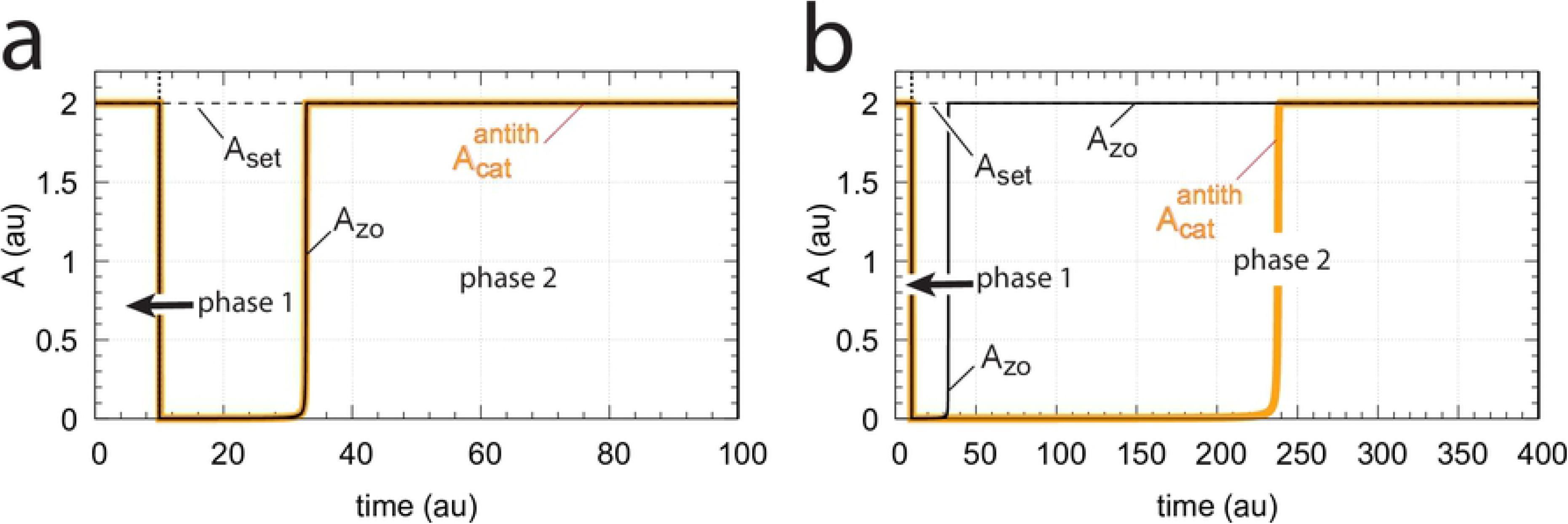
Avoiding enzyme overload. Same system as in Fig 7 with perturbation 3 applied, i.e., during phase 1 (0-10 time units) *k*_2_=10.0, while during phase 2 *k*_2_ = 2×10^4^. All other rate constants are as in Fig 7, except that in panel (a) the total amount of enzyme Ez has been increased by one order of magnitude to Ez_0_=10^−5^, while in panel (b) Ez_0_=10^−6^, but *k*_5_ and *k*_6_ have been decreased in phase 2 by one order of magnitude to 1.0 and 2.0, respectively. Initial concentrations: (a) A_0_=2.0, E_1,0_=454.4, E_2,0_=0.0204, Ez_0_=4.4×10^−10^, (E_1_·Ez)_0_=9.98×10^−6^, (E_1_·Ez·E_2_)_0_=2.0×10^−8^, (Ez·E_2_)_0_=1.98×10^−14^; (b) as in Fig 7.

#### Switching between dual-E and single-E control mode at zero-order conditions

We found that a change in the control mode of the dual-E controller (Fig 5) occurs in dependence to the relative values of *k*_5_ and *k*_6_. When *k*_6_ is lower than the rate *k*_7_Ez_tot_ the controller works in an antithetic/dual-E mode. We assume here that the dual-E controller works under zero-order conditions with large values of *k*_9_ and *k*_11_ relative to *k*_10_ and *k*_12_ (Fig 5) leading that *v* is at its maximum velocity, i.e.

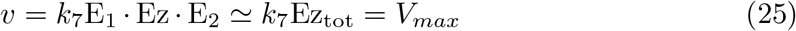

In dual-E mode both E_1_ and E_2_ participate in the regulation of A and A_set_ is given by Eq 14. However, when *k*_6_ is larger than *k*_7_Ez_tot_ the system switches to a single-E control mode where only E_1_ takes part in the regulation of A. A_set_ is now described by Eq 15. Fig 9 illustrates the behavior. Panel a shows the steady state values of A (A_ss_, gray solid circles) as a function of *k*_6_ when *k*_5_=0.4, *k*_7_=1×10^6^ and Ez_tot_=1×10^−6^. For *k*_6_ values lower than *k*_7_Ez_tot_ (=1.0) the system shows dual-E control with a set-point of *k*_6_/*k*_5_, while when *k*_6_ is larger than *k*_7_Ez_tot_=1 single-E control is observed with A_set_ being *k*_7_Ez_tot_/*k*_5_ (=2.5). In such a setting the system behaves precisely as a single-E controller (Fig 6) where E is replaced by E_1_ and Ez is replaced by Ez · E_2_. Under single-E mode conditions E_2_ does not participate in the control of A and its concentration rises continuously (showing wind-up).

**Fig 9.**
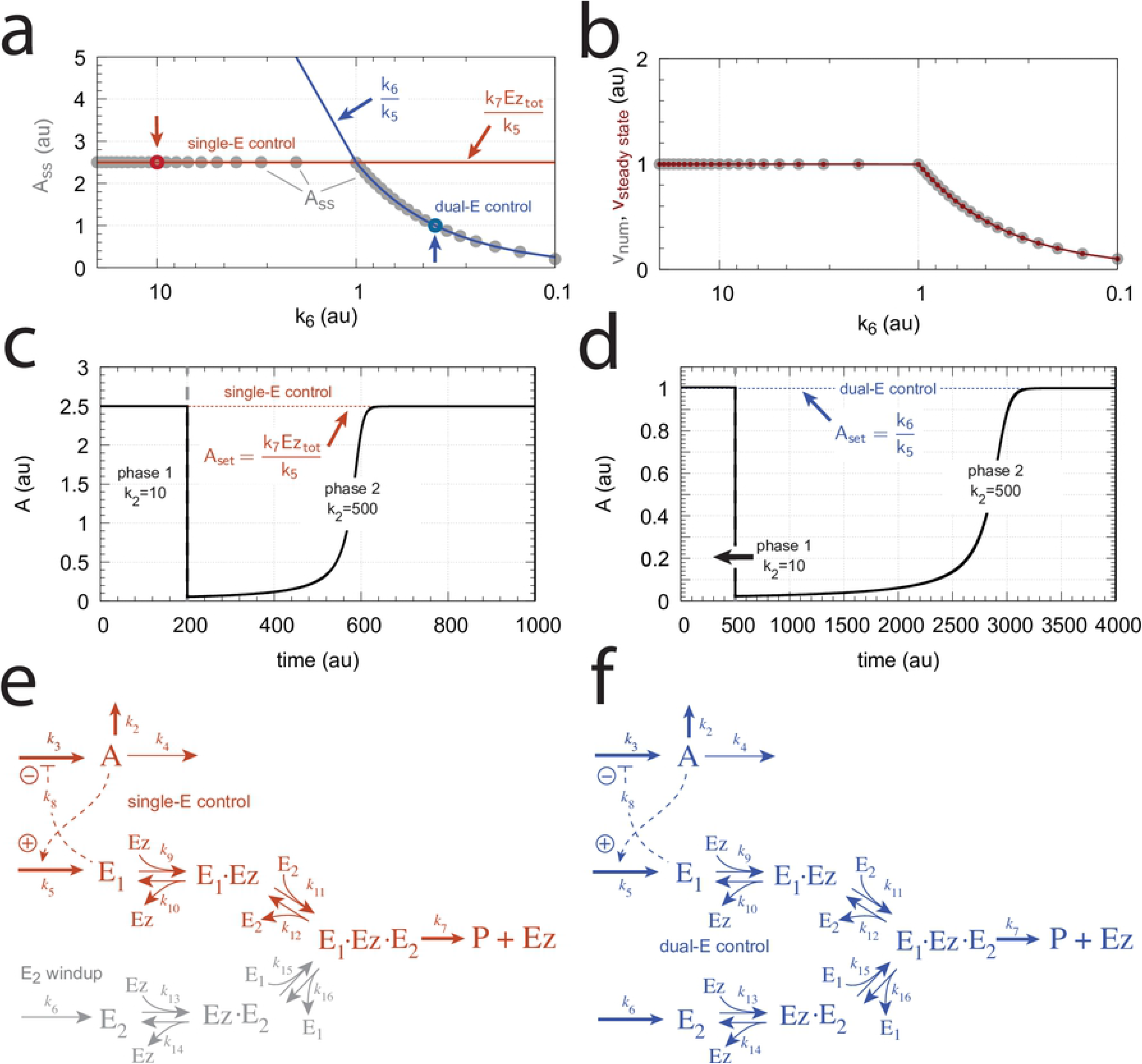
Switch between dual-E and single-E control in the motif 2 antithetic controller with a random-order ternary-complex mechanism removing E_1_ and E_2_ (Fig 5). (a) A_ss_ (steady state in A) as a function of *k*_6_. Red and blue lines indicate the respective set-point values for single-E and dual-E control. Gray solid points show the numerically calculated steady state levels. The outlined red and blue circles (indicated by the vertical arrows) show the *k*_6_ values (10.0 and 0.4) used in panels c and d when changes in *k*_2_ are applied. (b) Steady state values of *v* (Eq 13) obtained by the King-Altman method (inner red dots, S1 Text) and numerically calculated velocities (gray dots). (c) and (d) Single-E and dual-E control when *k*_6_ values are respectively 10.0 and 0.4, and *k*_2_ changes step-wise from 10.0 to 500. Other rate constants: *k*_3_=1×10^5^, *k*_4_=1.0, *k*_5_=0.4, *k*_7_=1×10^6^, *k*_8_=0.1, *k*_9_=1×10^8^, *k*_10_=1×10^3^, *k*_11_=1×10^8^, *k*_12_=1×10^3^, *k*_13_=1×10^8^, *k*_14_=1×10^3^, *k*_15_=1×10^8^, *k*_16_=1×10^3^. Initial concentrations, panel c: A_0_=2.5, E_1,0_=363.5, E_2,0_=4.5×10^4^, Ez_0_=3.04×10^−13^, (E_1_·Ez)_0_=4.3×10^−13^, (E_1_·Ez·E_2_)_0_=1.0×10^−6^, (Ez·E_2_)_0_=2.7×10^−11^. Initial concentrations, panel d: A_0_=1.0, E_1,0_=905.3, E_2,0_=6.7×10^−3^, Ez_0_=4.4×10^−12^, (E_1_·Ez)_0_=6.0×10^−7^, (E_1_·Ez·E_2_)_0_=4.0×10^−7^, (Ez·E_2_)_0_=4.5×10^−15^. (e) Outlined in red: the active part of the network during single-E control. E_2_ is continuously increasing (wind-up). (f) In dual-E control the entire network participates in the control of A (outlined in blue).

Fig 9b shows numerical and steady state values of *v* (Eq 13); they are in excellent agreement.

As illustrations, Figs 9c and d show that the set-points of single- and dual-E control are indeed defended. The two panels show the homeostatic responses when *k*_6_=10 (vertical downward red arrow in Fig 9a) and when *k*_6_=0.4 (vertical upright blue arrow in Fig 9a).

Fig 9e shows the part of the network (outlined in red) when single-E control is active. At the steady state in A, the rate *k*_5_A_set_ becomes equal to the degradation rate *v*=*k*_7_(E_1_ · Ez · E_2_)= *k*_7_ Ez_tot_. Typical for the dual-E control (Fig 9f) is that *k*_5_A_set_ and *k*_6_ are equal to *v*=*k*_7_(E_1_ · Ez · E_2_).

#### Switching between dual-E and single-E control mode at nonzero-order conditions

In this section we compare the dual-E and single-E control modes when *v*=*k*_7_(*E*_1_·*Ez*·*E*_2_) is not zero-order with respect to (*E*_1_·*Ez*·*E*_2_).

For single-E control (Fig 9e) nonzero-order conditions imply that

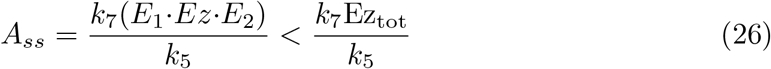

For the dual-*E* control (Fig 9f) *A_ss_* is given by Eq 14 *independent* whether the removal of the ternary complex is zero-order or not. However, dual-*E* mode will switch to single-*E* mode when

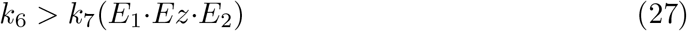

In this case *E*_2_ will show wind-up (i.e., continuously increase unless there is a removal of *E*_2_) and *A_ss_* is determined by the relationship:

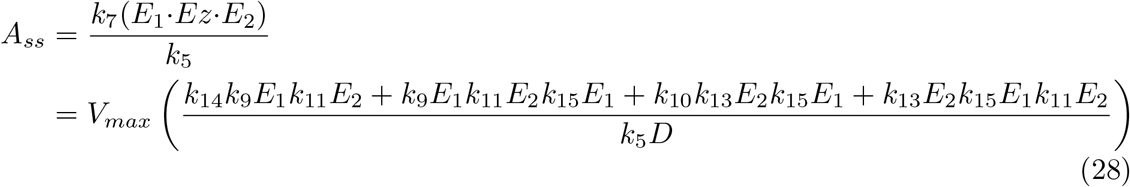

where *V_max_*=*k*_7_(*Ez_tot_*). *D* is the sum of all King-Altman numerator terms described in S1 Text.

Fig 10 illustrates the behavior going from zero-order to nonzero-order conditions. To impose nonzero-order conditions we have for the sake of simplicity, changed the values of *k*_9_, *k*_11_, *k*_13_, and *k*_15_ from 1×10^8^ (practical zero-order, panels a and b) to 1×10^6^ (panels c and d) and 1×10^4^ (panels e and f), while other rate constants are kept unchanged. In all panels single-*E* control responses are outlined in red, while dual-*E* control is outlined in blue. Fig 10 clearly shows that when the system moves into a nonzero-order kinetics regime (by lowering *k*_9_, *k*_11_, *k*_13_, and *k*_15_) the performance by single-*E* control gets successively worse. However, although dual-*E* control can maintain/defend its set-point (Eq 14) the range of the dual-*E* working mode shrinks with increasing nonzero-order kinetics (i.e., with decreasing values of *k*_9_, *k*_11_, *k*_13_, and *k*_15_).

**Fig 10.**
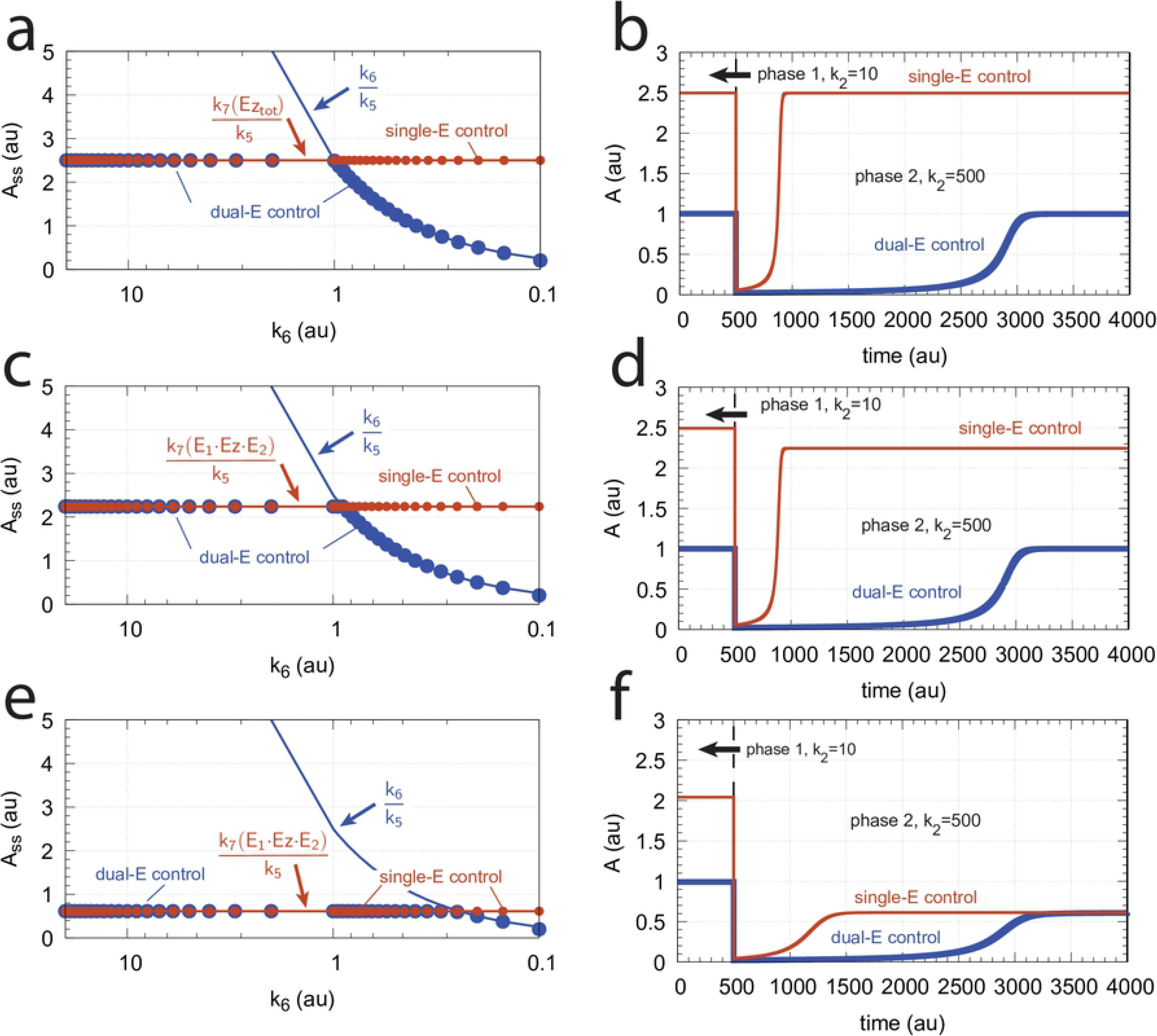
Behaviors of single-E control and dual-E control for the schemes in Fig 9e and Fig 9f when going from zero-order to nonzero-order conditions. In panels (a) and (b), *k*_9_=*k*_11_=*k*_13_=*k*_15_=1 × 10^8^ (zero-order condition); in panels (c) and (d), *k*_9_=*k*_11_=*k*_13_=*k*_15_=1 × 10^6^ (weak nonzero-order); in panels (e) and (f), *k*_9_=*k*_11_=*k*_13_=*k*_15_=1 × 10^4^ (strong nonzero-order). Panels b, d, and f to the right show the time-dependent kinetics of *A* for a step-wise perturbation in *k*_2_ from 10 (phase 1) to 500 (phase 2) applied at t=500. The *k*_6_ values in these calculations were 0.4. Other rate constants as in Fig 9. Initial concentrations for panels (a), (c), and (e), dual-*E* controller: A_0_=2.0, E_1,0_=4.5×10^2^, E_2,0_=2.0×10^−1^, Ez_0_=4.4×10^−10^, (E_1_·Ez)_0_=9.7×10^−7^, (E_1_·Ez·E_2_)_0_=2.0×10^−8^, (Ez·E_2_)_0_=2.0×10^−13^; single-*E* controller: A_0_=2.5, E_0_=3.6×10^2^, Ez_0_=2.8×10^−11^, (E·Ez)_0_=1.0×10^−8^; steady state concentrations were obtained after 2000 time units. Initial concentrations panels (b) and (d): dual-*E* controller: A_0_=1.0, E_1,0_=9.1×10^2^, E_2,0_=6.7×10^−1^, Ez_0_=4.4×10^−10^, (E_1_·Ez)_0_=6.0×10^−7^, (E_1_·Ez·E_2_)_0_=4.0×10^−7^, (Ez·E_2_)_0_=7.6×10^−13^; single-*E* controller: A_0_=2.5, E_0_=3.6×10^2^, Ez_0_=2.7×10^−9^, (E·Ez)_0_=1.0×10^−8^. Initial concentrations panels panel (f): dual-*E* controller: A_0_=1.0, E_1,0_=9.1×10^2^, E_2,0_=6.7×10^−1^, Ez_0_=4.4×10^−10^, (E_1_·Ez)_0_=6.0×10^−7^, (E_1_·Ez·E_2_)_0_=4.0×10^−7^, (Ez·E_2_)_0_=7.6×10^−13^; single-*E* controller: A_0_=2.04, E_0_=4.5×10^2^, Ez_0_=1.8×10^−7^, (E·Ez)_0_=8.1×10^−7^.

#### Motif 2 dual-E controller removing *E*_1_ and *E*_2_ by enzymes using compulsory-order ternary complex mechanisms

In the compulsory-order ternary-complex mechanisms E_1_ and E_2_ bind in an ordered manner to enzyme *Ez*, either E_1_ first (Fig 11a), or E_2_ first (Fig 11b).

**Fig 11.**
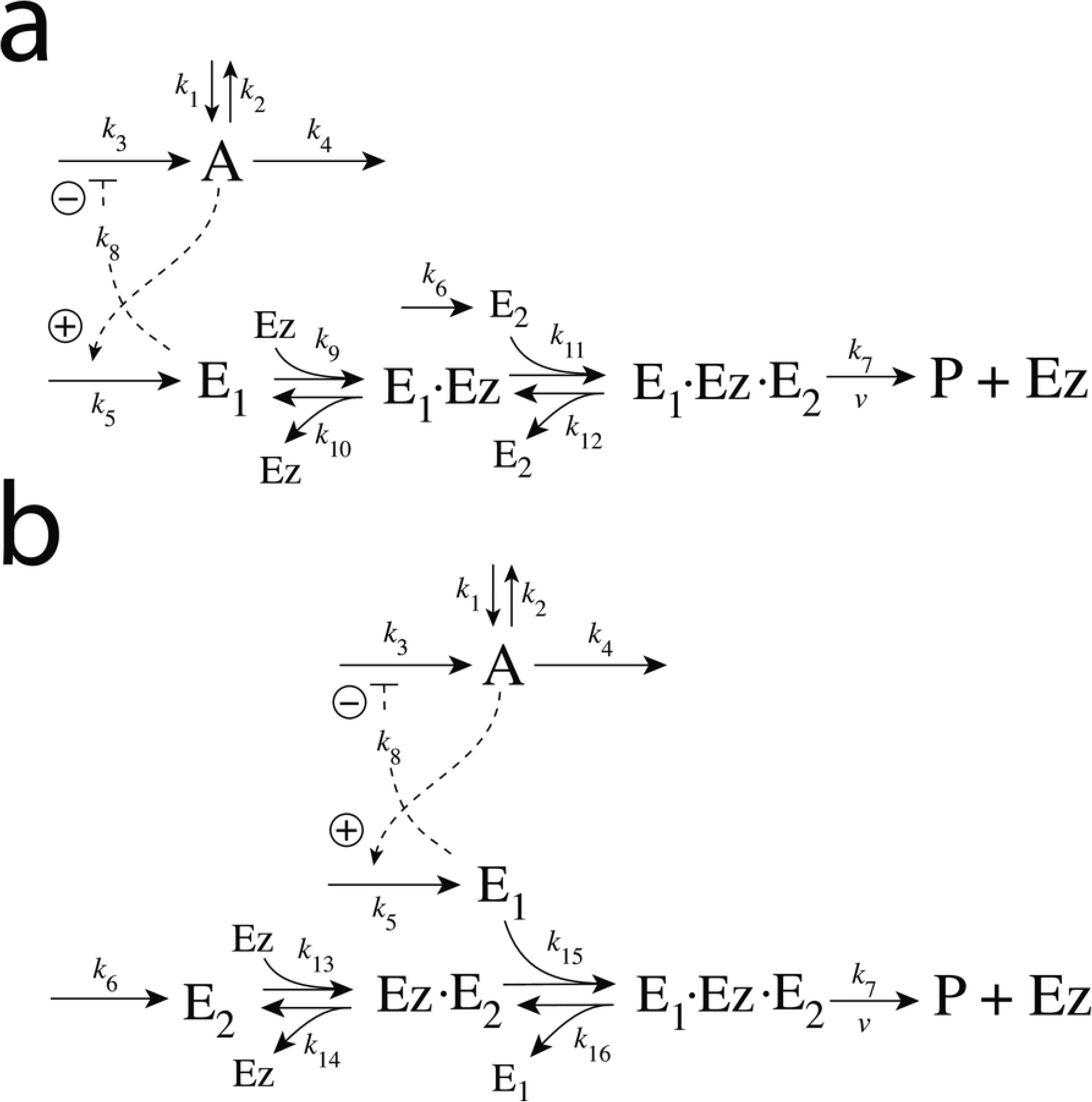
Motif 2 dual-E controller when *E*_1_ and *E*_2_ are removed enzymatically by compulsory-order ternary-complex mechanisms. Panel a: E_1_ binds first to free enzyme Ez. Panel b: E_2_ binding first to Ez.

Both mechanisms in Fig 11 can show single-E (E_1_) or dual-E control dependent on the value of *k*_6_.

We found that the mechanism when E_1_ binds first (Fig 11a) behaves analogous to the random-order ternary complex mechanism of Fig 5. Fig 12a shows the identical responses of the compulsory-order (E_1_ binds first) and the random-order ternary complex mechanisms when both controllers work in dual-E mode and both are subject to the same step-wise changes in *k*_2_ from 10.0 to 500.0. The switch of the compulsory-order (E_1_ binds first) controller (Fig 11a) from dual-E to single-E mode is shown in Fig 12b when in phase 2, besides the step-wise increase of *k*_2_, *k*_6_ is increased from 0.4 to 10.0.

**Fig 12.**
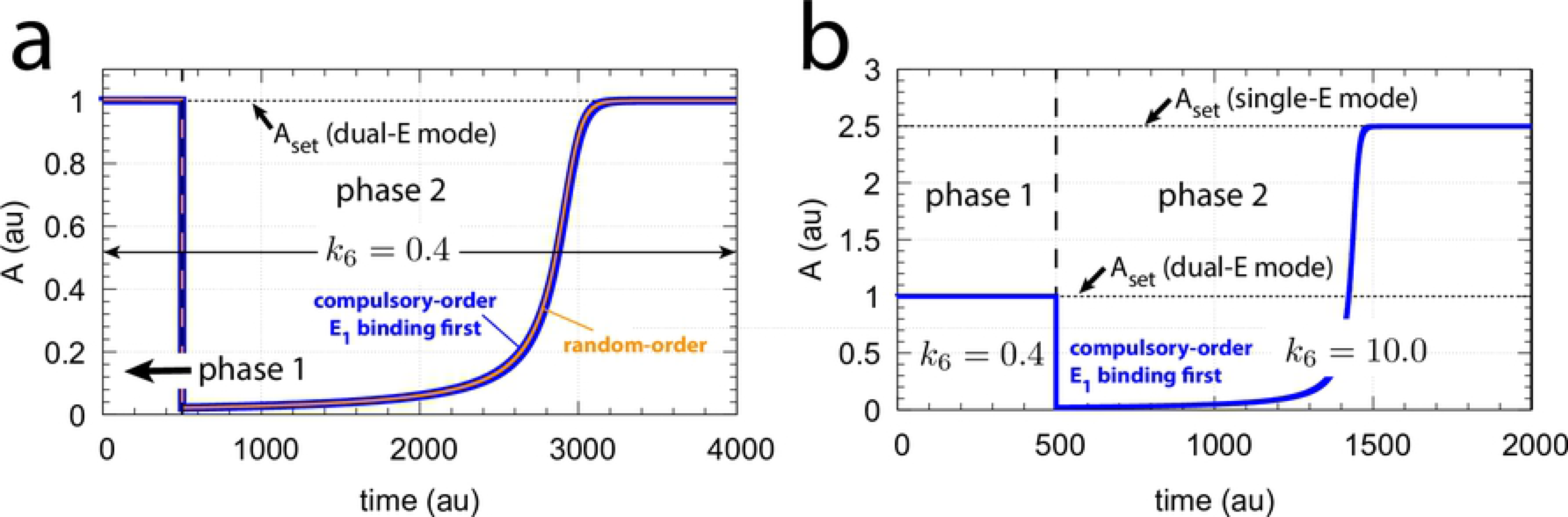
Dual- and single-E control mode of the m2 feedback loop when E_1_ and E_2_ are removed by a compulsory-order ternary complex mechanism and when E_1_ binds first to Ez (Fig 11a). Panel a, outlined in blue, shows the concentration of *A* for the mechanism of Fig 11a with a step-wise change of *k*_2_ from 10.0 (phase 1) to 500.0 (phase 2). For comparison, outlined in orange, the results of Fig 10b for the random-order ternary complex mechanism working in dual-E mode are shown. Rate constant *k*_6_=0.4 for both phases. Other rate constants and initial concentrations are the same as for Fig 10b. Panel b shows the concentration of *A* for the compulsory-order ternary complex mechanism from panel a, but *k*_6_ is changed in phase 2 from 0.4 to 10.0. The controller switches in phase 2 from dual-E mode to single-E mode with the associated change of *A_set_* from 1.0 (Eq 14) to 2.5 (Eq 15). Initial concentrations and rate constants as in panel a.

Fig 13 shows the single-E and dual-E control mode when E_1_ and E_2_ are removed by a compulsory-order ternary complex mechanism, but E_2_ binds first to Ez (Fig 11b). Panel a shows *A_ss_* as a function of *k*_6_ while panel b shows the numerical and the King-Altman steady state values of the degradation rate *v* of the ternary complex (S1 Text). In single-E mode the controller of Fig 11b behaves precisely as the single-E controller of Fig 6.

**Fig 13.**
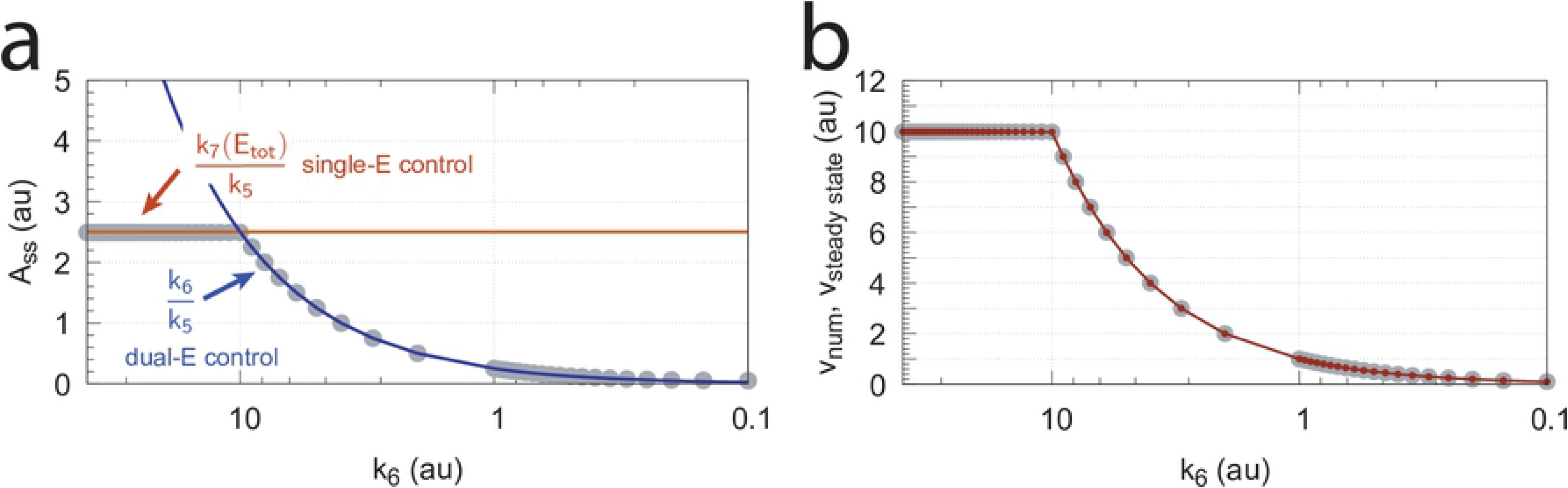
Switch between single-E and dual-E control for the m2 controller when E_1_ and E_2_ are removed by a compulsory-order ternary-complex mechanism with E_2_ binding first to Ez (Fig 11b). Panel a: steady state values of *A* (*A_ss_*) as a function of *k*_6_. Gray dots show numerical results. The line outlined in red describes the set-point of *A* (*k*_7_*Ez_tot_/k*_5_) at single-E control. The blue line shows the set-point of *A* (*k*_6_/*k*_5_) when the system is in dual-E control mode. Panel b: corresponding numerical (gray dots) and steady state values (red small dots, calculated by King-Altman method, S1 Text) of the degradation rate *v* of the ternary-complex (Eq 13). Rate constants: *k*_1_=0.0, *k*_2_=100.0, *k*_3_=1×10^5^, *k*_4_=1.0, *k*_5_=4.0, *k*_6_ varies between 40.0 and 0.05, *k*_7_=1×10^7^, *k*_8_=0.1, *k*_13_=*k*_15_=1×10^8^, *k*_14_=*k*_16_=1×10^3^. Initial concentrations: A_0_=1.0, E_1,0_=9.1×10^2^, E_2,0_=6.7×10^−2^, Ez_0_=6.0×10^−7^, (E_1_·Ez·E_2_)_0_=4.0×10^−7^, (Ez·E_2_)_0_=4.4×10^−11^. Ez_tot_=1.0×10^−6^. Steady state values were obtained after 10000 time units.

Also for this compulsory-order ternary-complex mechanism (Fig 11b) single-E control is observed when *k*_6_ is getting larger than *k*_7_(E_1_ · Ez · E_2_) or, as in Fig 13, *k*_6_ is larger than *k*_7_(Ez_tot_) in the case *v* is zero-order with respect to (E_1_ · Ez · E_2_). When *k*_6_ is smaller than *k*_7_(E_1_ · Ez · E_2_) (or *k*_7_(Ez_tot_)) the controller of Fig 11b will work in dual-E mode.

#### Critical slowing down at spontaneous single-E to dual-E mode transitions

We have seen above that when E_1_ and E_2_ are removed by an enzymatic ternary-complex mechanism then, dependent on *k*_6_, the m2-controller can work either in a single-E or in a dual-E mode, where each of the control modes can have separate set-points. However, even when the condition for dual-E control mode is fulfilled, i.e. when

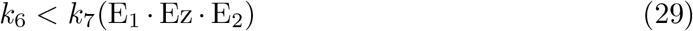

the system can still stay in single-E mode whenever *E*_2_ is kept at a high value. In this situation the single-E control mode is “*metastable*”, i.e., *A* will be kept at the set-point of the single-E control mode until *E*_2_ has reached its steady state. Then *A* changes abruptly to the set-point of the dual-E controller. This “metastability” of the single-E control mode, with the condition of Eq 29 fulfilled, is illustrated in Fig 14a with two values of *k*_6_. For this purpose we have chosen the controller described by Fig 11b, but the other mechanism (Fig 11a) also shows this phenomenon. Outlined in red are the traces of *A*, while blue lines indicate the concentrations of *E*_2_. Continuous lines have a *k*_6_ of 4.0 while the dotted lines relate to a *k*_6_ value of 8.0. Calculations start with a high initial values of *E*_2_ (see legend of Fig 14). While *E*_2_ gradually decreases *A* remains at the set-point of the single-E control mode until it ubruptly changes to the set-point of the dual-E control mode. Also note that even when the single-E controller is metastable, it can still defend its set-point (see the m5 motif below for an explicit example).

**Fig 14.**
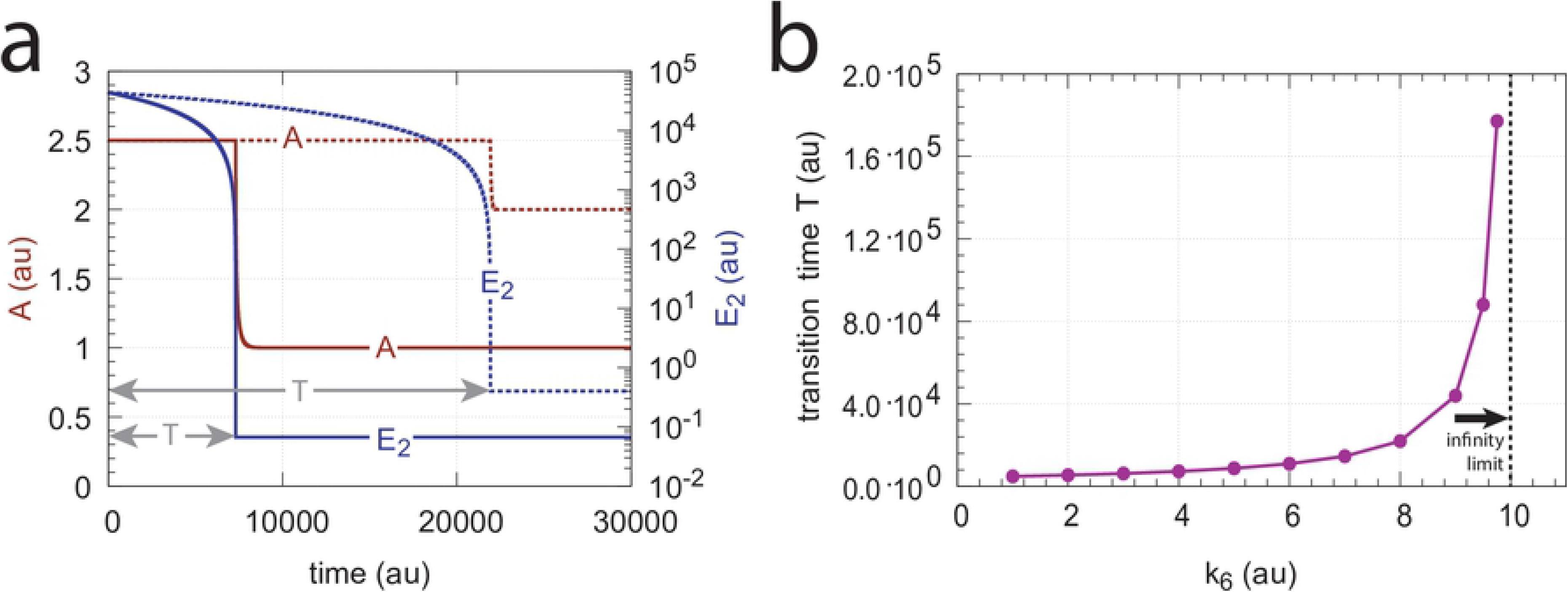
Critical slowing down in the transition from single-E to dual-E control in the negative feedback loop of Fig 11b. The set-point of A during single-E control is 2.5, but 1.0 during dual-E control. Panel a: Time profiles of *A* and *E*_2_ for *k*_6_=4.0 (solid lines) and *k*_6_=8.0 (dotted lines). T, the transition time, is the time difference from t=0 until *E*_2_ has reached steady state. Panel b: T as a function of *k*_6_. When *k*_6_ →10.0 the steady state of the dual-E control mode vanishes and T→ ∞. Rate constants (for each data point): *k*_1_=0.0, *k*_2_=10.0, *k*_3_=1×10^5^, *k*_4_=1.0, *k*_5_=4.0, *k*_6_ takes the values 1.0, 2.0, …, 9.0, 9.5 and 9.75, *k*_7_=1×10^7^, *k*_8_=0.1, *k*_13_=*k*_15_=1×10^8^, *k*_14_=*k*_16_=1×10^3^. Initial concentrations: A_0_=2.5, E_1,0_=3.635×10^2^, E_2,0_=4.38×10^4^, Ez_0_=2.28×10^−13^, (E_1_·Ez·E_2_)_0_=1.0×10^−6^, (Ez·E_2_)_0_=2.75×10^−11^.

The *transition time* T (Fig 14a) denotes the time span *A* is kept at the set-point of the single-E controller until its transition to dual-E control. With increasing *k*_6_ the system shows the behavior of critical slowing down [25], i.e. T increases and approaches infinity when

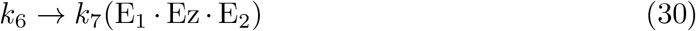

and the set-point for the dual-E control mode vanishes (Fig 14b).

#### Ping-pong mechanism: Influence of total enzyme concentration on single-E and dual-E control mode

In this section we turn, for completeness, to the ping-pong type of mechanisms (Fig 15). However, we should mention that no significant differences between the behaviors of ternary-complex mechanisms and ping-pong mechanisms have been observed. Although we could have used one of the ternary-complex mechanisms to illustrate how total enzyme concentration influences m2-controller dynamics and the transitions between single-E and dual-E control modes, we use here the ping-pong mechanism of Fig 15a. While in ternary-complex mechanisms E_1_ and E_2_ need both to bind to enzyme Ez to undergo catalysis, in ping-pong mechanisms one of the substrates (E_1_ or E_2_) binds first and creates an alternative enzyme form Ez* after forming a first product (for the sake of simplicity we have omitted it). Then Ez* can bind the second substrate which leads to the final product, and regenerates Ez (Fig 4a,b). The two mechanisms in Fig 15 differ in the binding order of E_1_ and E_2_. When E_1_ binds first to Ez (Fig 15a) the rate equations become:

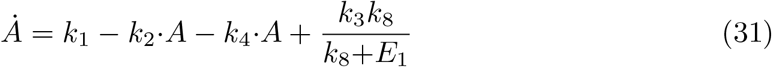

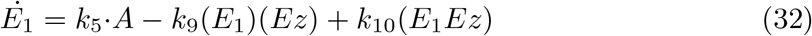

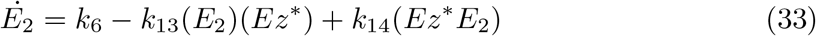

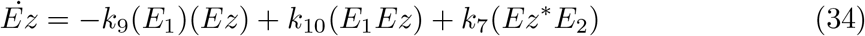

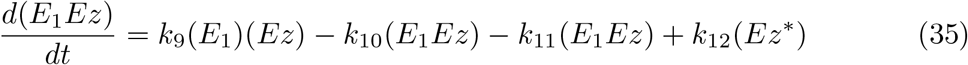

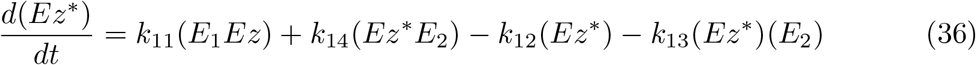

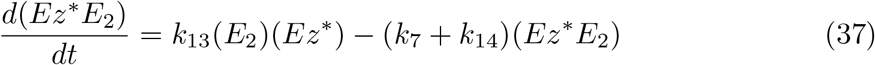

**Fig 15.**
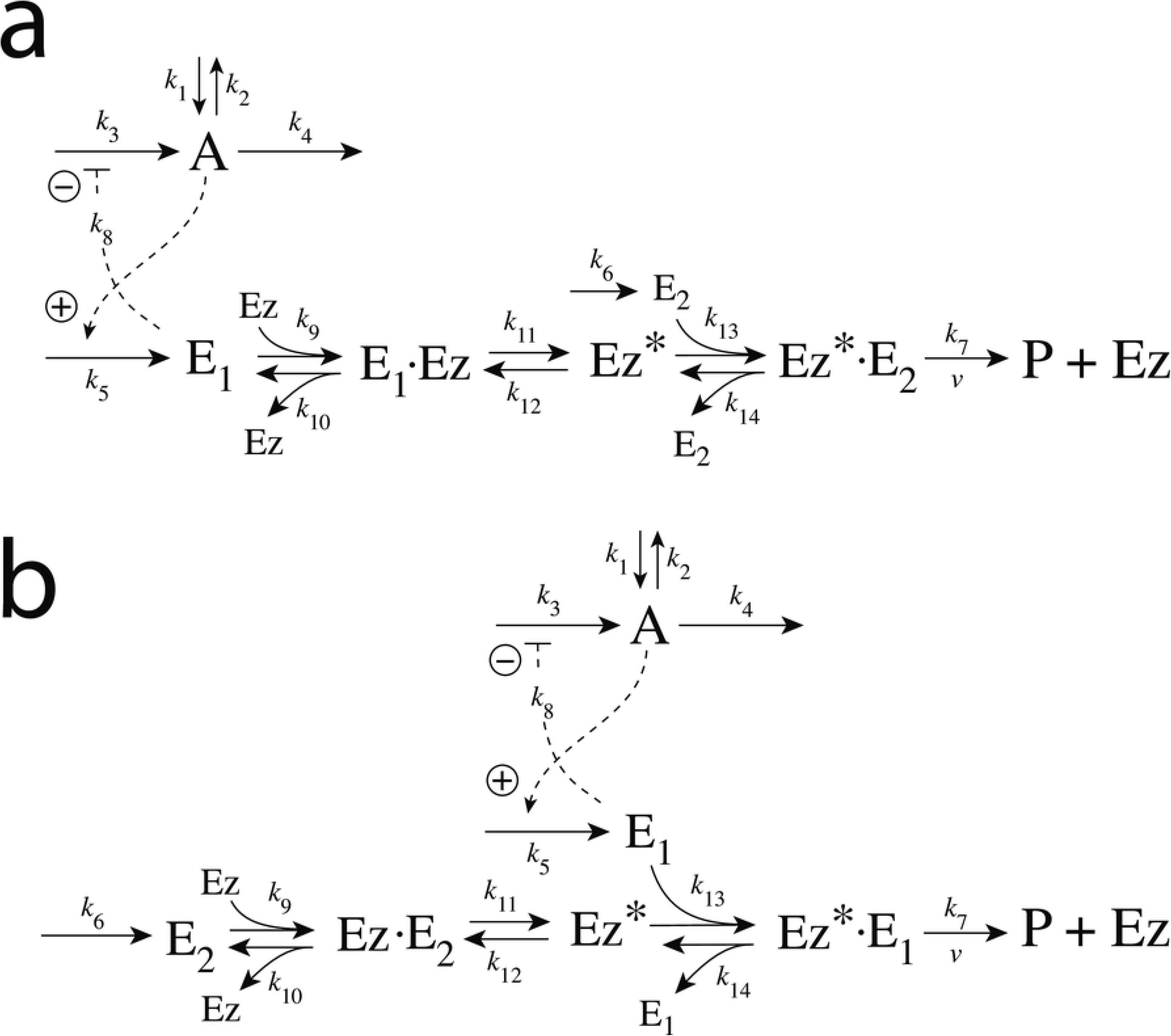
Enzymatic ping-pong mechanisms removing E_1_ and E_2_ in m2 dual-E controller. (a) E_1_ binds first to Ez. (b) E_2_ binds first to Ez.

Fig 16 shows the effect of total enzyme concentration (Ez_tot_) when in Fig 15a *k*_9_, *k*_11_, and *k*_13_ values are such high that the removal rate of E_1_ and E_2_, given by

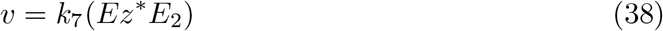

becomes zero order with respect to E_1_ and E_2_, i.e., *v* ≈ *V_max_*=*k*_7_*Ez_tot_*.

**Fig 16.**
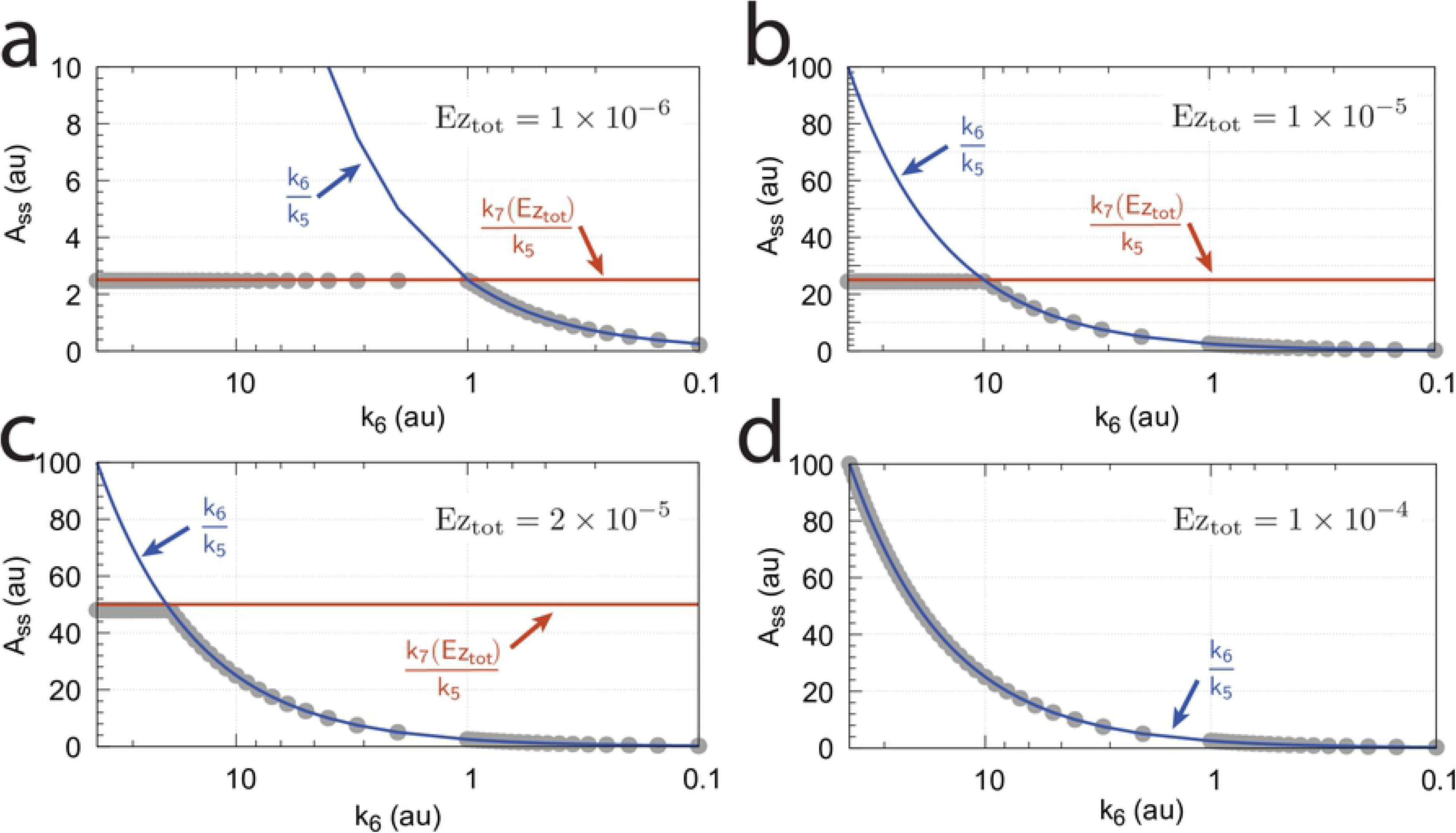
Influence of total enzyme concentration Ez_tot_ on the switch between dual-E and single-E control in the m2 controller with ping-pong mechanism of Fig 15a. (a) Ez_tot_=1×10^−6^, (b) Ez_tot_=1×10^−5^, (c) Ez_tot_=2×10^−5^, (d) Ez_tot_=1×10^−4^. Rate constants: *k*_1_=0.0, *k*_2_=500.0, *k*_3_=1×10^5^, *k*_4_=1.0, *k*_5_=0.4, *k*_6_ varies between 40.0 and 0.05, *k*_7_=1×10^6^, *k*_8_=0.1, *k*_9_=*k*_11_=*k*_13_=1×10^8^, *k*_10_=*k*_12_=*k*_14_=1×10^3^. Initial concentrations: A_0_=2.0, E_1,0_=4.5×10^2^, E_2,0_=2.0×10^−1^, Ez_0_=Ez_tot_, (E_1_·Ez)_0_=0.0, (Ez^*^)_0_=0.0, (Ez^*^E_2_)_0_=0.0. Steady state values were obtained after 4000 time units.

Panels a-d of Fig 16 show the steady state of A (A_ss_, gray dots) as a function of *k*_6_ when Ez_tot_ increases from 1 × 10^−6^ (panel a) up to 1 × 10^−4^. One sees clearly the increase of the operational range for the dual-E control mode to higher *k*_6_ values, while the set-point corresponding to the single-E control mode increases with increasing Ez_tot_ concentration.

#### Ping-pong mechanism: Influence of nonzero-order conditions on single-E and dual-E control mode

In this section we show how nonzero-order conditions of *v*=*k*_7_(Ez*E_1_) with respect to E_1_ and E_2_ influence the ping-pong mechanism. For this purpose we show the results for the mechanism of Fig 15b when E_2_ binds first to Ez.

The rate equations are:

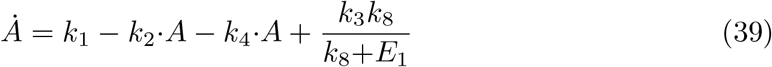

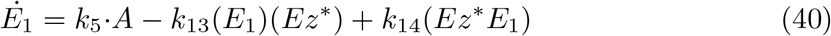

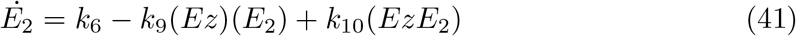

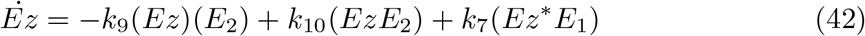

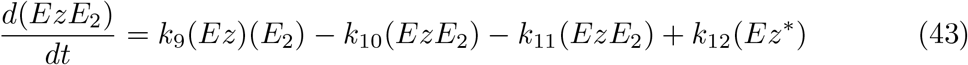

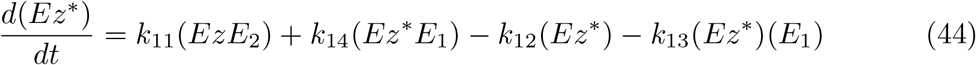

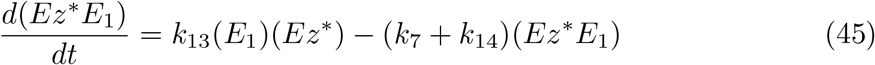

Fig 17 shows the switching behavior from dual-E control, gray dots on blue lines) to single-E control (horizontal gray dots) with changing *k*_6_ as a function of the rate constants *k*_9_, *k*_11_, and *k*_13_. The red lines indicate the steady state of A when single-E control mode works under zero-order conditions, i.e. at high values of *k*_9_, *k*_11_, and *k*_13_. In this case we have that

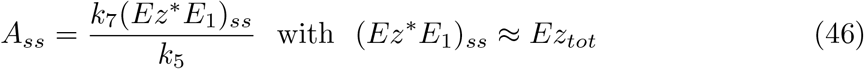

**Fig 17.**
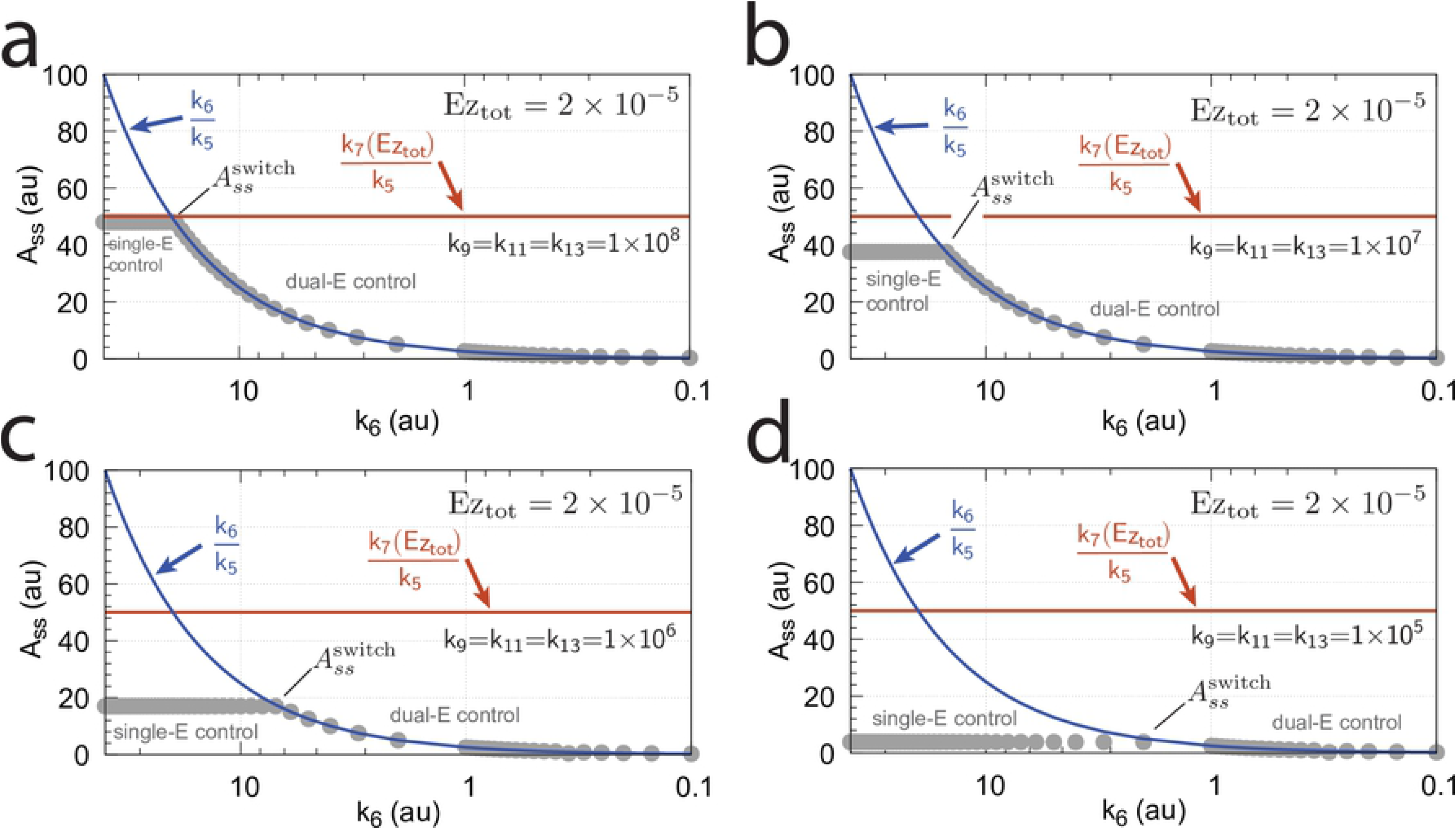
Change of the switch point between dual-E and single-E control with decreasing values of *k*_9_, *k*_11_, and *k*_13_. (a) *k*_9_=*k*_11_=*k*_13_=1×10^8^; (b) *k*_9_=*k*_11_=*k*_13_=1×10^7^; (c) *k*_9_=*k*_11_=*k*_13_=1×10^6^; (d) *k*_9_=*k*_11_=*k*_13_=1×10^5^. Other rate constants: *k*_1_=0.0, *k*_2_=500.0, *k*_3_=1×10^5^, *k*_4_=1.0, *k*_5_=0.4, *k*_6_ takes values between 0.1 and 40.0 (indicated by the gray dots), *k*_7_=1×10^6^, and *k*_8_=0.1. Initial concentrations: A_0_=2.0, E_1,0_=4.5×10^2^, E_2,0_=2.0×10^−1^, Ez_0_=Ez_tot_=2.0×10^−5^, (EzE_2_)_0_=0.0, (Ez^*^)_0_=0.0, (Ez^*^E_1_)_0_=0.0. Steady state values were obtained after 4000 time units.

In the calculations of Fig 17 the total enzyme concentration *Ez_tot_* is 2 × 10^−5^. With decreasing values of *k*_9_, *k*_11_, and *k*_13_ (from panel a to d), the system moves towards nonzero-order kinetics (with respect to E_1_ and E_2_) and the steady state value of (*Ez*^*^*E*_1_) decreases. The switch-point in *A_ss_* 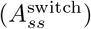 from dual-E control to single-E control occurs now at lower *A_ss_* values, described by the equation

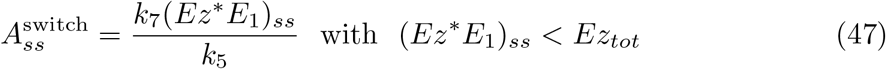

showing that nonzero-order conditions diminish the operational range of dual-E control.

Also increased values of the perturbation *k*_2_ reduces the operational range and moves 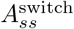 to lower values (Fig 18).

**Fig 18.**
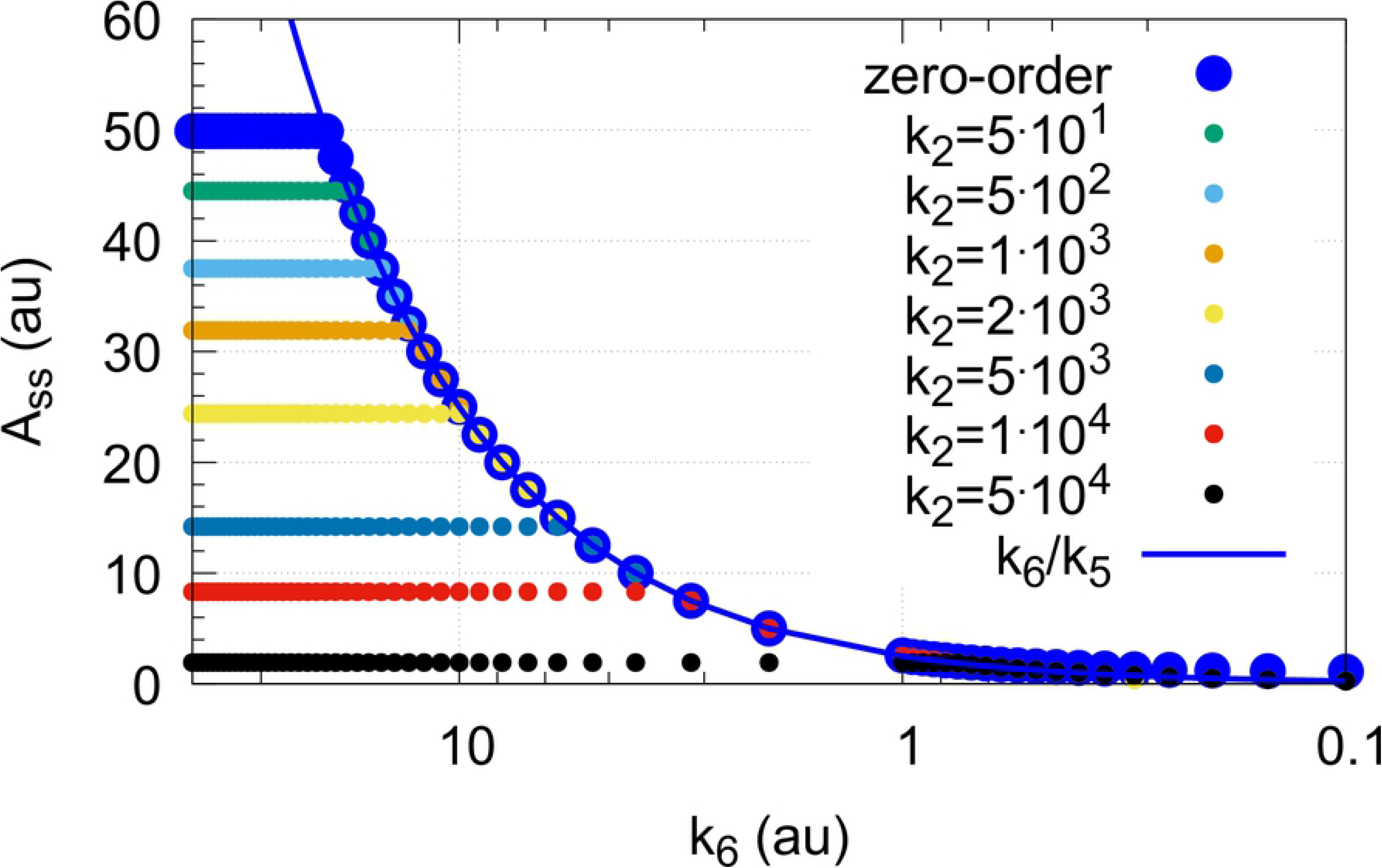
Influence of step-wise *k*_2_ for catalyzed m2 controller under nonzero-order conditions. The mechanism considered in that of Fig 15b. Small colored dots indicate *A_ss_* levels for different *k*_2_ values when *k*_9_=*k*_11_=*k*_13_=1×10^7^ and Ez_tot_=2.0×10^−5^. For comparison, large blue dots show the *A_ss_* values under zero-order conditions when *k*_9_=*k*_11_=*k*_13_=1×10^9^ and *k*_2_=1.0. Other rate constant values and initial concentrations are as in Fig 17.

#### Summary of the catalyzed m2 controllers

The catalyzed m2 controller works for all the four basic enzymatic mechanisms shown in Fig 4. Zero-order conditions for *v* (=*dP/dt*) with respect to *E*_1_ and *E*_2_ provide optimum controller performance, which, however, becomes limited at low enzyme concentrations and high perturbation (*k*_2_) values. Catalyzed antithetic controllers (i.e. controllers working in dual-E mode) become more aggressive by increased turnover numbers (*k*_7_ values). Switch to single-E control mode is observed when the rate forming *E*_2_ by *k*_6_ exceeds the degradation rate of the controller species *E*_1_ and *E*_2_. For nonzero-order conditions *A_ss_* in single-E control mode decreases with increasing *k*_2_ values. While this is also true for the dual-E control mode, in dual-E mode *A_ss_* is still determined by the ratio *k*_6_/*k*_5_ and thereby, unlike a single-E controller, shows robust control even for nonzero-order conditions.

### Controllers based on motif 4

Motif 4 is based on double inhibition. In the antithetic/dual-E setting (Fig 3), *A* is inhibiting the synthesis of *E*_1_, while *E*_2_ is now activating the compensatory flux by derepression.

#### Motif 4 dual-E controller removing *E*_1_ and *E*_2_ by a random-order ternary-complex mechanism

Fig 19 shows the dual-E m4-controller removing *E*_1_ and *E*_2_ by a random-order ternary-complex mechanism. It is, like the corresponding m2-controller, also an inflow type of controller, where the compensatory flux, 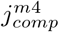 is based on derepression, now by *E*_2_, i.e

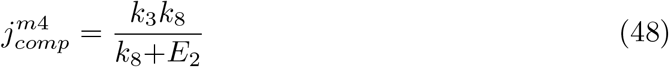

**Fig 19.**
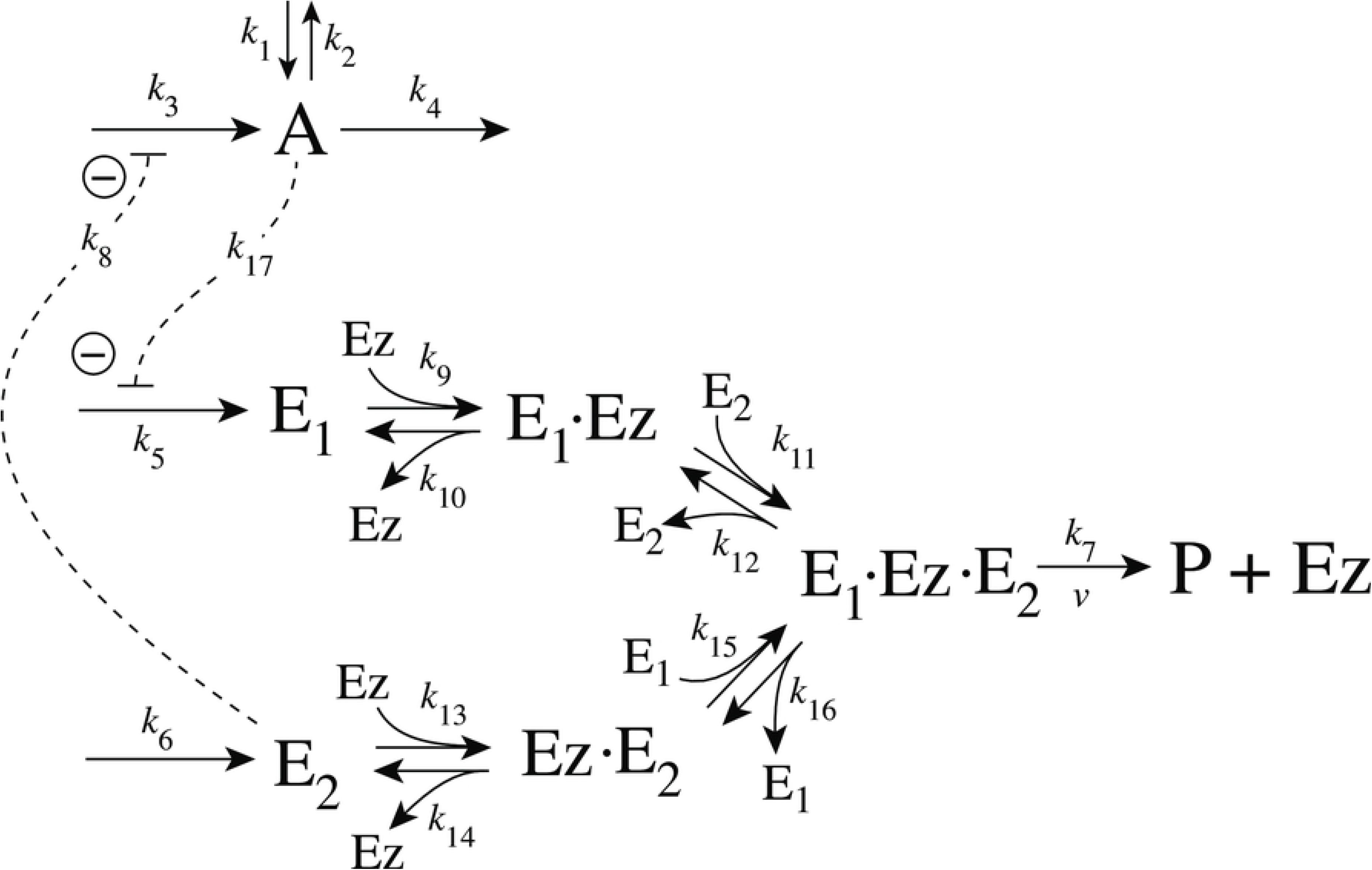
Motif 4 dual-E/antithetic controller using an enzymatic random-order ternary-complex mechanism for the removal of *E*_1_ and *E*_2_.

The rate equations for the m4-controller are:

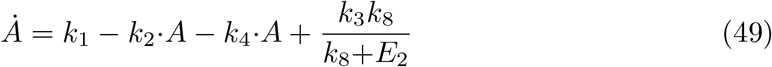

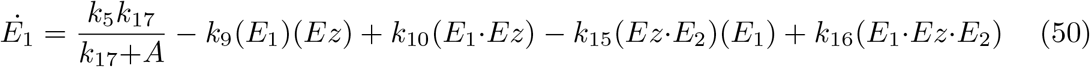

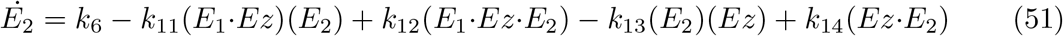

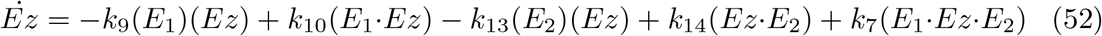

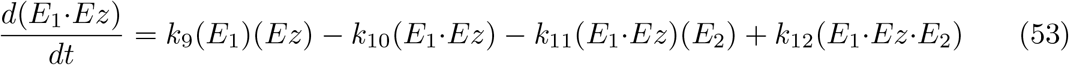

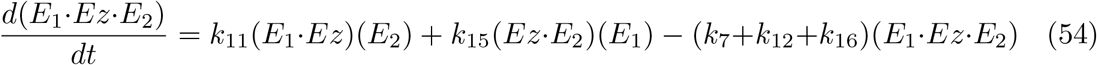

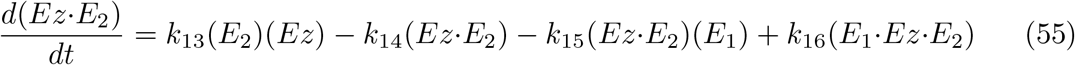

When the controller works in dual-E mode, its set-point is calculated from the following relationship

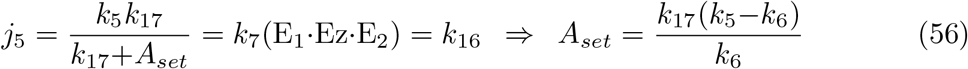

In comparison with the corresponding m2 dual-E controller (Fig 7) also for the m4 feedback arrangement the response time decreases with increased levels of step-wise perturbations in *k*_2_. Fig 20 shows the controller’s homeostatic behavior upon step-wise perturbations in *k*_2_ (curves 1-7) applied at time *t* = 50 from *k*_2_=10 (phase 1) up to *k*_2_=2×10^4^ (curve 7, phase 2).

**Fig 20.**
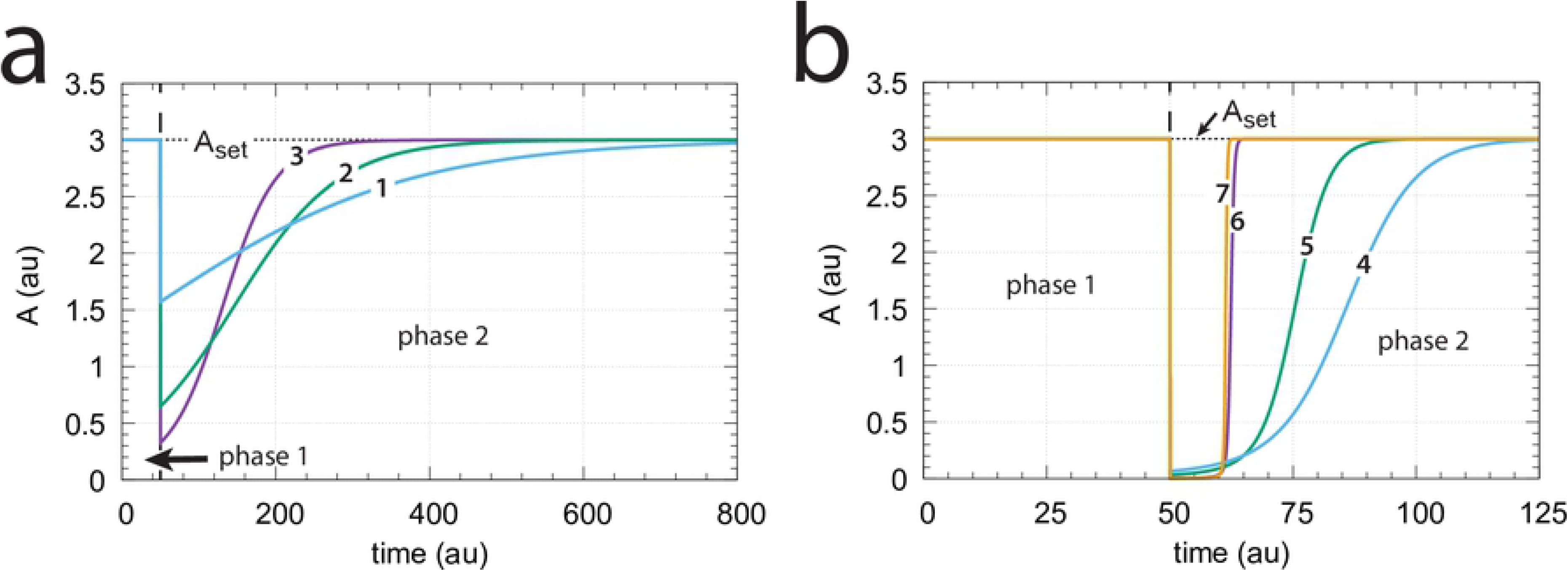
Response of the m4 random-order ternary-complex controller (Fig 19) with respect to step-wise changes in *k*_2_. (a) Phase 1: *k*_2_=10. At time *t* = 50 phase 2 starts with the following changes in *k*_2_: (1) *k*_2_=20, (2) *k*_2_=50, (3) *k*_2_=100. (b) Phase 1: *k*_2_=10. At time *t* = 50 phase 2 starts with the following changes in *k*_2_: (4) *k*_2_=500, (5) *k*_2_=1×10^3^, (6) *k*_2_=1×10^4^, (7) *k*_2_=2×10^4^. Other rate constants: *k*_1_=0.0, *k*_3_=1×10^5^, *k*_4_=1.0, *k*_5_=31.0, *k*_6_=1.0, *k*_7_=1×10^8^, *k*_8_=0.1, *k*_9_=*k*_11_=*k*_13_=*k*_15_=1×10^8^, *k*_10_=*k*_12_=*k*_14_=*k*_16_=1×10^3^, *k*_17_=0.1. Initial concentrations: A_0_=3.0, E_1,0_=1.0×10^−2^, E_2,0_=3.0×10^2^, Ez_0_=3.3×10^−11^, (E_1_·Ez)_0_=1.4×10^−15^, (E_1_·Ez·E_2_)_0_=1.0×10^−8^, (EzE_2_)_0_=9.9×10^−7^. Total enzyme concentration Ez_tot_=1.0×10^−6^.

For a given set-point *A_set_* the steady state condition of Eq 49 determines the range of *k*_2_ perturbations the controller can defend. By setting in Eq 49 *E*_2_=0 and *A*=*A_set_* the upper limit of *k*_2_, 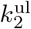, can be determined, i.e.,

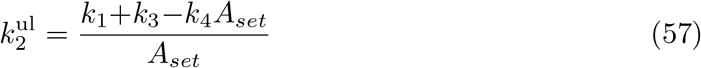

For 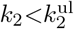 the m4 controller will defend the set-point described by Eq 56, i.e., *A_ss_*=*A_set_*. This is indicated by the the blue area in Fig 21a. The red area in Fig 21 shows the *k*_2_ values when 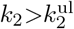 for a given set-point *A_set_*. In this case *A_ss_<A_set_* and an offset in *A* concentration from *A_set_* will be observed. Fig 21b illustrates this. During phase 1 (time between 0 and 50) *k*_2_=10.0 and the value of *A* is at its set-point *A_set_*=3.0. At time *t* = 50.0 (indicated by the blue downward arrow 1) *k*_2_ is increased to 2×10^4^. The controller is able to defend the perturbation and is still within the blue area as indicated in Fig 21a by point 1. At time *t* = 250.0 phase 3 starts with a *k*_2_ of 5×10^4^ (red downward arrow 2). Now 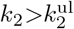 and the controller shows an offset in the controlled variable, i.e. the steady state value of *A* is below *A_set_*. With increasing *k*_2_ values the offset will increase.

**Fig 21.**
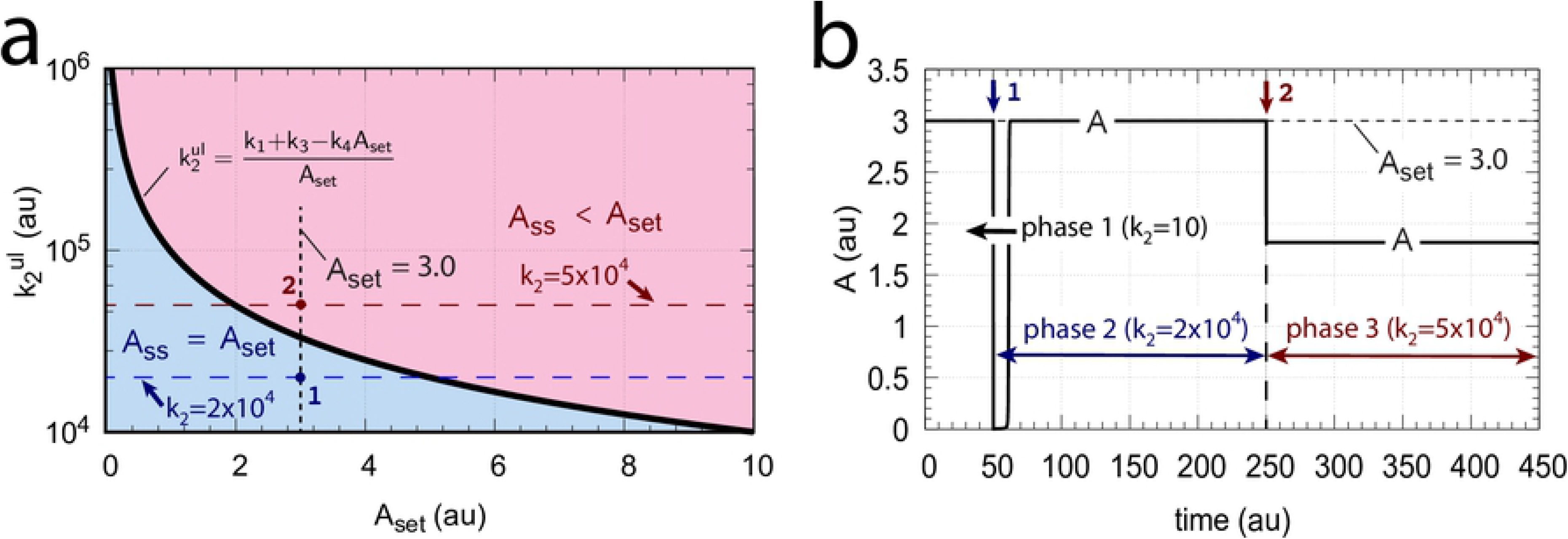
Operational range of m4 controller with upper defendable limit of *k*_2_ 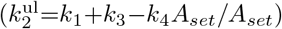. (a) Blue area indicates the 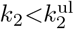 range as a function of *A_set_* in which the controller can defend *A_set_*. Black solid curve: 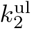 as a function of *A_set_* when *k*_1_=0.0, *k*_3_=1×10^5^, and *k*_4_=1.0. Red area: 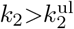 where *A_ss_* is lower than *A_set_*. (b) Computation showing the partial loss of homeostasis when *k*_2_ becomes larger than 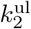. Phase 1 (0-50 time units): *k*_2_=10; phase 2 (50-250 time units): *k*_2_=2×10^4^; phase 3 (250-450 time units): *k*_2_=5×10^4^. Other rate constants and initial conditions as in Fig 20. For further descriptions, see text.

Another influence on the operational range of the m4 controller is the reaction-order by which the enzyme Ez removes *E*_1_ and *E*_2_. The reaction order is closely related to the ratios of *k*_10_/*k*_9_, *k*_12_/*k*_11_, *k*_14_/*k*_13_, and *k*_16_/*k*_15_. The ratios can be interpreted as *K_M_* values (in a rapid-equilibrium approach). For example, in the single-E m2 controller (Fig 6), an offset from *A_set_*=*k*_7_(*Ez_tot_*)/*k*_5_ (Eq 15) is observed when *k*_10_/*k*_9_ is relatively large, i.e. not small enough for the degradation of *E* to become zero-order (see Ref [4, 6] for more details). For the m4 controller (Fig 19) increasing values of the ratios *k*_10_/*k*_9_, *k*_12_/*k*_11_, *k*_14_/*k*_13_, and *k*_16_/*k*_15_ will lead to a reduction of the controller’s operational range. Fig 22 illustrates this. For the sake of simplicity, all odd-numbered rate constants *k*_9_,…*k*_15_ and all equal-numbered rate constants *k*_10_,…*k*_16_ have among themselves the same values, respectively. The turquoise areas in Fig 22 show the fully functional range of the controller as a function of *k*_5_, i.e. when the condition of Eq 56 is fulfilled and the controller works in dual-E mode. With increasing values of (*k*_10_/*k*_9_)=(*k*_12_/*k*_11_)=(*k*_14_/*k*_13_)=(*k*_16_/*k*_15_) the operational range of the controller is clearly reduced.

**Fig 22.**
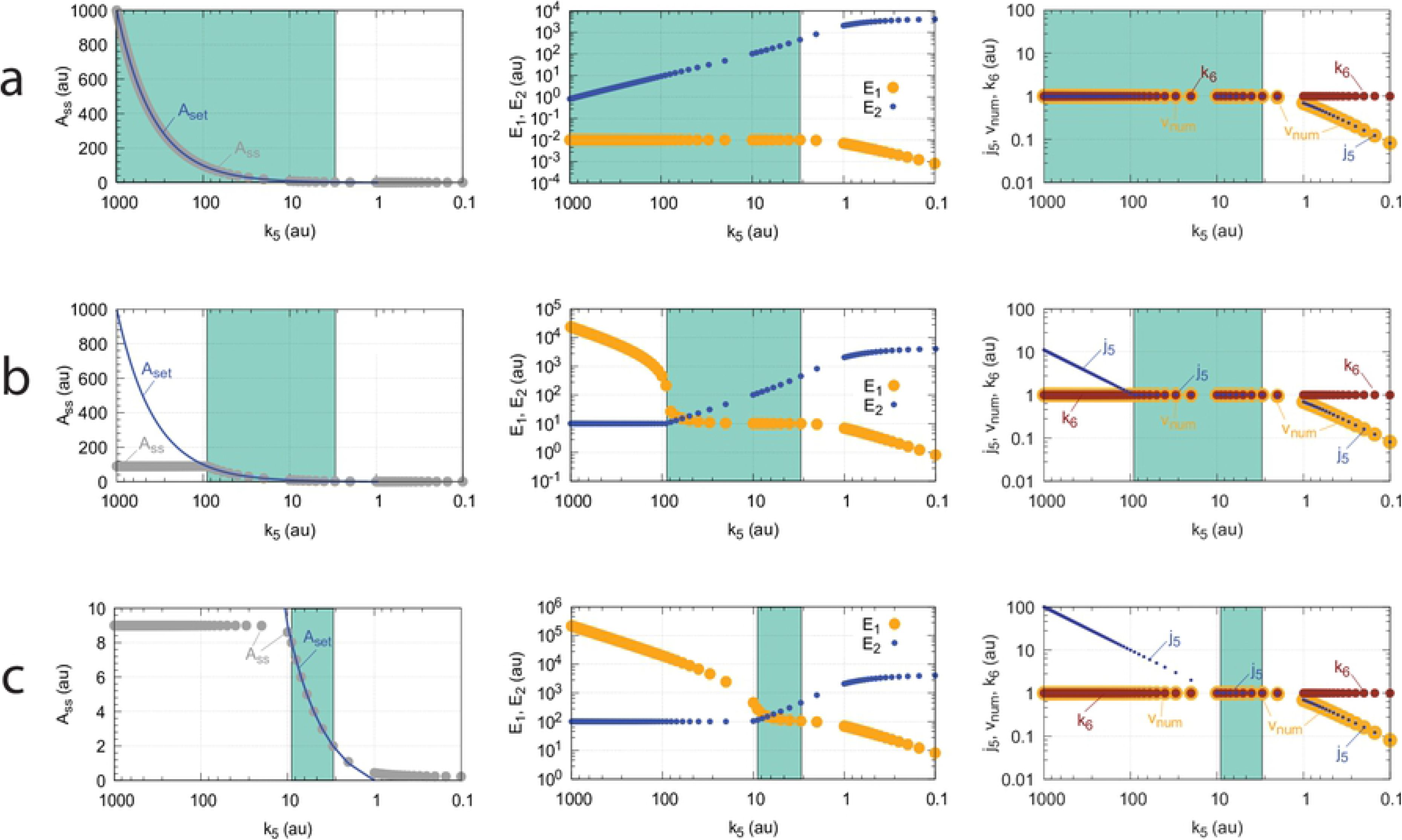
Operational range of the controller from Fig 19 as a function of *k*_5_ and the ratios (*k*_10_/*k*_9_)=(*k*_12_/*k*_11_)=(*k*_14_/*k*_13_)=(*k*_16_/*k*_15_). *A_set_* in the left panels is the theoretical set-point described by Eq 56. *A_ss_* (gray dots) are the numerically calculated steady state values of *A*. Middle panels show the concentrations of *E*_1_ and *E*_2_ indicated by blue and orange dots, respectively. Panels to the right show the flux *j*_5_ (small blue dots) which generates *E*_1_ by *A*-repression (Eq 56). *v_num_* (yellow dots) is the numerically calculated degradation velocity of the ternary-complex. Dark red dots show *k*_6_. Turquoise areas indicate the controllers operational range when Eq 56 is satisfied. (a) *k*_9_=*k*_11_=*k*_13_=*k*_15_=1×10^8^. (b) *k*_9_=*k*_11_=*k*_13_=*k*_15_=1×10^5^. (c) *k*_9_=*k*_11_=*k*_13_=*k*_15_=1×10^4^. Remaining rate constants and initial concentrations are as in Fig 20.

Interestingly, also in these calculations critical slowing down is observed, similar as in Fig 14b, when the border between dual-E control (turquoise area) and constant *A_ss_* values is approached with increasing *k*_5_ values.

Importantly, unlike the corresponding m2-controller which goes into a regime of defended single-E control under zero-order conditions (Fig 9), the constant *A_ss_* regime of the m4 controller *is not defended*, but *A_ss_* decreases with increasing *k*_2_ (perturbation) values. This is shown in Fig 23, where the (*k*_10_/*k*_9_)=(*k*_12_/*k*_11_)=(*k*_14_/*k*_13_)=(*k*_16_/*k*_15_) ratios are kept constant at 1×10^−4^, while *k*_2_ is changed from 50 (panel a) to 500 (panel b). Finally, in panel c *k*_2_=5000. With increasing *k*_2_ values a reduction in *A_ss_* and the controller’s operational range is observed.

**Fig 23.**
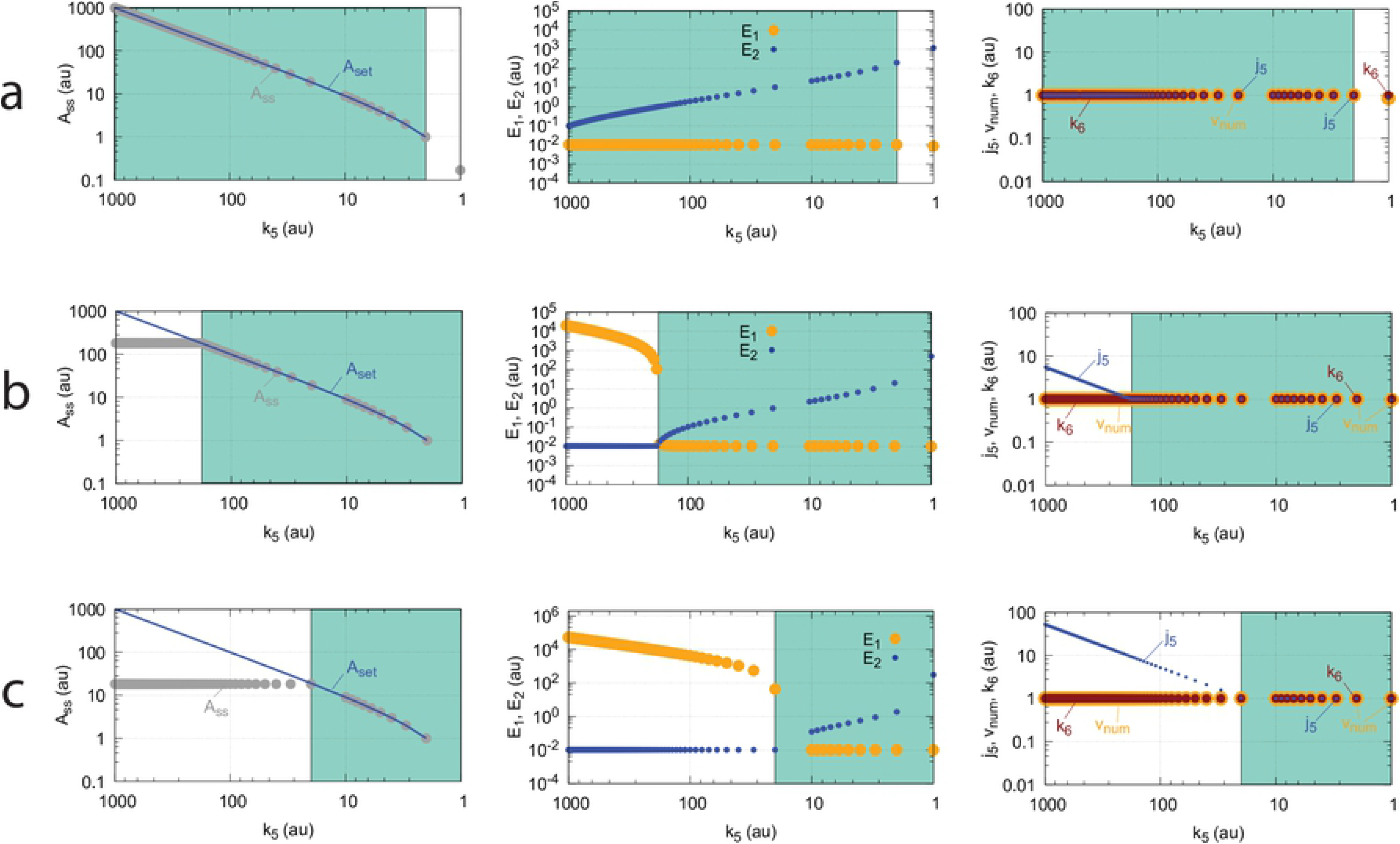
Influence of *k*_2_ on the operational range of the m4 controller Fig 19. See Fig 22 for explanation of symbols. (a) *k*_2_ = 50.0, (b) *k*_2_ = 500.0, (c) *k*_2_ = 5000.0. Other rate constants and initial concentrations are as in Fig 20.

#### Motif 4 dual-E controller removing *E*_1_ and *E*_2_ by compulsory-order ternary-complex mechanisms

Fig 24 shows the two mechanisms when the removal of *E*_1_ and *E*_2_ goes through a compulsory-order ternary-complex. In panel a *E*_1_ binds first to the free enzyme *Ez*, while in panel b *E*_2_ binds first.

**Fig 24.**
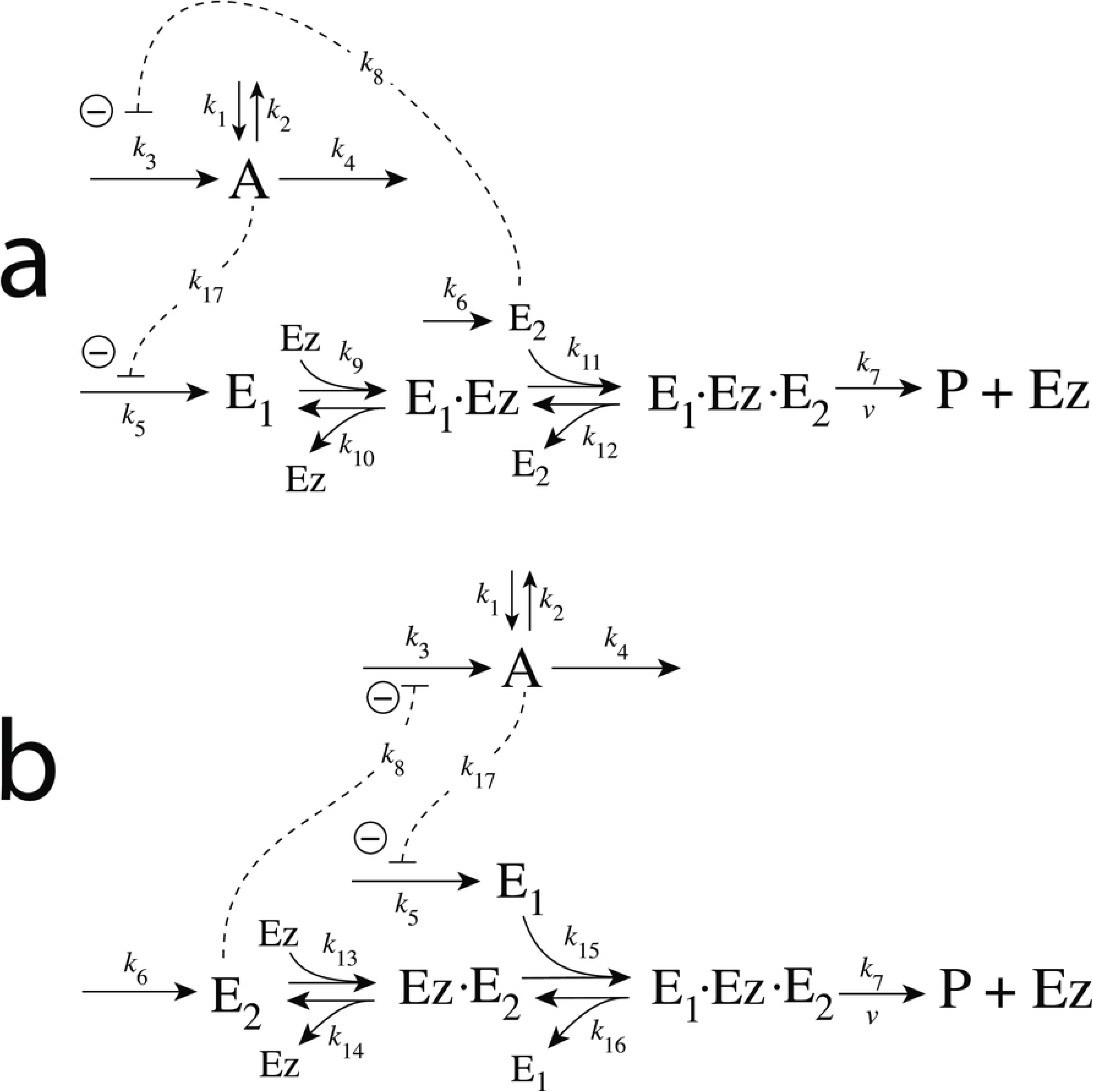
Reaction schemes when *E*_1_ and *E*_2_ in a m4-type of control structure (Fig 3) are removed by enzyme *Ez* with two compulsory-order ternary-complex mechanism. In (a) *E*_1_ binds first to *Ez*, while in (b) *E*_2_ binds first.

In case *E*_1_ binds first to *Ez* (Fig 24a), the rate equations are:

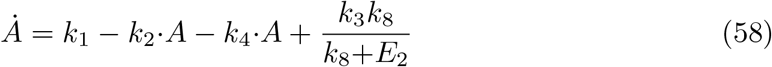

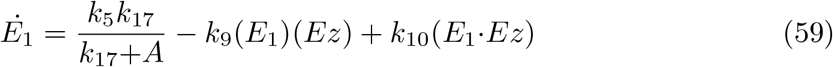

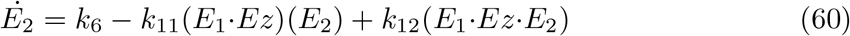

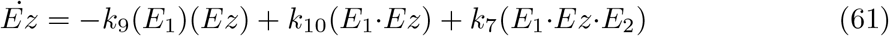

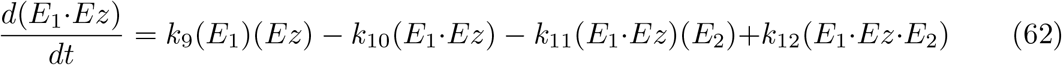

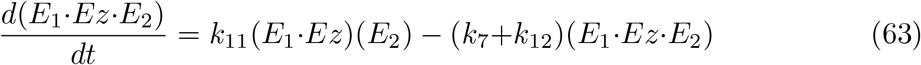

When *E*_2_ is binding first to free *Ez* (Fig 24b), the rate equations are:

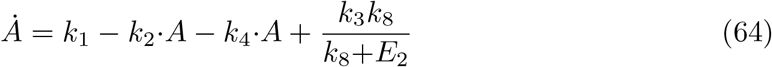

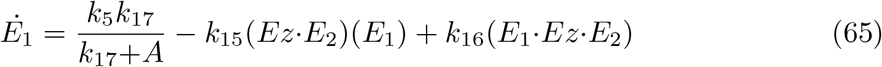

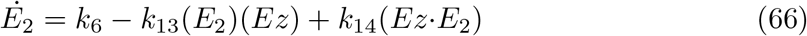

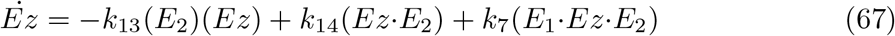

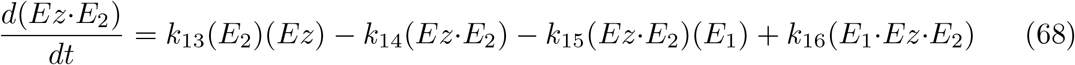

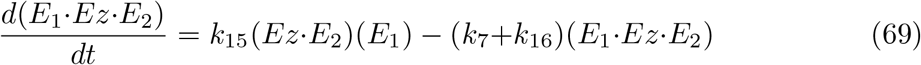

For both reaction schemes the set-point for the dual-E controller

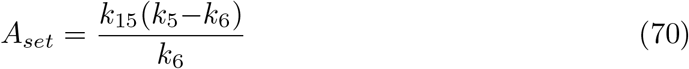

is given by the same balance conditions as for the m4 random-order ternary-complex mechanism, i.e., we have a balance between the two inflow rates *j*_5_=*k*_5_*k*_17_/(*k*_17_+*A*)=*k*_6_, and the outflow rate *k*_7_(*E*_1_·*Ez*·*E*_2_) (see Eq 56).

As already seen for the m2-controller (Fig 12) when working in dual-E mode, the random-order and compulsory-order ternary-complex mechanisms show for the m4-feedback schemes the same kinetic behavior on step-wise changes in *k*_2_. Fig 25 illustrates this for the three m4-controllers removing *E*_1_ and *E*_2_ by enzymatic ternary-complex mechanisms (Fig 19 and Fig 24). Even the breakdown at large *k*_2_ values show identical kinetics in *A* (Fig 25d).

**Fig 25.**
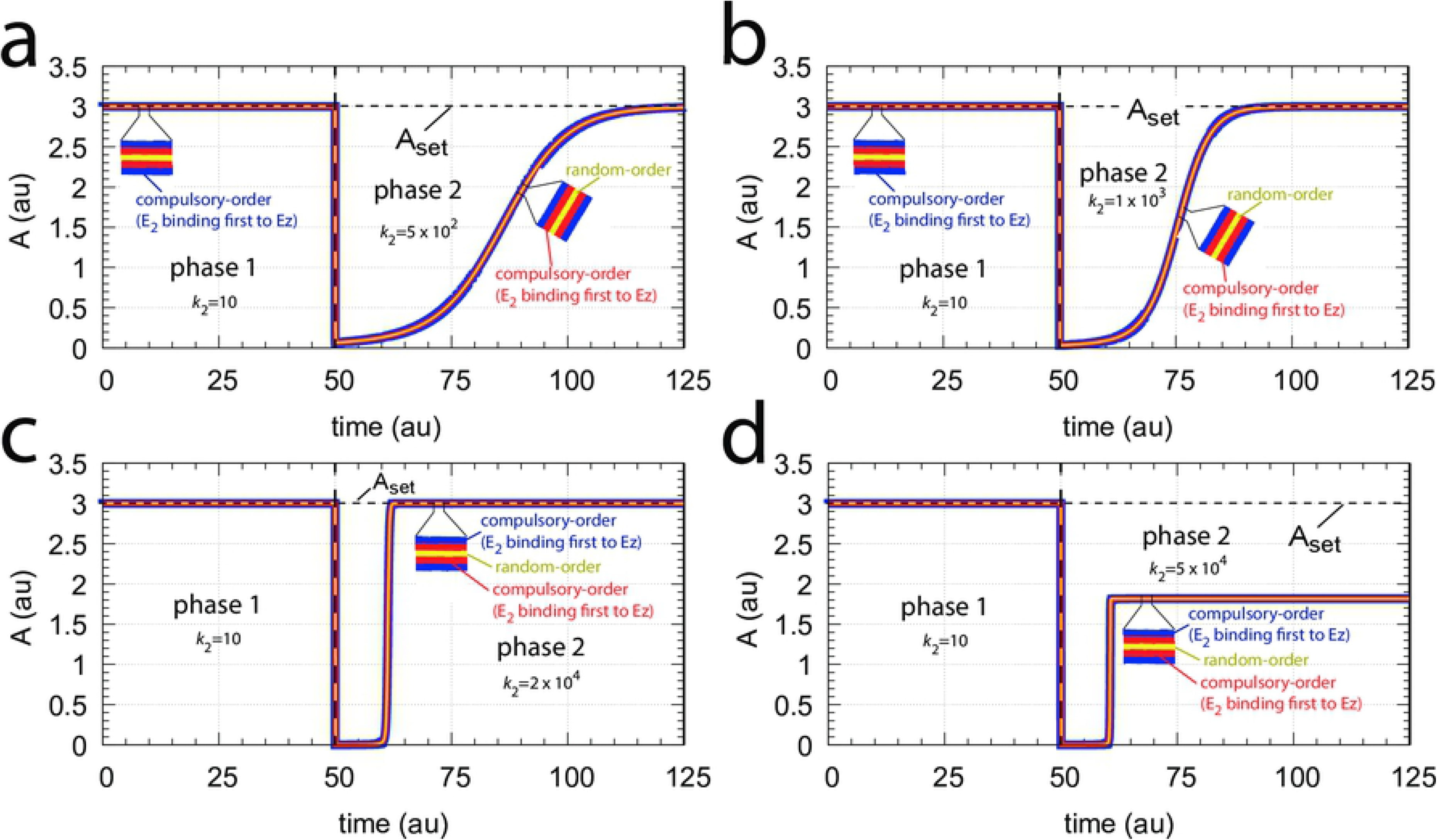
Comparison between the three m4-controllers when *E*_1_ and *E*_2_ are removed by enzymatic ternary-complex mechanisms (Fig 19 and Fig 24) upon step-wise changes at time *t*=50 from *k*_2_=10 to (a) *k*_2_=500, (b) *k*_2_=1×10^3^, (c) *k*_2_=2×10^4^, (d) *k*_2_=5×10^4^. Color coding: Thick blue line, compulsory-order mechanism with *E*_2_ binding first to *Ez*; overlaid red line, compulsory-order mechanism with *E*_1_ binding first to *Ez*; top overlaid yellow line, random-order mechanism. Rate constants and initial concentrations as for the random-order ternary-complex mechanism (Fig 20).

Fig 26 shows the concentration profiles of *E*_1_, *E*_2_ and the different enzyme species for the three m4 controller arrangements in case of their breakdown described in Fig 25d.

**Fig 26.**
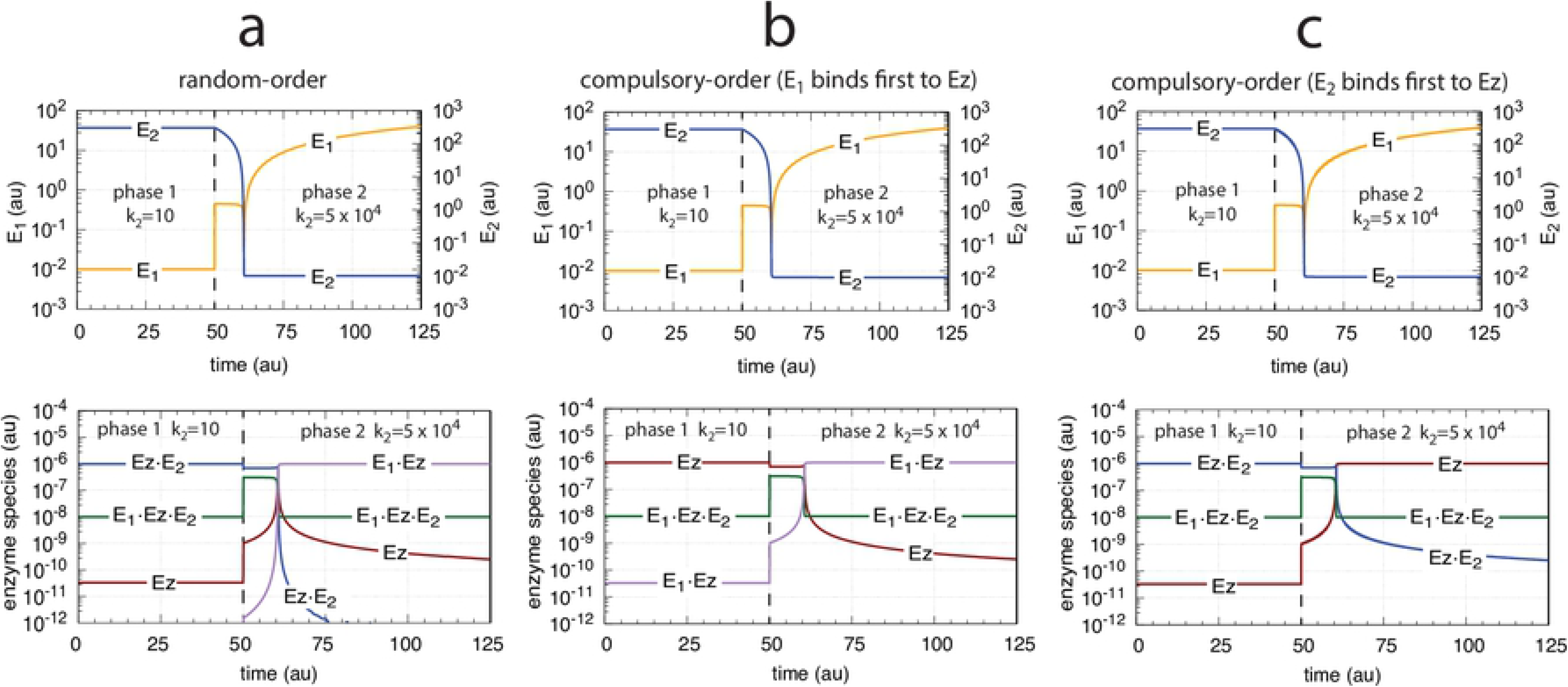
Concentration profiles of *E*_1_, *E*_2_, and enzyme species with respect to the controllers’ breakdown shown in Fig 25d. Column a: Random-order mechanism (Fig 19). Column b: Compulsory-order mechanism (Fig 24a). Column c: Compulsory-order mechanism (Fig 24b). Rate constants and initial concentrations as in Fig 25.

Although the concentration profiles of *A*, *E*_1_, *E*_2_, and the ternary-complex (*E*_1_·*Ez*·*E*_2_) are identical for the three controller configurations the other enzyme species replace each other in their functions. For example, when *E*_1_ binds first in the compulsory-order mechanisms of Fig 24a the complex (*E*_1_·*Ez*) is low during phase 1 but becomes close to the total enzyme concentration Ez_tot_ during the breakdown in phase 2 (Fig 26b). In the compulsory-order mechanism when *E*_2_ binds first (Fig 24b) the role of (*E*_1_·*Ez*) is now taken over by the free enzyme *Ez* (Fig 26c). In the random-order mechanism the role of the enzyme species is slightly more complex: during phase 1 *Ez* and (*Ez*·*E*_2_) have the same concentration profiles as in the compulsory-order mechanism where *E*_2_ binds first to *Ez* (Fig 24b). However, in phase 2 the *Ez* profile of the random-order mechanism is that of the other compulsory-order mechanism (Fig 26b)!

#### Motif 4 dual-E enzymatic controller in which *E*_1_ and *E*_2_ are removed by ping-pong mechanisms

Fig 27 shows the two possibilities when enzyme *Ez* removes *E*_1_ and *E*_2_ by a ping-pong mechanism. In panel a *E*_1_ binds to the free enzyme and creates the alternative enzymatic form *Ez*^*^, which then bind the derepressing controller species *E*_2_. In panel b this is reversed. Here *E*_2_ binds first and forms *Ez*^*^, which can bind *E*_1_. As for the m2 controller case we have, for the sake of simplicity, omitted the release of a product prior to the formation of *Ez*^*^.

**Fig 27.**
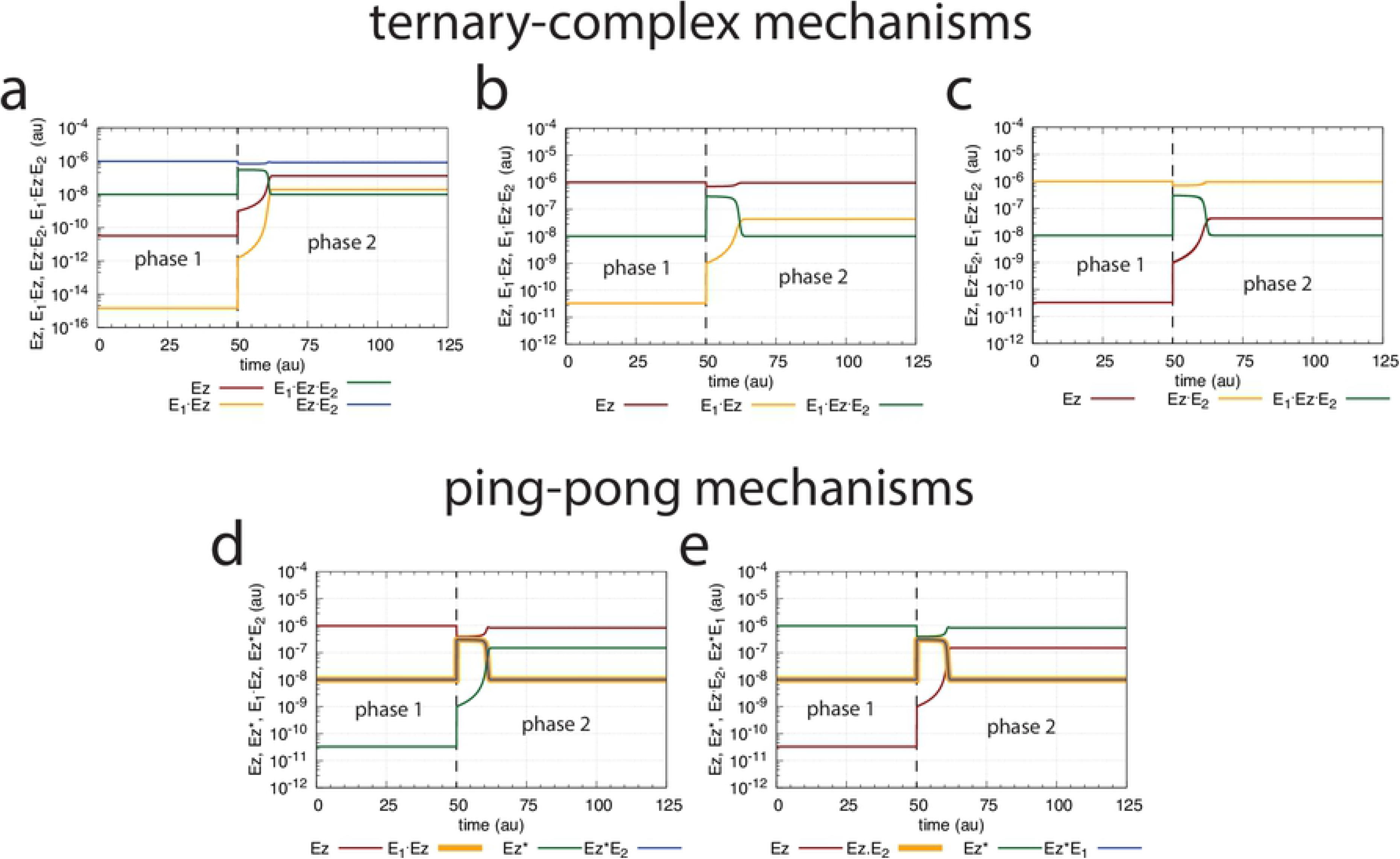
Reaction schemes when *E*_1_ and *E*_2_ in a m4-type of control structure (Fig 3) are removed by enzyme *Ez* following two ping-pong mechanisms. In (a) *E*_1_ binds first to the free enzyme *Ez*, while in (b) *E*_2_ binds first.

For the scheme of Fig 27a the rate equations are:

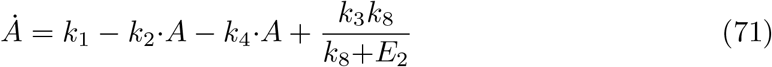

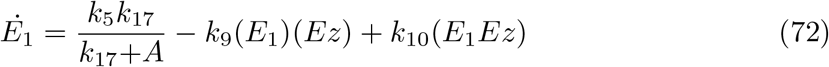

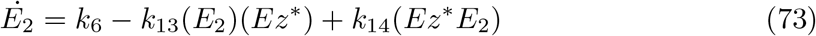

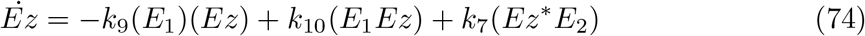

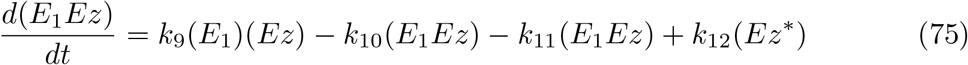

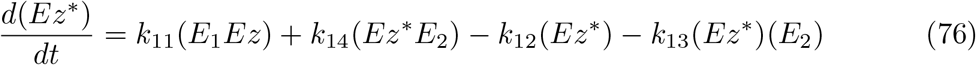

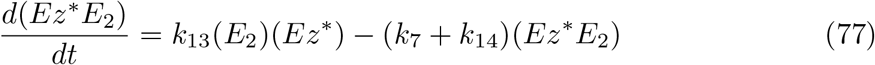

In case *E*_2_ binds first to *Ez* (Fig 27b), the rate equations are:

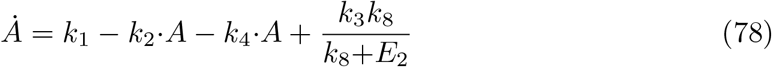

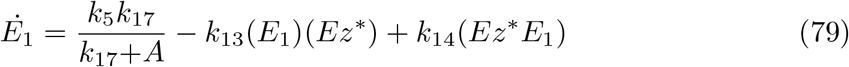

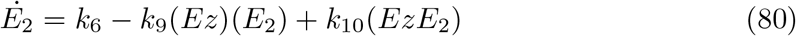

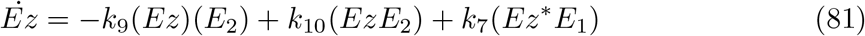

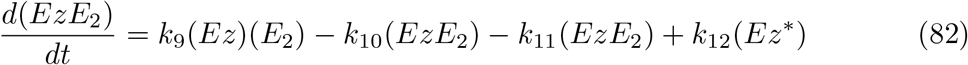

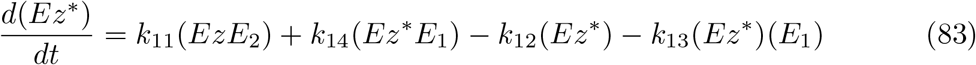

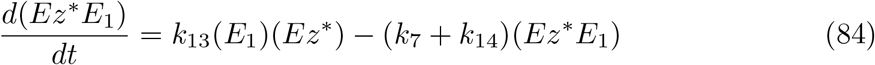

We have compared the two m4 ping-pong mechanisms (Fig 27) with the three m4 ternary-complex mechanisms (Fig 19 and Fig 24) and found that their homeostatic behavior in *A* as well as the concentration profiles in *E*_1_ and *E*_2_ have identical dynamics with those shown in Fig 25 and the upper row in Fig 26, respectively (data not shown). However, despite their identical dynamical behaviors in the controlled variable *A* as well as in the controller variables *E*_1_ and *E*_2_ the different enzyme species show, like in the lower row of Fig 26, a mechanism-dependent restructuring of the enzyme species’ concentration profiles. This indicates that in the different mechanisms different enzyme species take over the tasks to decrease *E*_2_ (causing an increase in the compensatory flux when *k*_2_ is increased) and to increase *E*_1_, thereby leading to homeostasis in *A*. Specifically, for the intact m4 dual-E *ternary-complex controllers* (i.e. no breakdown occurs) the condition of Eq 56 defines the profiles of the enzyme species, while for the m4 ping-pong controllers the conditions

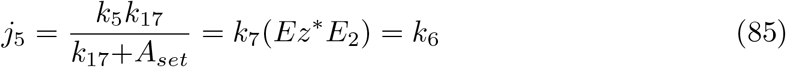

or

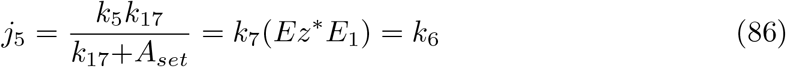

determine the enzyme species concentration profiles when *E*_1_ or *E*_2_ bind first to *Ez*, respectively (see Fig 27). In both cases the set-point is

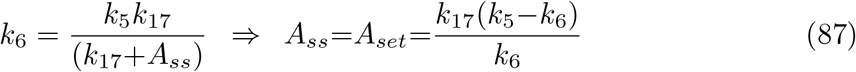

Fig 28 illustrates the concentration profiles of the enzyme species when all five mechanisms show the same homeostatic behavior in *A* as in Fig 25c with identical changes in *E*_1_ and *E*_2_.

**Fig 28.**
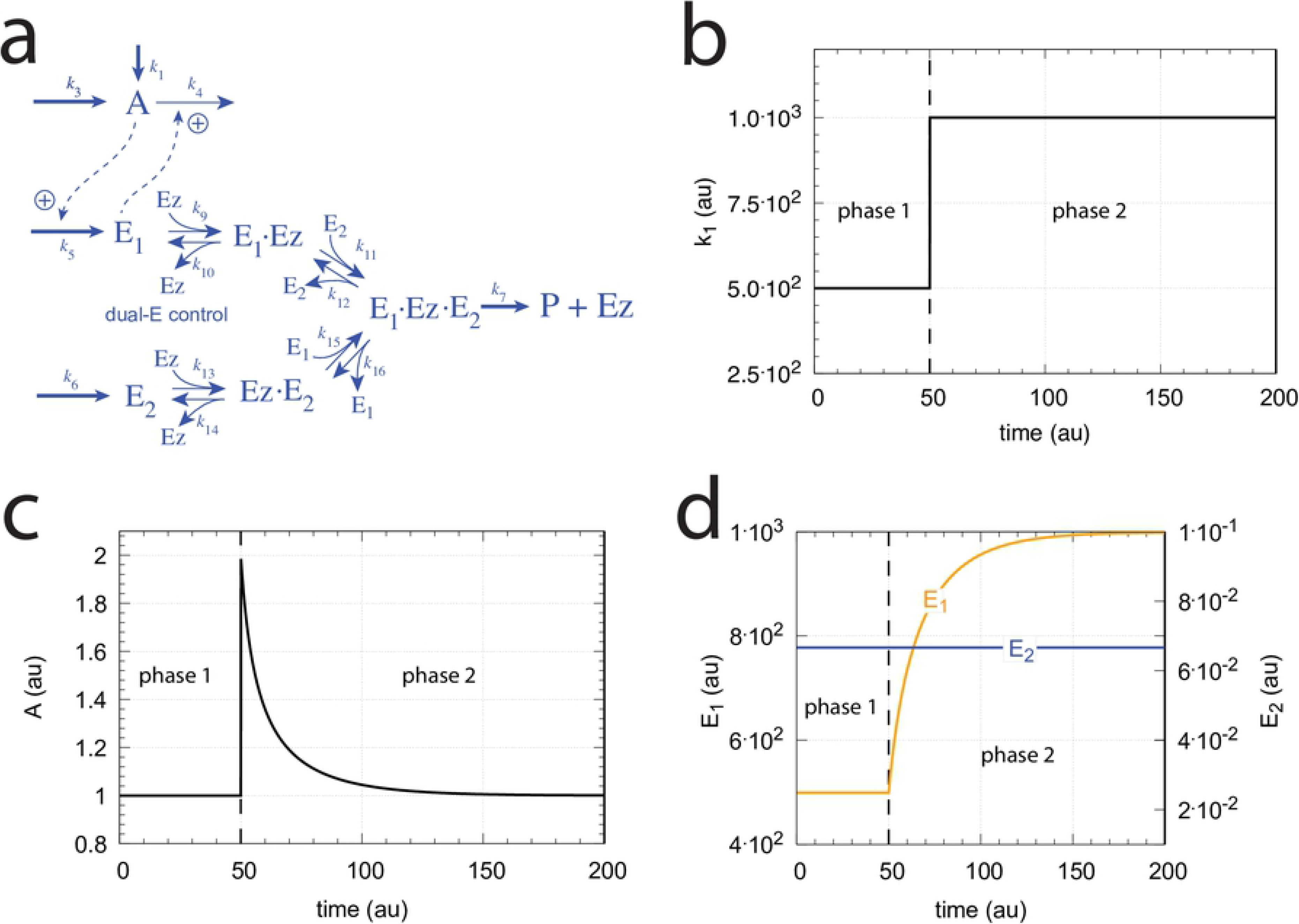
Enzyme species profiles of the m4 ternary-complex (Fig 19 and Fig 24) and ping-pong mechanisms (Fig 27) when *k*_2_=10 in phase 1, and *k*_2_=2×10^4^ in phase 2. (a) random-order ternary-complex mechanism, (b) compulsory-order ternary-complex mechanism with *E*_1_ binding first to *Ez*, (c) compulsory-order ternary-complex mechanism with *E*_2_ binding first to *Ez*, (d) ping-pong mechanism with *E*_1_ binding first to *Ez*, (e) ping-pong mechanism with *E*_2_ binding first to *Ez*. Rate constants (if applicable) are as in Fig 20. Initial concentrations: (a) A_0_=3.0, E_1,0_=1.0×10^−2^, E_2,0_=3.0×10^2^, Ez_0_=3.3×10^−11^, (E_1_·Ez)_0_=1.4×10^−15^, (E_1_·Ez·E_2_)_0_=1.0×10^−8^, (EzE_2_)_0_=9.9×10^−7^. (b) A_0_=3.0, E_1,0_=1.0×10^−2^, E_2,0_=3.0×10^2^, Ez_0_=9.9×10^−7^, (E_1_·Ez)_0_=3.3×10^−11^, (E_1_·Ez·E_2_)_0_=1.0×10^−8^. (c) A_0_=3.0, E_1,0_=1.0×10^−2^, E_2,0_=3.0×10^2^, Ez_0_=3.3×10^−11^, (EzE_2_)_0_=9.9×10^−7^, (E_1_·Ez·E_2_)_0_=1.0×10^−8^. (d) A_0_=3.0, E_1,0_=1.0×10^−2^, E_2,0_=3.0×10^2^, Ez_0_=9.8×10^−7^, (E_1_·Ez)_0_=1.0×10^−8^, 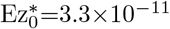, (Ez^*^E_2_)_0_=1.0×10^−8^. (e) A_0_=3.0, E_1,0_=1.0×10^−2^, E_2,0_=3.0×10^2^, Ez_0_=3.3×10^−11^, (Ez^*^E_1_)_0_=1.0×10^−8^, 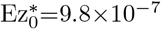, (EzE_2_)_0_=1.0×10^−8^.

As an example, in the ping-pong mechanisms the role of the ternary-complex *E*_1_·*Ez·E*_2_ (Figs 28a-c, outlined in green) is replaced by *Ez*^*^*E*_2_ (Fig 28d, *E*_1_ binding first to *Ez*) or by *Ez*^*^*E*_1_ (Fig 28e, *E*_2_ binding first to *Ez*) as implied by Eqs 56, 85, and 86. Likewise, the steady state concentrations of the other enzyme species can be derived from the above rate equations (see the King-Altman expressions in the Supporting information), but are not further elaborated here.

### Controllers based on motif 5

As indicated in Fig 2, motif m5 is an outflow controller [6] and opposes inflow perturbations on the controlled variable *A*. Like the m2 controller the dual-E (antithetic) version of m5 has an “inner-loop” signaling (Fig 3).

#### Motif 5 dual-E controller removing *E*_1_ and *E*_2_ by a random-order ternary-complex mechanism

Fig 29a shows the reaction scheme when in a m5 controller configuration *E*_1_ and *E*_2_ are removed by an enzymatic random-order ternary-complex mechanism.

**Fig 29.**
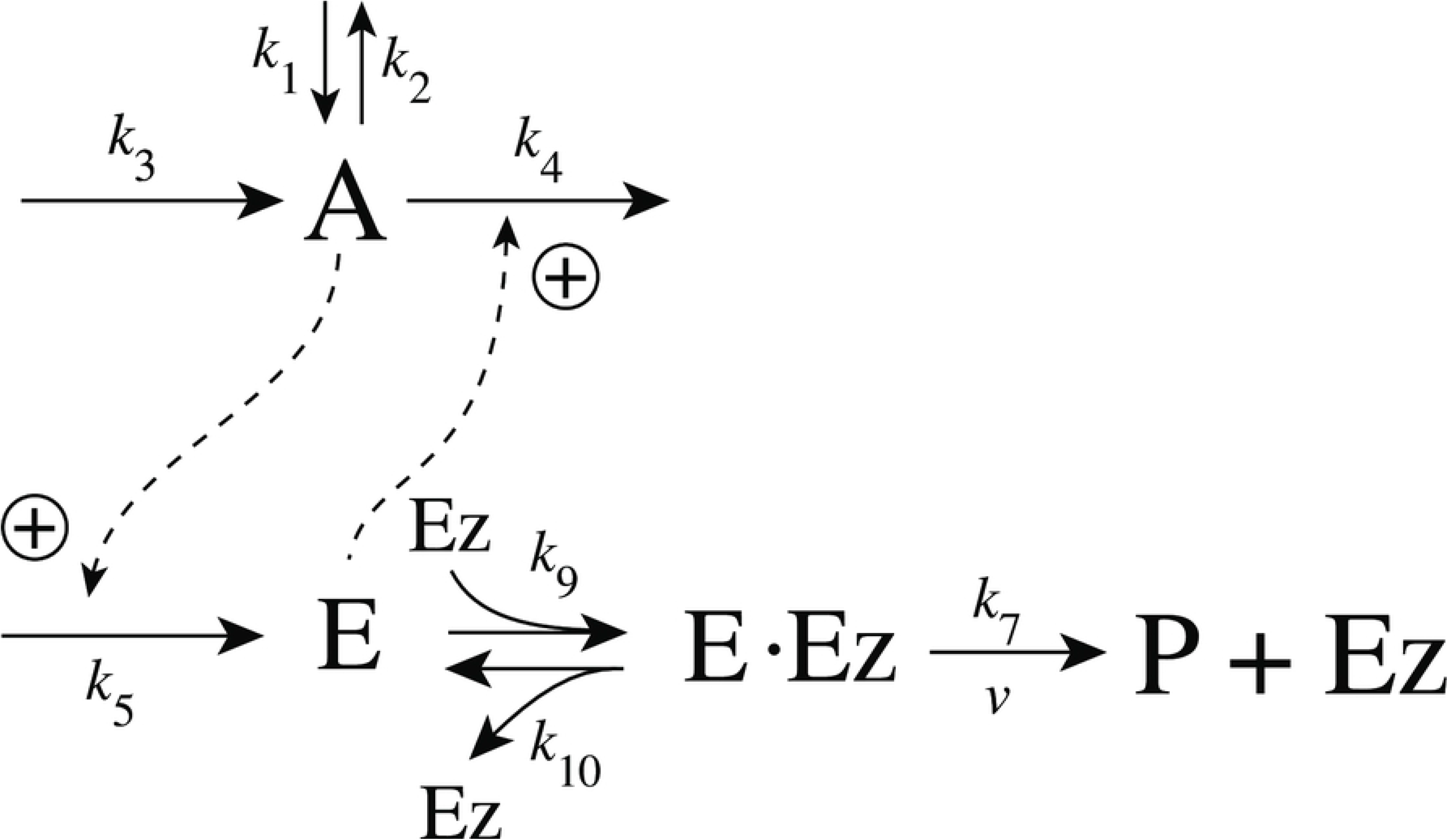
Example of m5 feedback loop where *E*_1_ and *E*_2_ are removed by a random-order ternary-complex mechanism which works under dual-E control. (a) Reaction scheme. (b) Step-wise change of *k*_1_ from 500.0 to 1000.0 at time *t* = 50. (c) In dual-E mode the set-point is *A_set_*=*k*_6_/*k*_5_ (= 1.0) which is defended. The panel shows the response of *A* with respect to the step-wise change of *k*_1_ in panel (a). (d) Change of *E*_1_ and *E*_2_ in response to the step-wise change of *k*_1_ in panel (a). Rate constants: *k*_1_=500.0 (phase 1), *k*_1_=1000.0 (phase 2), *k*_2_=1.0, *k*_3_=0.0, *k*_4_=1.0, *k*_5_=40.0, *k*_6_=40.0 *k*_7_=1×10^8^, *k*_9_=*k*_11_=*k*_13_=*k*_15_=1×10^9^, *k*_10_=*k*_12_=*k*_14_=*k*_16_=1×10^3^. Initial concentrations: A_0_=1.0, E_1,0_=499.0, E_2,0_=6.67×10^−2^, Ez_0_=8.02×10^−11^, (E_1_·Ez)_0_=5.99×10^−7^, (E_1_·Ez·E_2_)_0_=4.0×10^−7^, (EzE_2_)_0_=1.15×10^−14^.

The rate equations are:

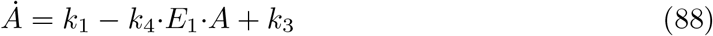

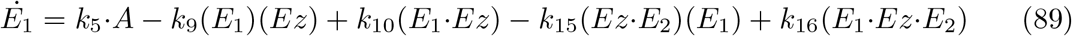

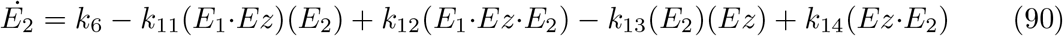

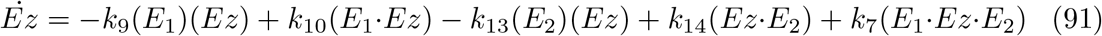

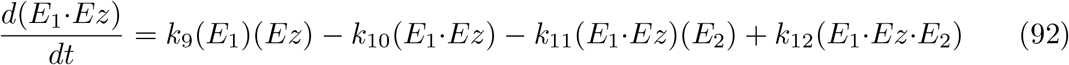

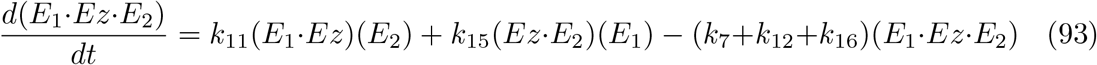

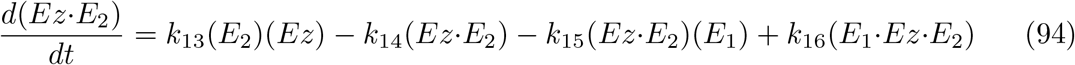

The corresponding single-E controller is shown in Fig 30

**Fig 30.**
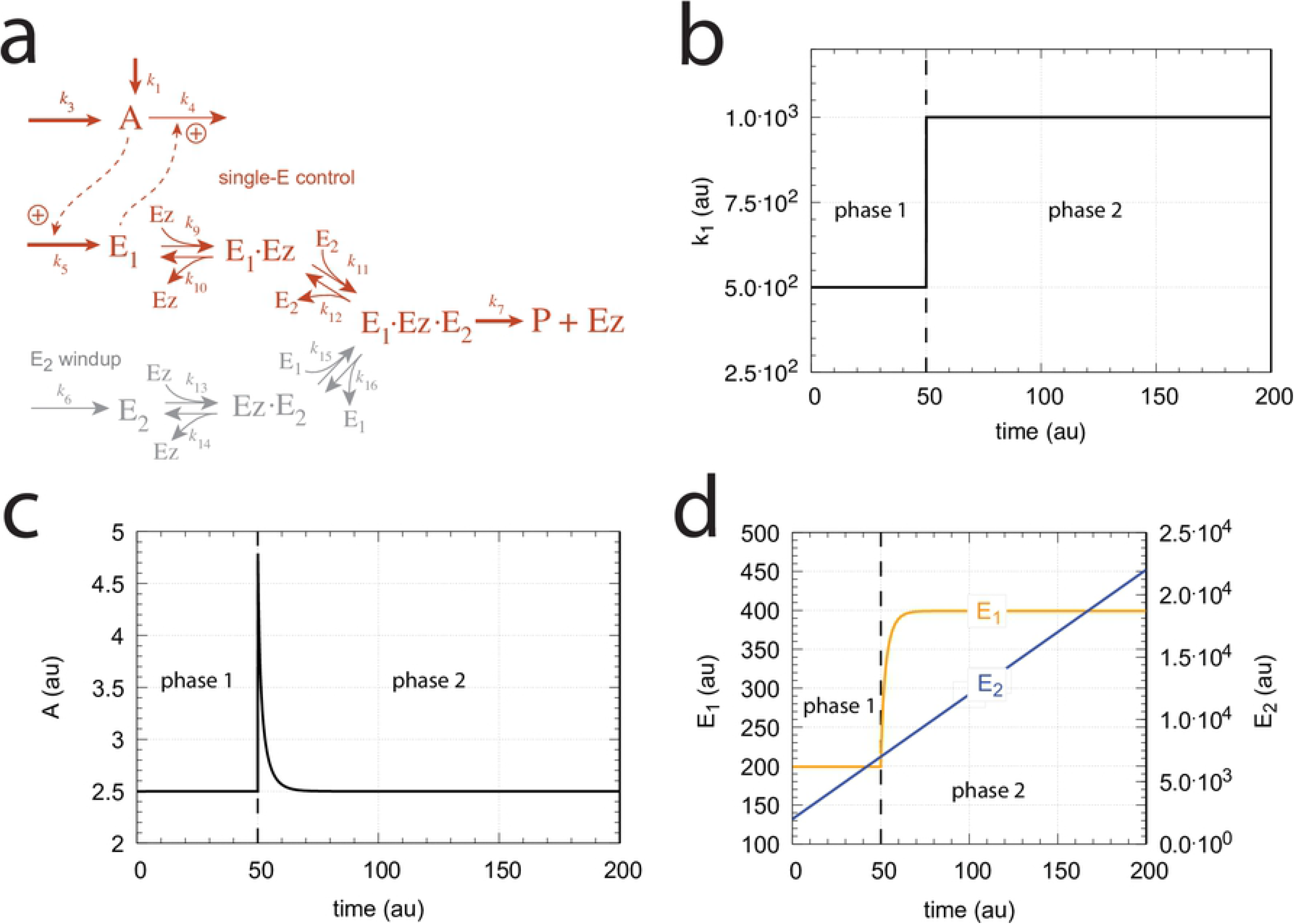
Reaction scheme of the catalyzed single-E m5-type of controller.

with the corresponding rate equations:

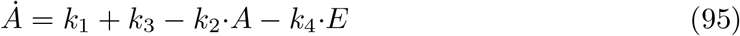

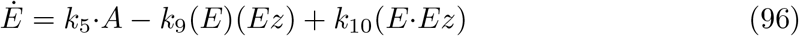

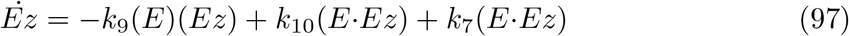

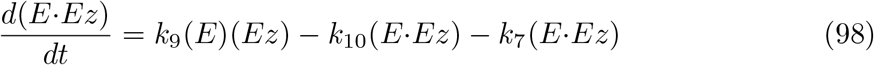

The single-E feedback loop in Fig 30 shows robust homeostatic control when enzyme *Ez* works under zero-order conditions, i.e., *K_M_* =(*k*_10_+*k*_7_)/*k*_9_ is low and *k*_9_≫*k*_10_+*k*_7_. In this case the set-point for *A* is given by the condition

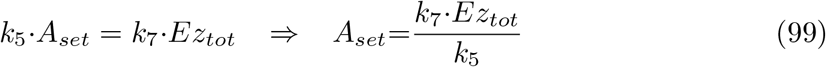

For the dual-E m5 controller (Fig 29a) robust homeostasis in *A* is obtained by the condition

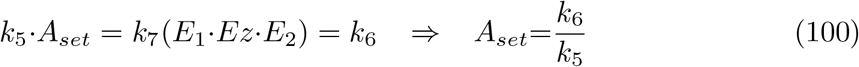

Figs 29b-d show the controller’s behavior upon a step-wise change in *k*_1_ when working in dual-E mode, i.e., when the homeostatic set-point for *A* is given by Eq 100.

A switch from dual-E to single-E control mode occurs when *k*_6_ becomes larger than *k*_7_*Ez_tot_*. For large *k*_9_/*k*_10_, *k*_11_/*k*_12_, *k*_13_/*k*_14_, and *k*_15_/*k*_16_ ratios *A_set_* of the single-E controller is given by the condition

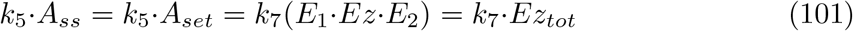

Fig 31 gives an example, where *k*_6_ has been increased to 200.0, while the other rate constant values are as in Fig 29. Fig 31a shows the operative part of the single-E controller outlined in red. The grayed part does not participate in the control of *A*, but shows a steady increase of *E*_2_ (wind-up). The controller is subject to the same step-wise increase as in *k*_1_ (panel b) as in Fig 29, but has now changed its set-point to 2.5 as described by Eq 99 (panel c). Fig 31d shows the wind-up behavior of *E*_2_ along with the change of *E*_1_, which activates the compensatory flux removing *A* and compensating for the increasing inflow of *A* by *k*_1_.

**Fig 31.**
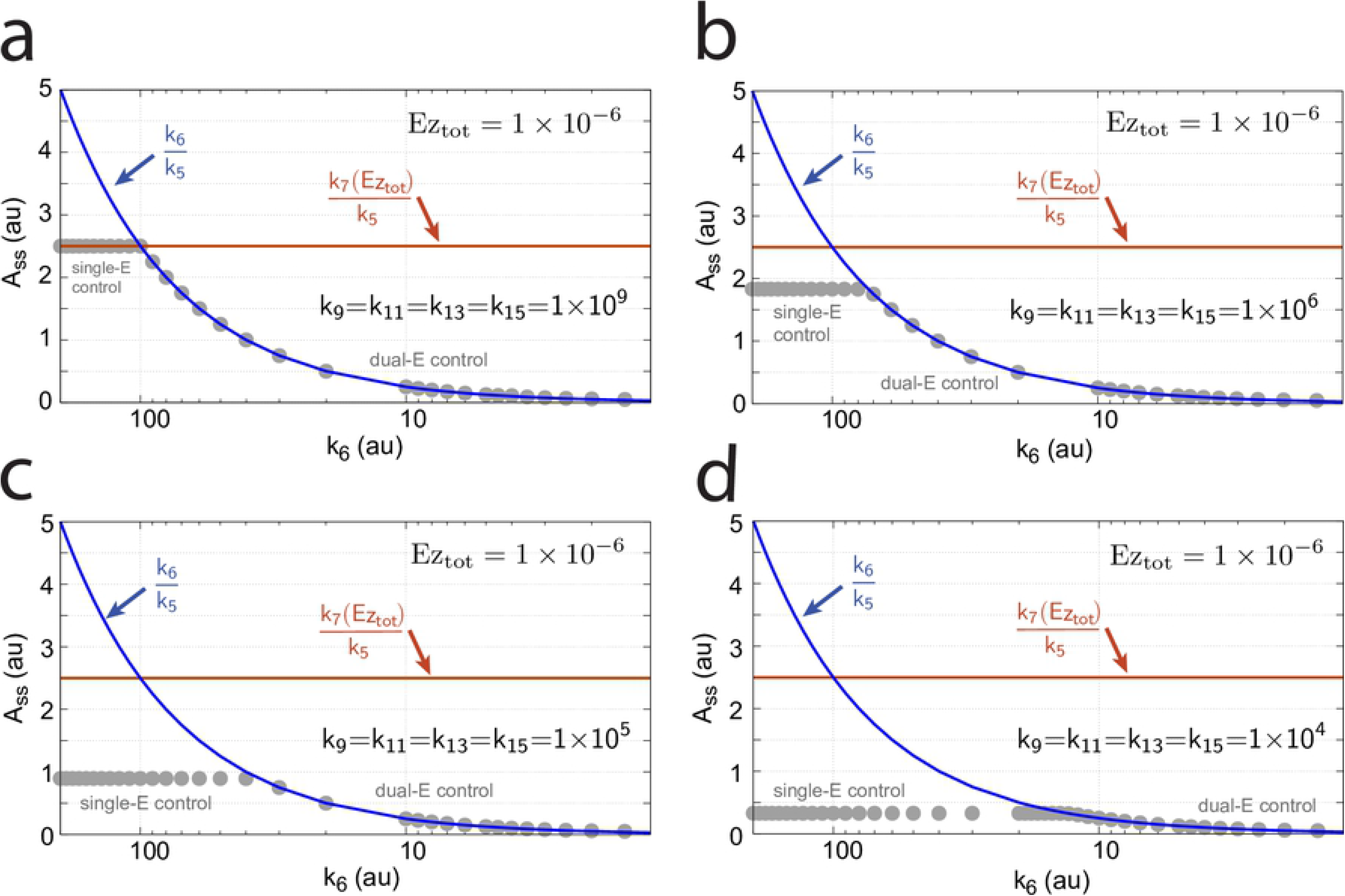
Example of the m5 feedback loop with *E*_1_ and *E*_2_ being removed by a random-order ternary-complex mechanism working in single-E control mode. (a) Scheme outlined in red shows part of the network participating in the control of *A*. (b) Step-wise change of *k*_1_ from 500.0 to 1000.0 at time *t* = 50.0. (c) Homeostatic response of *A*, i.e. the controller defends its set-point (=2.5) defined by Eq 99. (d) Change of *E*_1_ and wind-up of *E*_2_. Rate constants as in Fig 29, except that *k*_6_=200. Initial concentrations: A_0_=2.5, E_1,0_=199.2, E_2,0_=2.0×10^3^, Ez_0_=4.54×10^−11^, (E_1_·Ez)_0_=4.51×10^−12^, (E_1_·Ez·E_2_)_0_=9.995×10^−7^, (EzE_2_)_0_=4.56×10^−10^.

At low *k*_9_/*k*_10_, *k*_11_/*k*_12_, *k*_13_/*k*_14_, and, *k*_15_/*k*_16_ ratios the operational range of the dual-E controller decreases and the single-E controller’s steady state in *A* drops below *A_set_*. Under these conditions the dual-E controller will defend its set-point *A_set_*=*k*_6_/*k*_5_ exactly, while the single-E controller shows an offset, i.e. *A_ss_<A_set_* = *k*_7_ *Ez_tot_/k*_5_. Fig 32 illustrates this behavior when the total enzyme concentration is kept constant at 1×10^−6^.

**Fig 32.**
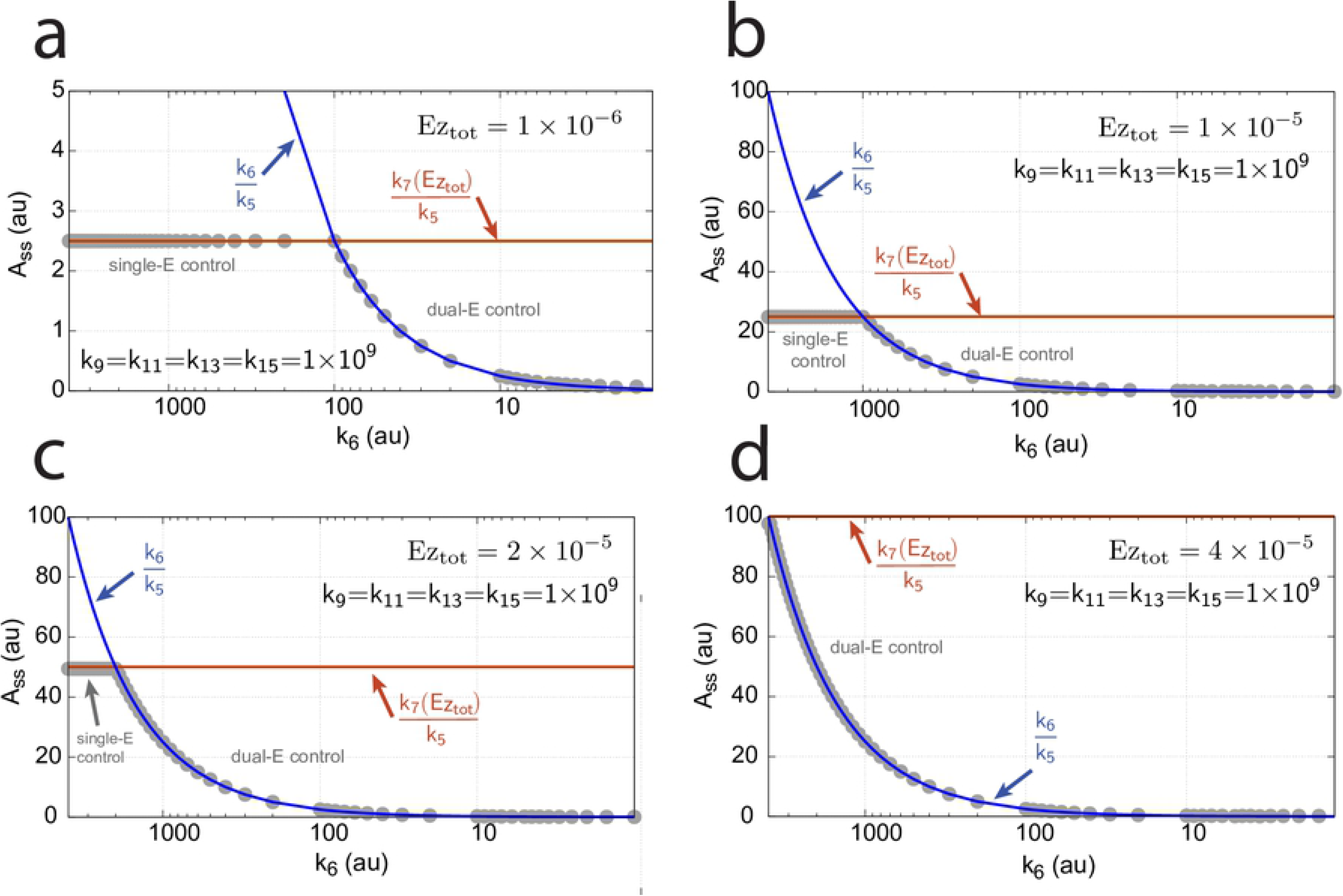
Switching between single-E control (Fig 31) and dual-E control (Fig 29) as a function of *k*_6_ for different values of *k*_9_, *k*_11_, *k*_13_, and *k*_15_. Panel (a): high value (1×10^9^) of *k*_9_, *k*_11_, *k*_13_, and *k*_15_. The dual-E controller shows its maximum operational range. In this case the switch occurs when *k*_6_>*k*_7_*Ez_tot_*. Panels (b)-(d): for the lower values of *k*_9_, *k*_11_, *k*_13_, and *k*_15_ (indicated inside the figure) the ternary-complex concentration (*E*_1_·*Ez*·*E*_2_) is lower than *Ez_tot_* and the switch occurs at lower *k*_6_ values, which leads to a decreased operational range of the dual-E controller. Due to the lower (*E*_1_·*Ez·E*_2_) concentration the single-E control mode (which occurs analogous to the red-outlined part in Fig 9e) shows an offset below *k*_7_*Ez_tot_/k*_5_. Note however, that *A_ss_* will depend on the perturbation *k*_1_ and move towards *A_set_* with increasing *k*_1_, thereby reducing the single-E controller’s offset. Other rate constants: *k*_1_=500.0, *k*_2_=1.0, *k*_3_=0.0, *k*_4_=1.0, *k*_5_=40.0, *k*_7_=1×10^8^, *k*_10_=*k*_12_=*k*_14_=*k*_16_=1×10^3^. Initial concentrations: A_0_=2.0, E_1,0_=5.49×10^−2^, E_2,0_=5.21×10^3^, Ez_0_=7.4×10^−14^, (E_1_·Ez)_0_=9.09×10^−8^, (E_1_·Ez·E_2_)_0_=9.09×10^−7^, (EzE_2_)_0_=1.66×10^−10^.

When *Ez_tot_* increases the operational range of the dual-E controller increases as a function of *k*_6_. This is shown in Fig 33 when *k*_9_=*k*_11_=*k*_13_=*k*_15_=1×10^9^ and *Ez_tot_* varies from 1×10^−6^ to 4×10^−5^. In agreement with Eq 99 we observe that with changing *Ez_tot_* the set-point of the single-E controller changes accordingly.

**Fig 33.**
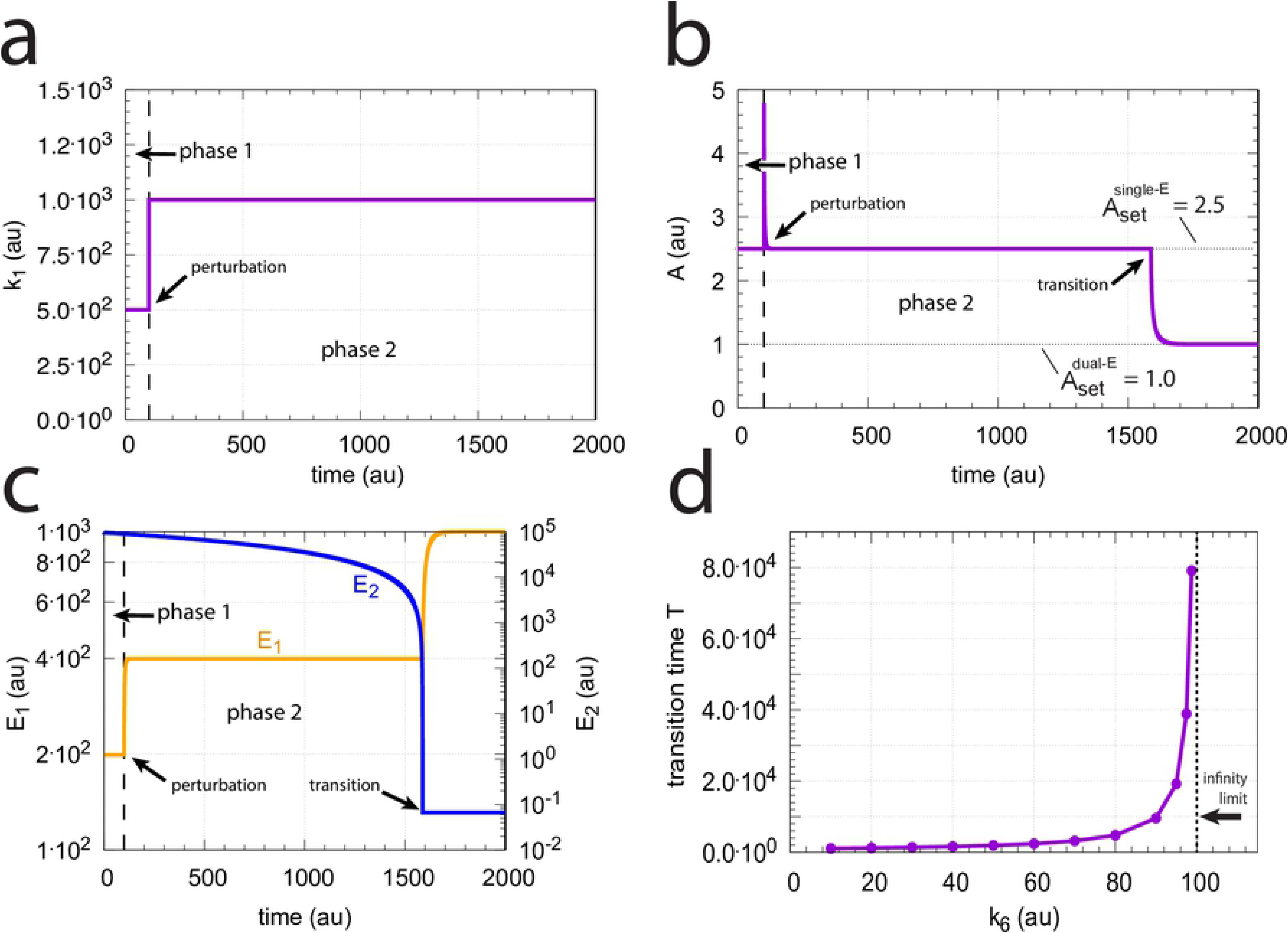
Switching between dual-E and single-E control in the random-order ternary-complex m5 controller at different *Ez_tot_* concentrations. Rate constants are as in Fig 32a with *Ez_tot_* values as indicated in the four panels. With increasing *Ez_tot_* values (from panel (a) to panel (d)) the operational range of the dual-E controller increases together with increasing set-point values of the single-E controller (Eq 99). Initial concentrations, panel (a): A_0_=2.0, E_1,0_=5.49×10^−2^, E_2,0_=5.21×10^3^, Ez_0_=7.4×10^−14^, (E_1_·Ez)_0_=9.09×10^−8^, (E_1_·Ez·E_2_)_0_=9.09×10^−7^, (EzE_2_)_0_=1.66×10^−10^. Initial concentrations panel (b): A_0_=2.5, E_1,0_=5.49, E_2,0_=5.21×10^1^, Ez_0_=1.18×10^−12^, (E_1_·Ez)_0_=9.96×10^−6^, (E_1_·Ez·E_2_)_0_=4.5×10^−8^, (EzE_2_)_0_=1.19×10^−17^. Initial concentrations panel (c): A_0_=2.5, E_1,0_=5.49, E_2,0_=5.21×10^1^, Ez_0_=1.21×10^−12^, (E_1_·Ez)_0_=1.995×10^−5^, (E_1_·Ez·E_2_)_0_=4.5×10^−8^, (EzE_2_)_0_=1.21×10^−17^. Initial concentrations panel (d): A_0_=97.63, E_1,0_=4.12, E_2,0_=9.87×10^3^, Ez_0_=3.95×10^−10^, (E_1_·Ez)_0_=1.66×10^−13^, (E_1_·Ez·E_2_)_0_=3.905×10^−5^, (EzE_2_)_0_=9.47×10^−7^.

#### Transition from single-E to dual-E control and critical slowing down

In the previous section we saw that dual-E control occurs in the m5 random-order ternary-complex mechanism when the condition *k*_6_*<k*_7_Ez_tot_ is met. In this case, both *E*_1_ and *E*_2_ are engaged in the control of *A*. On the other hand, single-E control is observed when *k*_6_>*k*_7_Ez_tot_. Here, only *E*_1_ acts as the controller variable while *E*_2_ shows wind-up. i.e., increases continuously. However, even when the condition for dual-E control is fulfilled, i.e., *k*_6_*<k*_7_Ez_tot_, single-E control can be temporarily present, as observed for the m2 controller (Fig 14), when the initial concentration of *E*_2_ is above its steady state value for a given perturbation value of *k*_1_. In this case, the single-E controller is *metastable*: *E*_2_ will decrease and approach its steady state, but during this period the set-point of the single-E controller will be defended when working under zero-order conditions. Figs 34a-c illustrates the behavior. In this example *E*_2_ concentration starts out high at 9.5×10^4^, while its steady state value is 6.7×10^−2^. At time *t* = 100 *k*_1_ is changed from 500.0 to 1000.0 (Fig 34a) and the set-point of *A* for the single-E controller (=2.5) is defended as long as *E*_2_>*E*_2,*ss*_ (Fig 34b). Note that during single-E control the *E*_1_ value is responsible for keeping *A* at its set-point, but the *E*_1_ level changes once *E*_2_ is at its steady state and the controller has reached dual-E control mode (Figs 34b and c).

**Fig 34.**
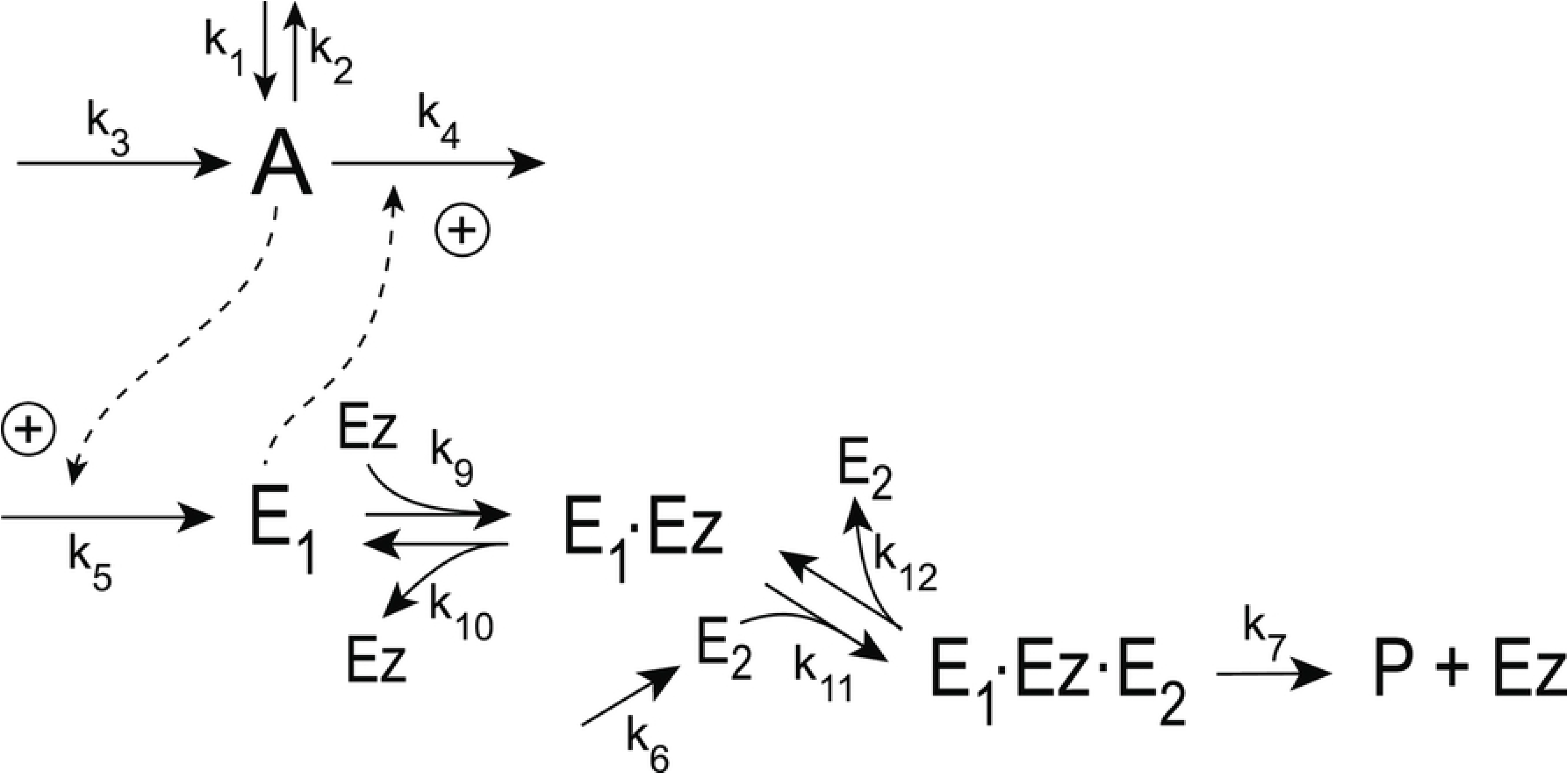
Metastable single-E controller and critical slowing down in the autonomous transition from single-E to dual-E control mode. (a) Step-wise change of *k*_1_ at t=100.0 from 500.0 (phase 1) to 1000.0 (phase 2). (b) Metastable single-E control mode. The single-E controller defends its set-point (=2.5), but transition to dual-E control mode (indicated by arrow) occurs at approximately 1600 time units when *E*_2_ reaches its steady state. (c) The metastable single-E control mode is operative as long as *E*_2_ is above its steady state value. The transition from single-E to dual-E control mode occurs when *E*_2_ has reached its steady state (indicated by arrow). Rate constants: *k*_1_=500.0 (phase 1), *k*_1_=1000.0 (phase 2), *k*_2_=1.0, *k*_3_=0.0, *k*_4_=1.0, *k*_5_=40.0, *k*_6_=40.0 *k*_7_=1×10^8^, *k*_9_=*k*_11_=*k*_13_=*k*_15_=1×10^9^, *k*_10_=*k*_12_=*k*_14_=*k*_16_=1×10^3^. Initial concentrations: A_0_=2.5, E_1,0_=199.1, E_2,0_=9.52×10^4^, Ez_0_=1.08×10^−12^, (E_1_·Ez)_0_=2.37×10^−15^, (E_1_·Ez·E_2_)_0_=9.995×10^−7^, (EzE_2_)_0_=5.01×10^−10^. (d) Transition time *T* as a function of *k*_6_. *k*_1_=500.0; all other rate constants and initial concentrations as for (a)-(c).

We have further tested how the transition time *T* depends on *k*_6_ for this controller at constant *k*_1_ (for a definition of *T* see Fig 14a). As for the m2 controller (Fig 14b) *T* increases with increasing *k*_6_ values. For each value of *k*_6_ it takes *T* time units until *A* settles at the set-point of the dual-E controller (=*k*_6_/*k*_5_). *T*→∞ as *k*_6_ approaches 100.0, the value at which the set-point of the dual-E controller approaches the set-point of the single-E controller.

#### Motif 5 dual-E controller removing *E*_1_ and *E*_2_ by a compulsory-order ternary-complex mechanism with *E*_1_ binding first to *Ez*

The scheme of this mechanism is shown in Fig 35.

**Fig 35.**
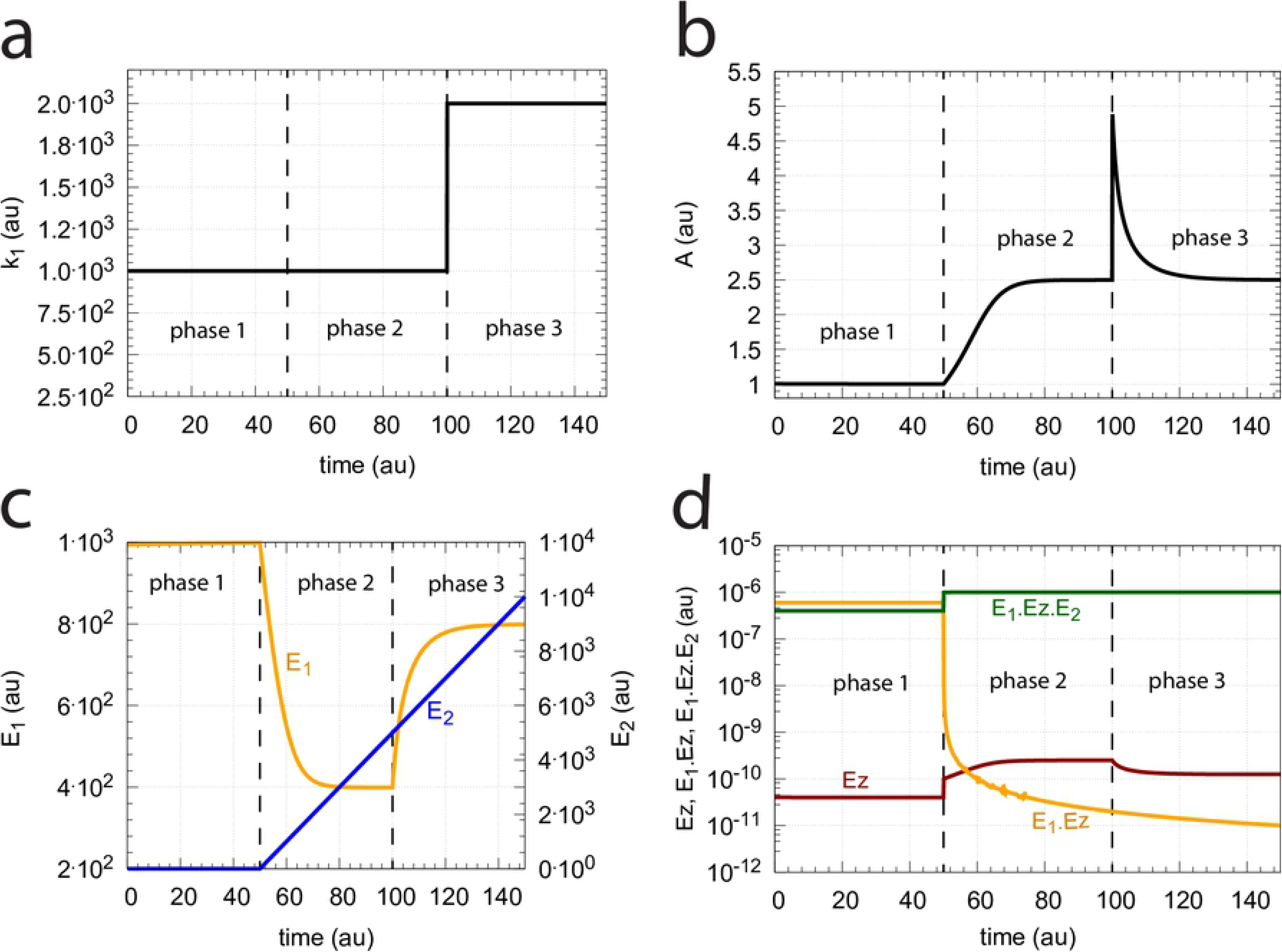
Scheme of the m5 controller when *E*_1_ and *E*_2_ are removed by a compulsory ternary-complex mechanism with *E*_1_ binding first to the free enzyme *Ez*.

The rate equations are:

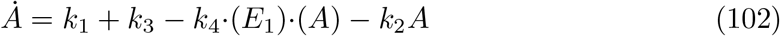

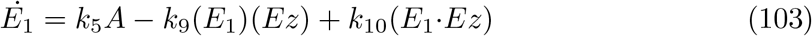

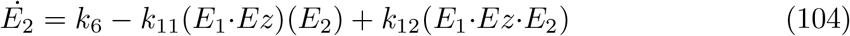

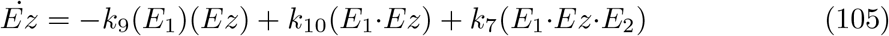

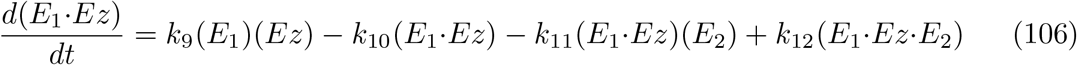

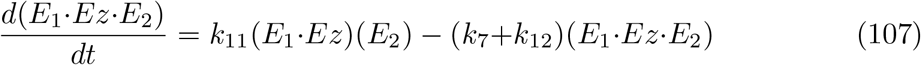

The reaction velocity producing *P* and recycling *Ez* is:

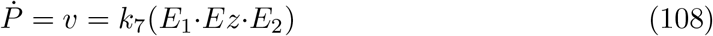

The conditions for the set-point of the dual-E controller for this mechanism are the same as for the random-order ternary-complex case (Eq 100), i.e.

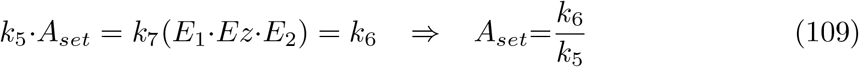

As shown for the random-order ternary-complex case (Eq 100), also for this compulsory ternary-complex mechanism dual-E control requires that

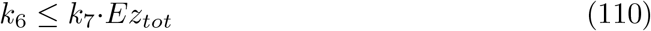

In case *k*_6_>*k*_7_·*Ez_tot_* the feedback switches to single-E control, with *E*_2_ showing wind-up. The set-point switching and *E*_2_ wind-up is illustrated in Fig 36. There, during phase 1, *A* is under dual-E control with set-point of 1.0, where *k*_5_=*k*_6_=40.0 at perturbation *k*_1_=1000.0 (Fig 36a). At the beginning of phase 2 *k*_6_ is increased to 200.0, which leads to a change in the set-point of *A* to 2.5 (=*k*_7_·*Ez_tot_/k*_5_, analogous to Eq 99, Fig 36b) and to wind-up of *E*_2_ (Fig 36c). In phase 3 *k*_1_ is increased to 2000.0 showing that the single-E controller defends its set-point. During single-E control the *E*_1_*Ez* enzyme species rapidly depletes (Fig 36d) and the concentration of the ternary-complex *E*_1_·*Ez*·*E*_2_ becomes practically equal to the total enzyme concentration *Ez_tot_*.

**Fig 36.**
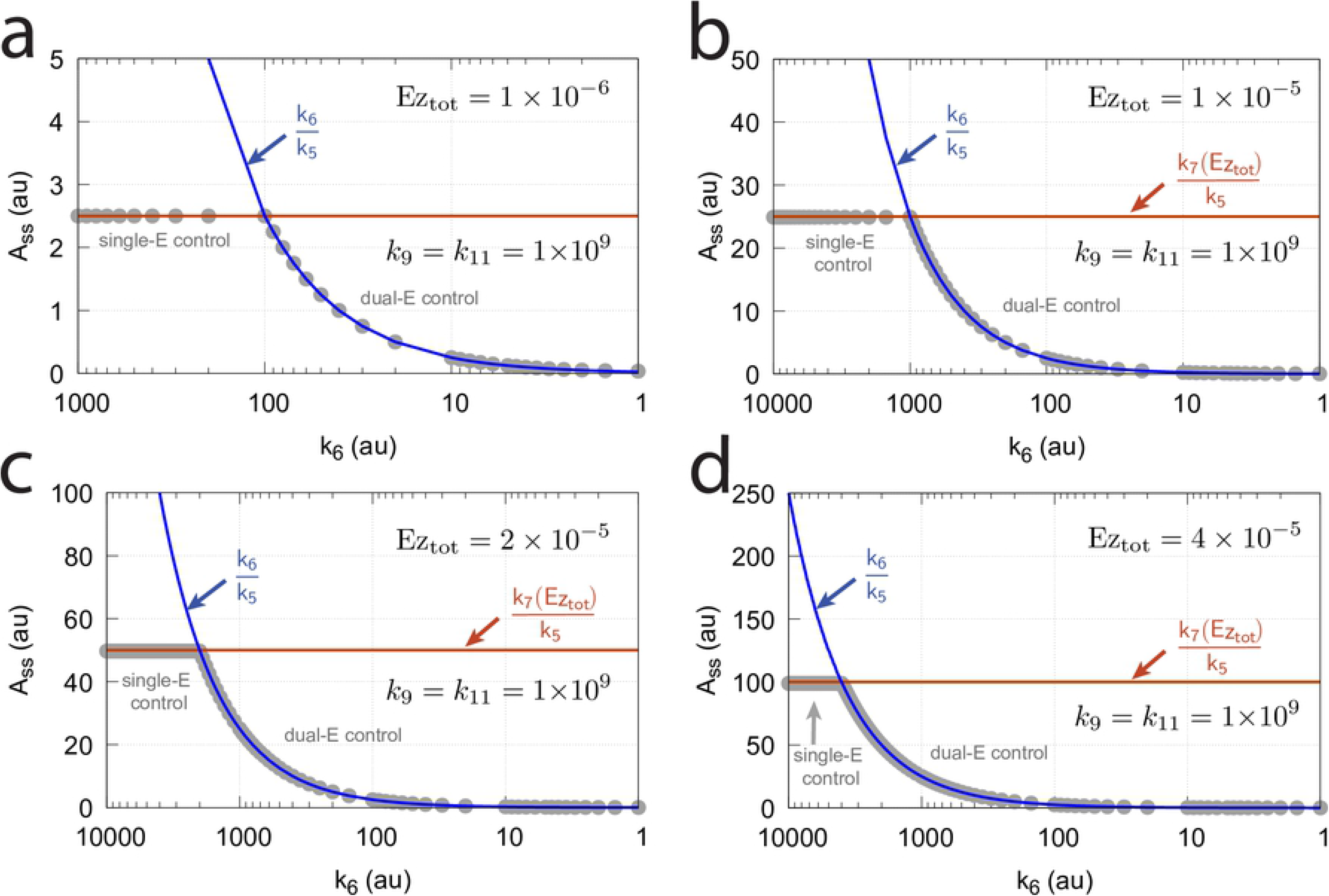
Switch from dual-E control to single-E in the compulsory ternary-complex mechanism of motif 5 when *E*_1_ binds first to free *Ez* (Fig 35). (a) Perturbation *k*_1_ as a function of time. (b) Change of the controlled variable *A*’s concentration as a function of time. Phase 1: dual-E control; phases 2 and 3: single-E control. (c) Concentration of *E*_1_ and *E*_2_ as a function of time. (d) Concentration of the enzymatic species *Ez*, *E*_1_·*Ez*, and *E*_1_·*Ez*·*E*_2_ as a function of time. Rate constants: *k*_1_=1000.0 (phases 1 and 2), *k*_1_=2000.0 (phase 3), *k*_2_=1.0, *k*_3_=0.0, *k*_4_=1.0, *k*_5_=40.0, *k*_6_=40.0 (phase 1), *k*_6_=200.0 (phases 2 and 3) *k*_7_=1×10^8^, *k*_9_=*k*_11_=1×10^9^, *k*_10_=*k*_12_=1×10^3^. Initial concentrations: A_0_=1.0, E_1,0_=993.4, E_2,0_=6.67×10^−2^, Ez_0_=4.02×10^−11^, (E_1_·Ez)_0_=5.999×10^−7^, and (E_1_·Ez·E_2_)_0_=4.00×10^−7^.

Fig 37 shows the switching between dual-E and single-E control as a function of *k*_6_ at four different total enzyme concentrations. Clearly, as previously observed for the other mechanisms, an increase in the enzyme concentration leads to an extended range upon which the dual-E controller is able to act. In addition, the *k*_6_ switch-point for the transition to dual-E control occurs at higher values as total enzyme concentration increases. The set-point for the single-E controller increases accordingly.

**Fig 37.**
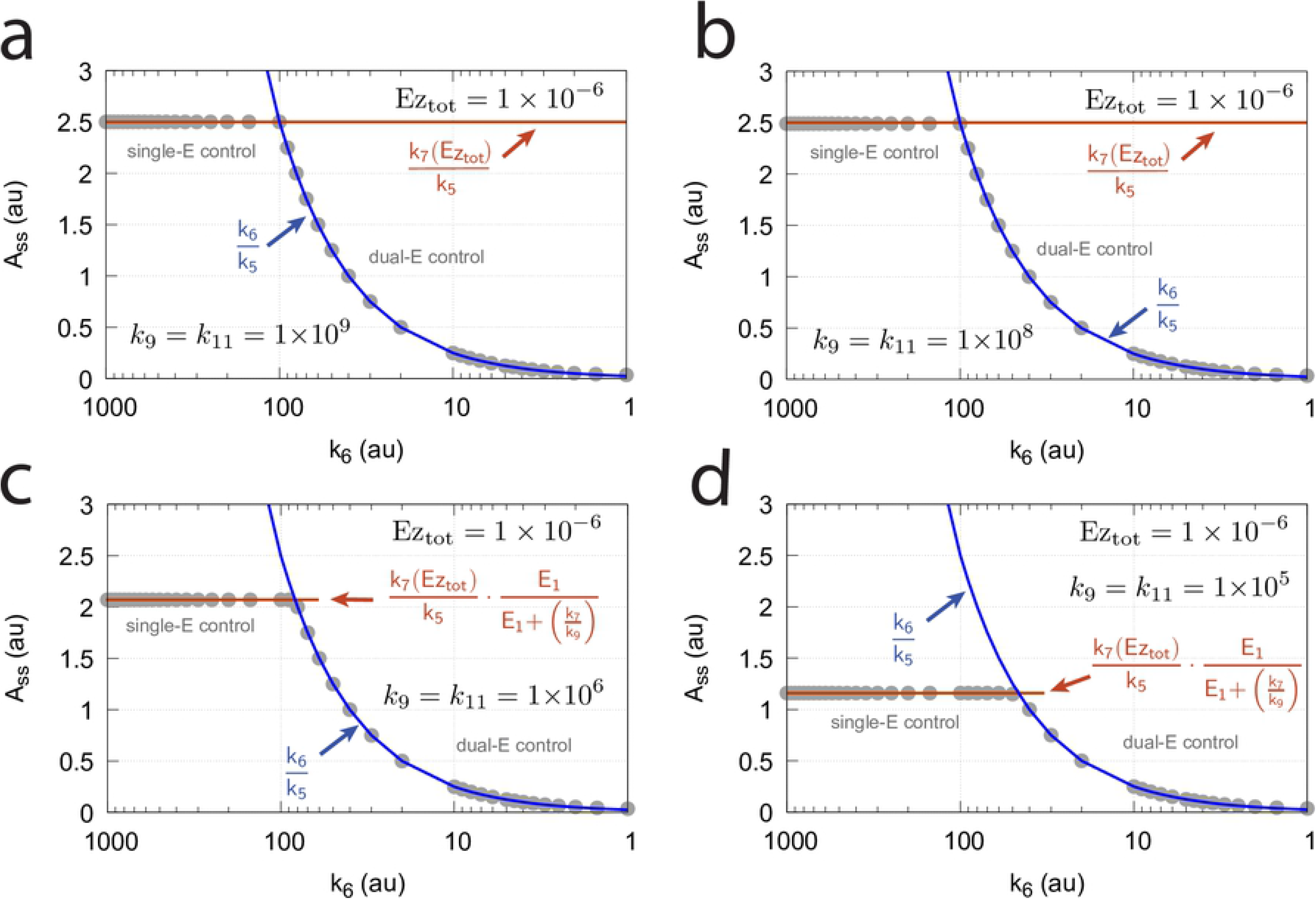
Switch between dual-E and single-E control in the m5 compulsory-order ternary-complex mechanisms (*E*_1_ binding first to *Ez*) as a function of *k*_6_ and total enzyme concentration *Ez_tot_*. (a) *Ez_tot_*=1×10^−6^. (b) *Ez_tot_*=1×10^−5^; (c) *Ez_tot_*=2×10^−5^; (d) *Ez_tot_*=4×10^−5^. Set-points for dual-E and single-E control are indicated in blue and red, respectively. Numerical values are shown as gray filled dots. Rate constants: *k*_1_=1000.0, *k*_2_=1.0, *k*_3_=0.0, *k*_4_=1.0, *k*_5_=40.0, *k*_6_ variable, *k*_7_=1×10^8^, *k*_9_=*k*_11_=1×10^9^, *k*_10_=*k*_12_=1×10^3^. Initial concentrations: A_0_=1.0, E_1,0_=993.4, E_2,0_=6.67×10^−2^ Panel (a): Ez_0_=1×10^−6^, (E_1_·Ez)_0_=0, and (E_1_·Ez·E_2_)_0_=0. Panel (b): Ez_0_=1×10^−5^, (E_1_·Ez)_0_=0, and (E_1_·Ez·E_2_)_0_=0. Panel (c): Ez_0_=2×10^−5^, (E_1_·Ez)_0_=0, and (E_1_·Ez·E_2_)_0_=0. Panel (d): Ez_0_=4×10^−5^, (E_1_·Ez)_0_=0, and (E_1_·Ez·E_2_)_0_=0. *A_ss_* values were taken after a simulation time of 20000 time units.

The velocity how fast P is produced by this mechanism can be expressed analytically using the King-Altman method [22]. The King-Altman treatment leads to

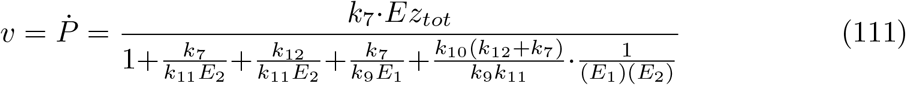

Eq 111 shows that when *k*_9_ and *k*_11_ are much larger than *k*_7_, *k*_10_, and *k*_12_ the velocity becomes zero-order with respect to *E*_1_ and *E*_2_ such that *v* = *k*_7_*Ez_tot_*.

However, when *k*_9_ and *k*_11_ become equal or lower than *k*_7_, *k*_10_, and *k*_12_ the zero-order condition with respect to *E*_1_ and *E*_2_ does no longer hold. In such a case, and when the mechanism shows single-E control, the wind-up of *E*_2_ makes the *E*_2_ terms in Eq 111 disappear, such that at steady state, we have

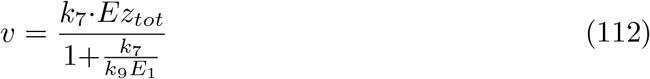

The condition *k*_5_·*A_ss_* = *v* defines the steady-state of *A* at single-E (single-E) control, i.e.

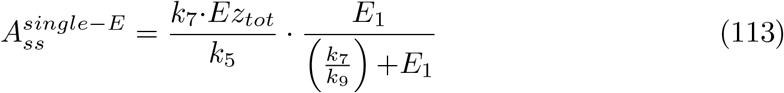

The influence of decreased *k*_9_ and *k*_11_ values in this mechanism is shown in Fig 38. At single-E control, Eq 113 shows excellent agreement with the numerical steady-state values of *A*.

**Fig 38.**
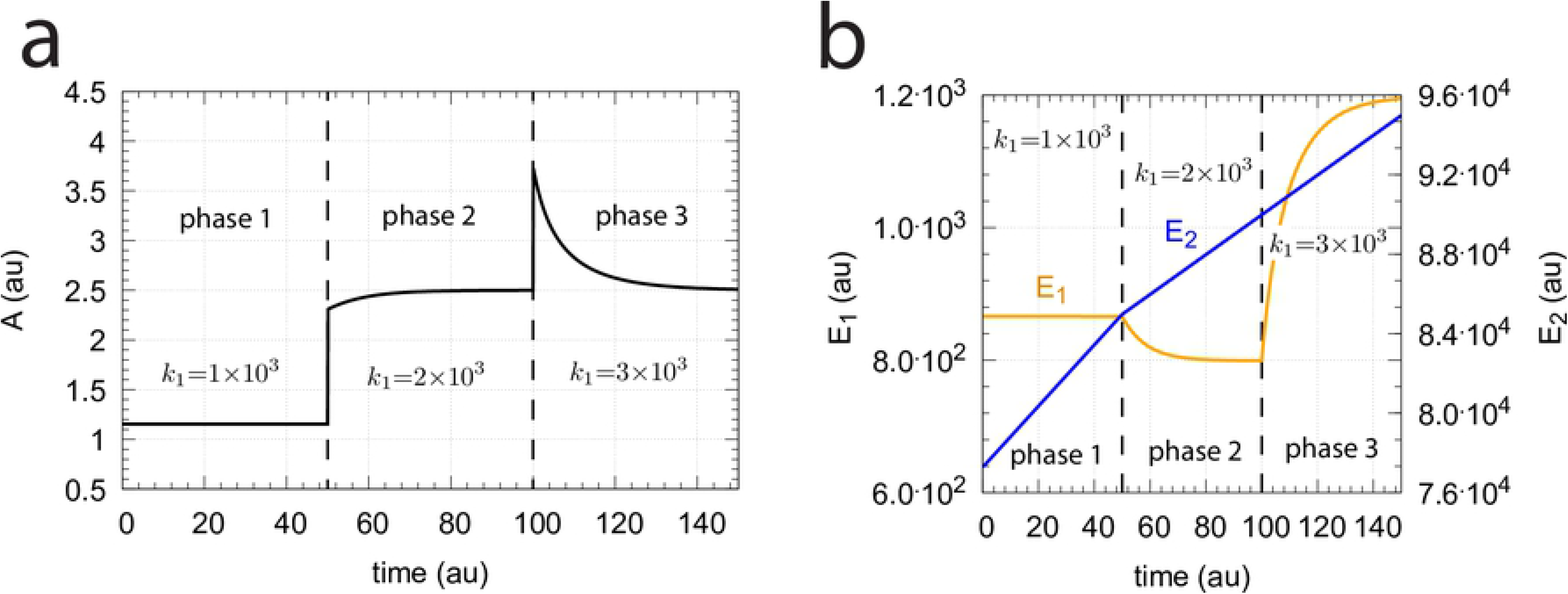
Switch between dual-E and single-E control in the m5 compulsory-order ternary-complex mechanisms (*E*_1_ binding first to *Ez*) as a function of *k*_6_, *k*_9_, and *k*_11_. The total enzyme concentration is 1×10^−6^ and constant. (a) *k*_9_=*k*_11_=1×10^9^. *A_set_* of the single-E controller is 2.5=*k*_7_·*Ez_tot_/k*_5_. (b) *k*_9_=*k*_11_=1×10^8^. Also in this case *A_set_* of the single-E controller is still close to 2.5. (c) *k*_9_=*k*_11_=1×10^6^. *v* (*Ṗ*) is no longer zero-order but is described by Eq 112, and *A_ss_* of the single-E controller is described by Eq. 113 with *E*_1_=4.82×10^2^ and 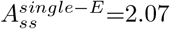. (d) *k*_9_=*k*_11_=1×10^5^. At single-E control conditions we have *E*_1_=4.82×10^2^ and 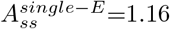 (Eq 113). Other rate constant values and initial concentrations as for Fig 37a.

Note, however, that *A_ss_* will depend on the perturbation *k*_1_. With increasing *k*_1_ values (at single-E control) *E*_1_ will increase such that

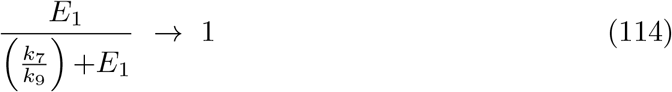

and 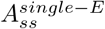 (Eq 113) will move towards the *A_set_* value at zero-order conditions.

Fig 39 shows that increasing *k*_1_ perturbations move *A_ss_* towards the set-point of the single-E controller, as with perturbation-induced increases of the controller variable *E*_1_ the factor *E*1/((*k*_7_/*k*_9_)+*E*_1_) in Eq 114 is getting close to 1.

**Fig 39.**
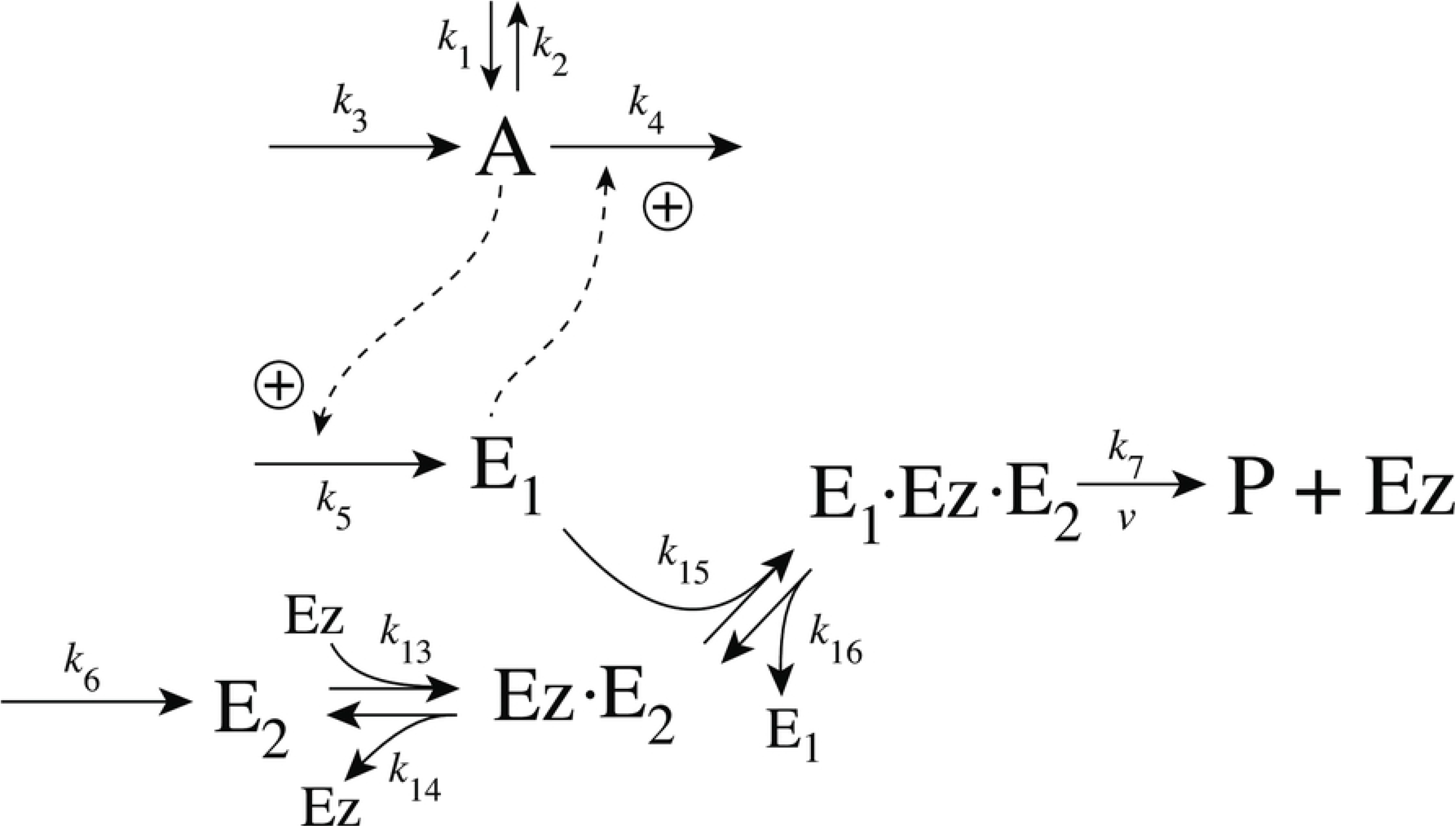
Under single-E control an increased perturbation *k*_1_ moves *A_ss_* in the m5 compulsory-order ternary-complex mechanisms (*E*_1_ binding first to *Ez*) towards *A_set_*=*k*_7_·*Ez_tot_/k*_5_. (a) Phase 1: the system is that from Fig 38d with *k*_6_=200 and *k*_1_=1000.0. In phases 2 and 3 *k*_1_ is stepwise increased to respectively 2000.0 and 3000.0. In phases 2 and 3 *A* is moved to *A_set_*=*k*_7_·*Ez_tot_/k*_5_=2.5. Rate constant values as in Fig 38d. (b) Corresponding changes in *E*_1_ and *E*_2_. Note the wind-up of *E*_2_ and that only *E*_1_ is the controller species. Initial concentrations: A_0_=1.153, E_1,0_=866.2, E_2,0_=7.728×10^6^, Ez_0_=4.027×10^−11^, (E_1_·Ez)_0_=5.999×10^−7^, and (E_1_·Ez·E_2_)_0_=4.000×10^−7^.

#### Motif 5 dual-E controller removing *E*_1_ and *E*_2_ by a compulsory-order ternary-complex mechanism with *E*_2_ binding first to *Ez*

The scheme of this mechanism is shown in Fig 40.

**Fig 40.**
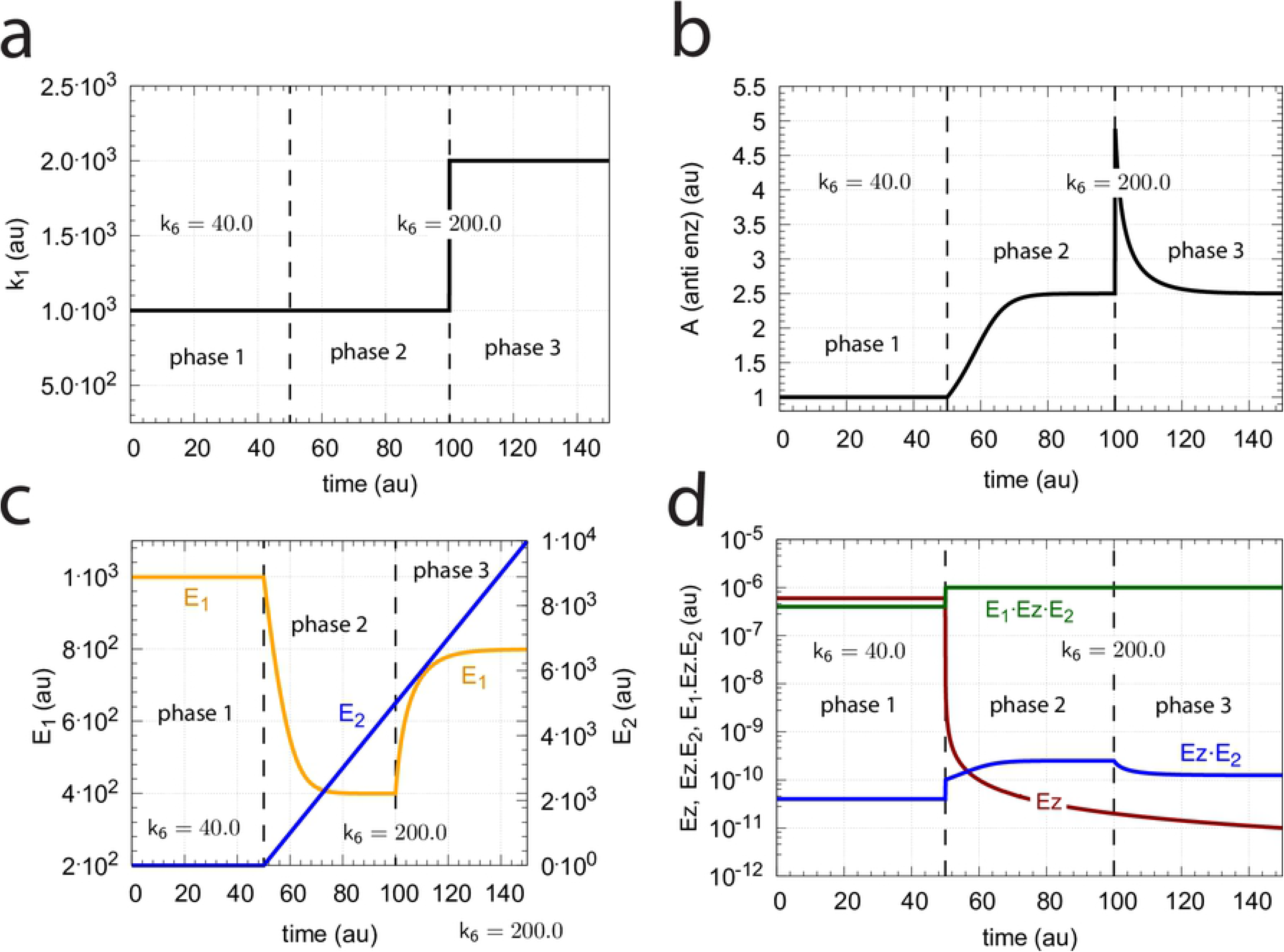
Scheme of the m5 controller when *E*_1_ and *E*_2_ are removed by a compulsory ternary-complex mechanism with *E*_2_ binding first to *Ez*.

The rate equations are:

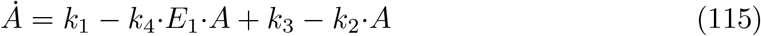

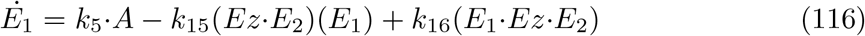

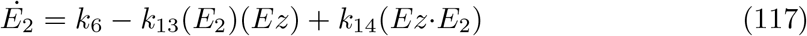

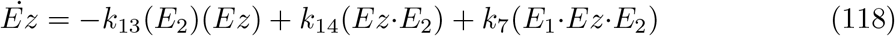

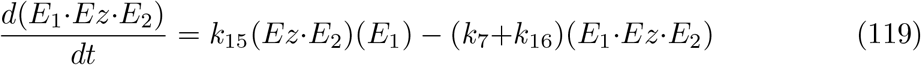

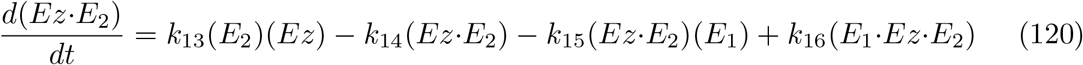

The set-points for dual-E and single-E control are as described in the previous chapter for the m5-compulsory-order ternary-complex mechanism when *E*_1_ binds first (Eq 41), i.e., 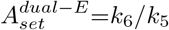 and 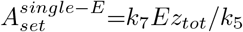.

The velocity *v* how fast P is produced by this mechanism can be expressed analytically using the King-Altman steady-state method

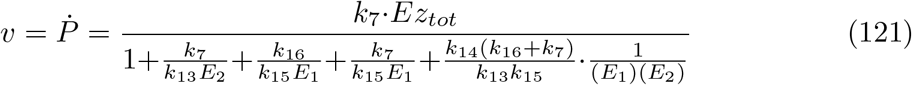

Comparison with numerical results show that Eq 121 gives an excellent description of *v* as a function of *E*_1_ and *E*_2_. Eq 121 also shows that when *k*_13_ and *k*_15_ are much larger than *k*_7_, *k*_14_, and *k*_16_ *v* becomes zero-order with respect to *E*_1_ and *E*_2_ such that *v* = *k*_7_*Ez_tot_*.

This mechanism’s behavior is in many respects identical to the other compulsory-order ternary-complex mechanism of Fig 35. Fig 41 shows as an example a calculation when the change from dual-E to single-E control occurs in an analogous way to that of the other m5 compulsory-order ternary-complex controller shown in Fig 36. For the same rate constant values and the same perturbation profile (Fig 41a) the behaviors of *A*, *E*_1_, and *E*_2_ are precisely the same when Figs 41b,c are compared with Figs 36b,c. Interestingly, *Ez* in this controller (Fig 41d) has now taken the role of *E*_1_·*Ez* in the other controller (Fig 36d), while the function/concentration profile of *Ez* in Fig 36d is identical to that of *Ez*·*E*_2_ in Fig 41d.

**Fig 41.**
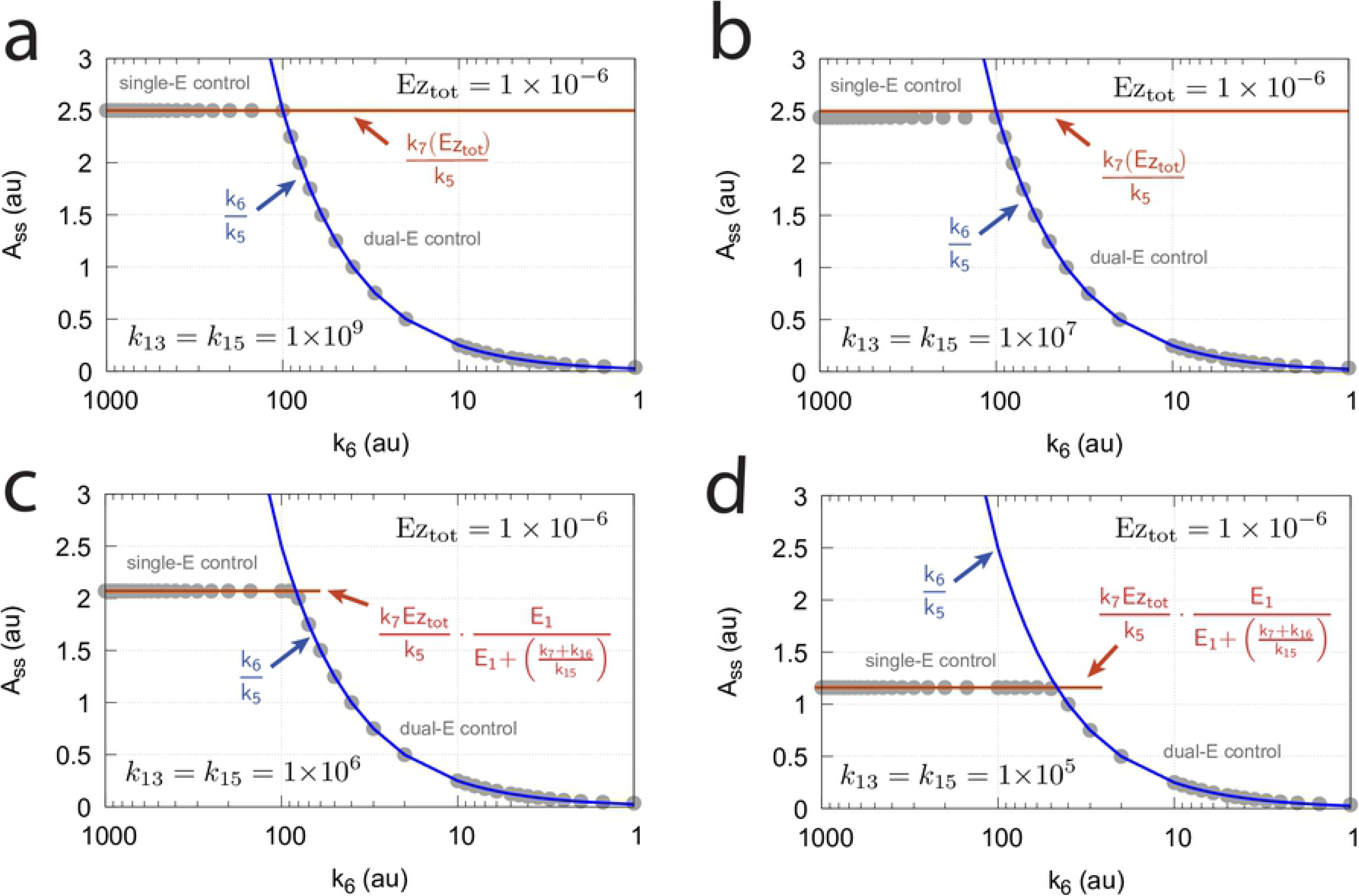
Switch from dual-E to single-E control by increase of *k*_6_ in the compulsory ternary-complex mechanism of motif 5 when *E*_2_ binds first to free *Ez* (Fig 40). An increase of *k*_1_ in phase 3 shows that the set-point of *A* under single-E control is defended. (a) Perturbation *k*_1_ as a function of time. (b) Change of the controlled variable *A*’s concentration as a function of time. Phase 1: dual-E control; phases 2 and 3: single-E control. (c) Concentration of *E*_1_ and *E*_2_ as a function of time. (d) Concentration of the enzymatic species *Ez*, *Ez*·*E*_2_, and *E*_1_·*Ez*·*E*_2_ as a function of time. Rate constants: *k*_1_=1000.0 (phases 1 and 2), *k*_1_=2000.0 (phase 3), *k*_2_=1.0, *k*_3_=0.0, *k*_4_=1.0, *k*_5_=40.0, *k*_6_=40.0 (phase 1), *k*_6_=200.0 (phases 2 and 3) *k*_7_=1×10^8^, *k*_13_=*k*_15_=1×10^9^, *k*_14_=*k*_16_=1×10^3^. Initial concentrations: A_0_=1.0, E_1,0_=9.99×10^2^, E_2,0_=6.67×10^−2^, Ez_0_=5.999×10^−7^, (Ez·E_2_)_0_=4.004×10^−11^, and (E_1_·Ez·E_2_)_0_=4.00×10^−7^.

Fig 42 shows another example of identical behaviors between the two (compulsory-order ternary-complex) m5 controllers when the switching between dual-E and single-E control is investigated as a function of *k*_6_ and when zero-order conditions of *v* with respect to *E*_1_ and *E*_2_ are relaxed. Under single-E control wind-up of *E*_2_ is observed (Fig 41c) such that the expression of *v* (Eq 121) in this case is reduced to

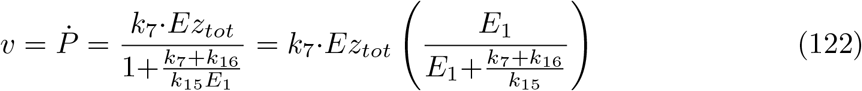

**Fig 42.**
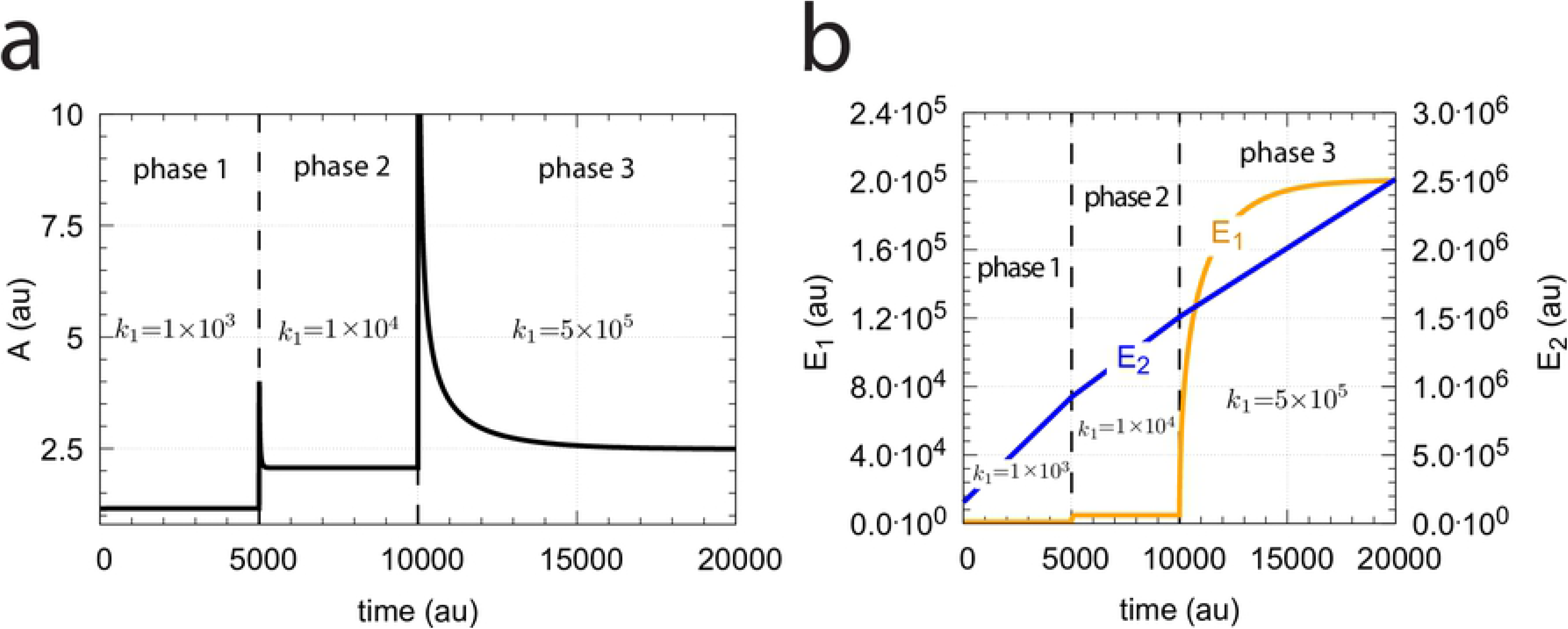
Switch between dual-E and single-E control in the m5 compulsory-order ternary-complex mechanisms (*E*_2_ binding first to *Ez*, Fig 40) as a function of *k*_6_, *k*_13_, and *k*_15_. The total enzyme concentration is 1×10^−6^ and constant. (a) *k*_13_=*k*_15_=1×10^9^. *A_set_* of the single-E controller is 2.5 (=*k*_7_·*Ez_tot_/k*_5_, analogous to Eq 99. (b) *k*_13_=*k*_15_=1×10^8^. Also in this case *A_set_* of the single-E controller is still close to 2.5. (c) *k*_9_=*k*_11_=1×10^6^. *v* (*Ṗ*) is no longer zero-order with respect to *E*_1_ and *E*_2_, but is described by Eq 123, and *A_ss_* of the single-E controller is described by Eq. 124 with *E*_1*,ss*_=4.82×10^2^ and 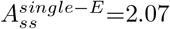. (d) *k*_9_=*k*_11_=1×10^5^. At single-E control conditions we have *E*_1,*ss*_=8.63×10^2^ and 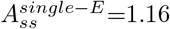 (Eq 124). Other rate parameters as for Fig 37a.

For a given perturbation *k*_1_ the steady states in *A* and *E*_1_ satisfy, under single-E control, the condition

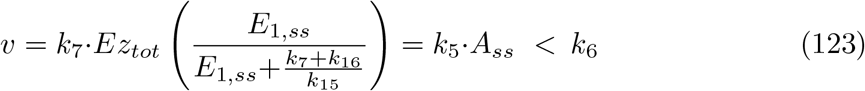

which results in the steady state for *A*:

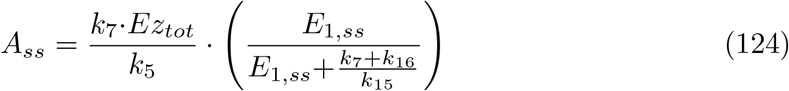

The switch between single-E and dual-E control occurs at

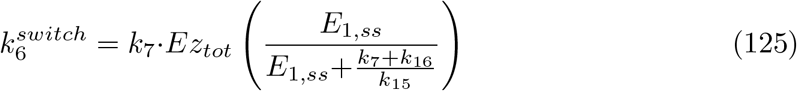

where 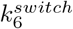 is the smallest *k*_6_ value which is equal to *v* from Eq 123. In the case 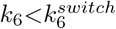 the controller is in dual-E control mode with 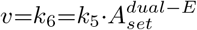.

As already addressed above in Fig 39, a typical property of m5 single-E control is that with increasing perturbation strength the controller species (*E*_1_) increases and *A_ss_* moves towards 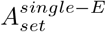. This is also observed for this controller, although higher *k*_1_ values are needed here to reach 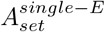. Fig 43 shows the behavior when the Fig 42d parametrization is used with *k*_6_=200.

**Fig 43.**
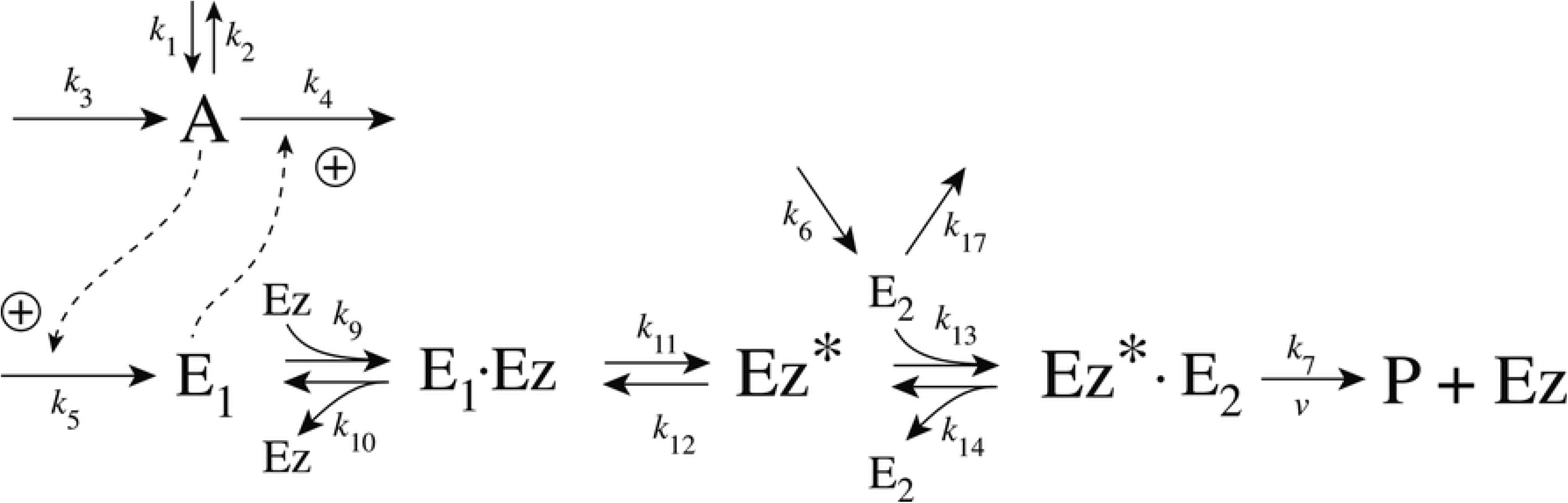
Under single-E control an increased perturbation *k*_1_ moves *A_ss_* in the m5 compulsory-order ternary-complex mechanisms (*E*_1_ binding first to *Ez*) towards *A_set_*=*k*_7_·*Ez_tot_/k*_5_. (a) Phase 1: the system is that from Fig 42d with *k*_6_=200 and *k*_1_=1000.0. In phases 2 and 3 *k*_1_ is stepwise increased to respectively 1×10^4^ and 5×10^5^. In phase 3 *A* is moved close to *A_set_*=*k*_7_·*Ez_tot_/k*_5_=2.5. Rate constant values as in Fig 42d. (b) Corresponding changes in *E*_1_ and *E*_2_. Note the wind-up of *E*_2_ and that only *E*_1_ is the active controller species. Initial concentrations: A_0_=1.156, E_1,0_=866.4, E_2,0_=1.543×10^5^, Ez_0_=2.995×10^−9^, Ez·(E_2_)_0_=5.347×10^−7^, and (E_1_·Ez·E_2_)_0_=4.622×10^−7^.

#### Motif 5 dual-E controller removing *E*_1_ and *E*_2_ by a ping-pong mechanism with *E*_1_ binding first to *Ez*

The scheme of this mechanism is shown in Fig 44.

**Fig 44.**
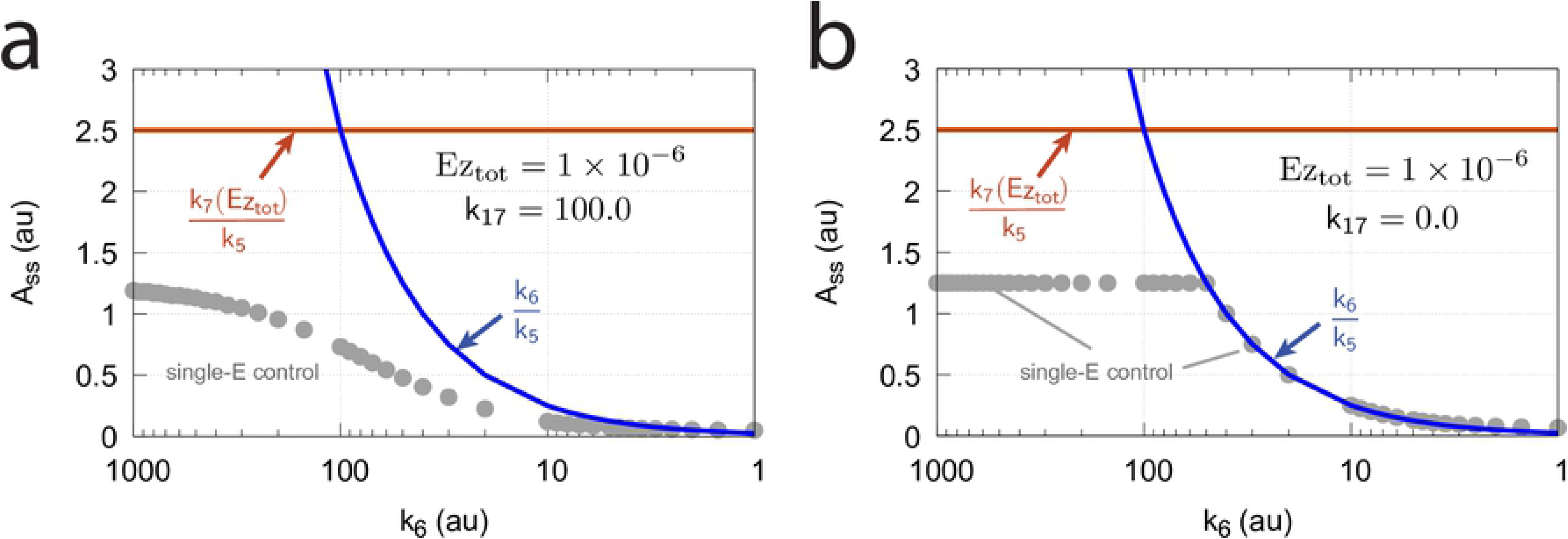
Scheme of the m5 controller when *E*_1_ and *E*_2_ are removed by a ping-pong mechanism with *E*_1_ binding first to the free enzyme *Ez*.

We have included a first-order degradation term of *E*_2_ with rate constant *k*_17_. The reason for this is the observation that for this controller only *E*_1_ acts as a control species while *E*_2_ remains to be constant. To see the influence of *E*_2_ on the set-point the first-order degradation of *E*_2_ is included.

The rate equations are:

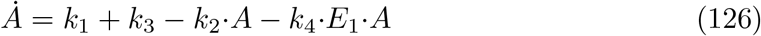

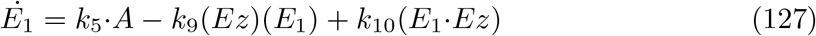

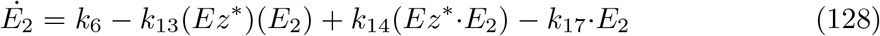

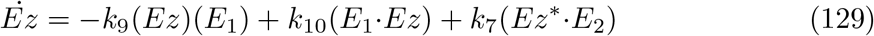

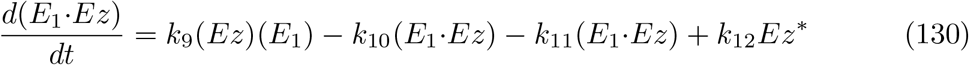

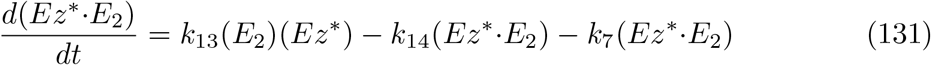

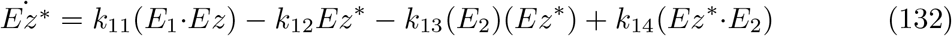

The numerically calculated velocity *v_num_* by which *P* is formed is calculated as

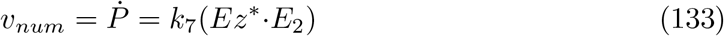

*v_num_* is in excellent agreement when *Ṗ* is calculated by using the steady-state approach with the help of the King-Altman method (see S1 Text).

In this case, *v_ss_* is

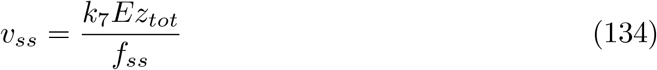

with

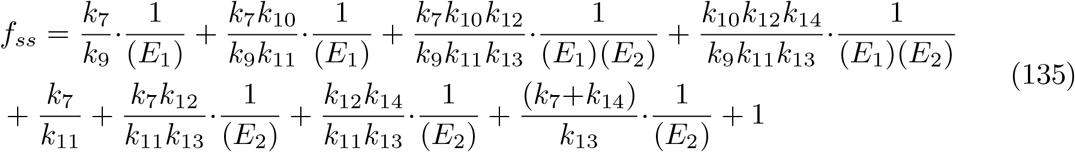

From the rate equation of *E*_2_ (Eq 128) we see that the concentrations of *E*_2_ are related to the concentrations of *Ez*^*^ and *Ez*^*^·*E*_2_. Since *Ez*^*^ and *Ez*^*^·*E*_2_ show constant steady-state values the concentration of *E*_2_ is constant in time, but its value is dependent on the values of the other rate constants.

The relationship

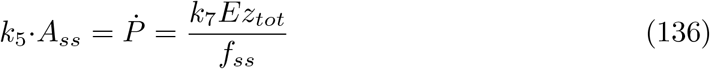

determines the set-point for *A*, *A_set_*, as

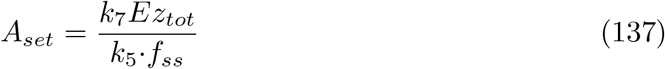

whenever *Ṗ* is constant and independent of the perturbation *k*_1_. Independence of *Ṗ* from *k*_1_ occurs when the terms *α_i_*/(*E*_1_) in the first line of Eq 135 become zero, either by sufficient large *E*_1_ values or/and by the *α_i_*’s being close to zero (large *k*_9_ and *k*_11_ values in comparison to *k*_7_ and *k*_10_). We found robust homeostasis in *A* for a large range of rate constant values. The rate constant values used here have been chosen such that comparisons with the other controllers can be made and for getting controller response times which are not too large.

A striking observation in comparison with the m5-based controllers based on ternary-complex mechanisms is that *E*_2_ has apparently no control function and that the ping-pong mechanism appears to be entirely controlled by *E*_1_, even if *A_ss_*=*A_set_*=*k*_6_/*k*_5_, when *k*_17_=0, described by the set-point condition

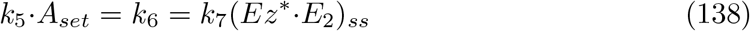

Fig 45 shows steady state values *A_ss_* as a function of *k*_6_ when *k*_17_=100.0 and 0.0, at a total enzyme concentration of 1.0×10^−6^. Each *A_ss_* values represents an actual set-point of *A*, which is defended against step-wise perturbations by *k*_1_.

**Fig 45.**
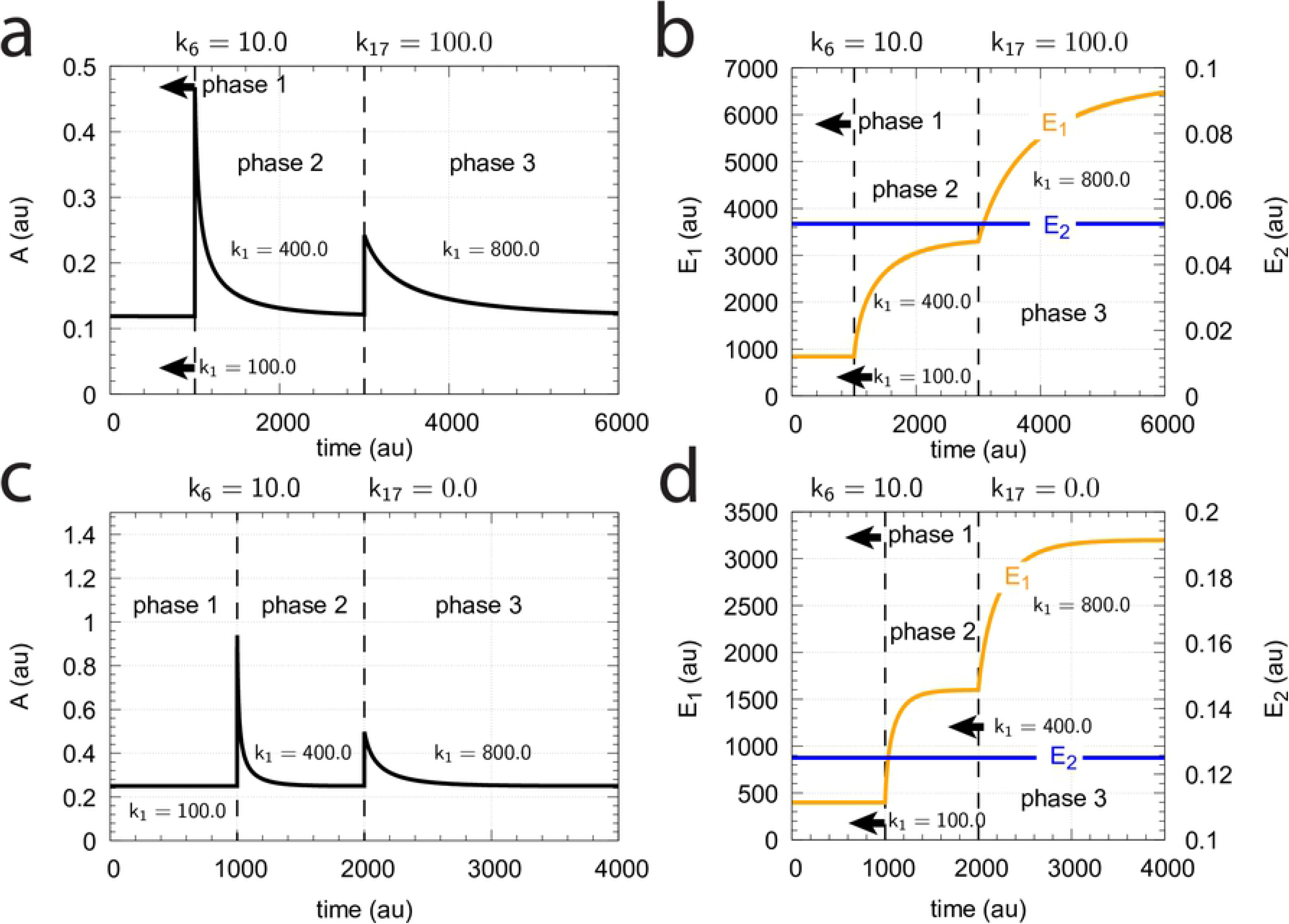
*A_ss_* (=*A_set_*) as a function of *k*_6_ when (a) *k*_17_=100.0 and (b) *k*_17_=0.0. Gray solid points show the numerically calculated values of *A_ss_*, while red and blue curves show the values of *k*_7_(*Ez_tot_*)/*k*_5_ and *k*_6_/*k*_5_, respectively. Other rate constant values: *k*_1_=800.0, *k*_2_=1.0, *k*_3_=0.0, *k*_4_=1.0, *k*_5_=40.0, *k*_7_=1×10^8^, *k*_9_=*k*_11_=*k*_13_=1×10^8^, and *k*_10_=*k*_12_=*k*_14_=1×10^3^. Initial concentrations: A_0_=1.0, E_1,0_=9.9×10^1^, E_2,0_=5.04×10^−1^, Ez_0_=6.03×10^−9^, (E_1_·Ez)_0_=4.97×10^−7^, (Ez^*^·E_2_)_0_=1.0×10^−7^, and 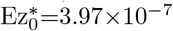. Simulation time: 5000 time units, step-length: 0.01 time units.

Fig 46 shows the homeostatic behavior of *A_ss_* in Fig 45 for *k*_6_=10.0 when *k*_17_=100.0 (panels a and b), or when *k*_17_=0.0 (panels c and d). Note that only *E*_1_ acts as the controller variable while *E*_2_ is constant independent of the values of the perturbation *k*_1_.

**Fig 46.**
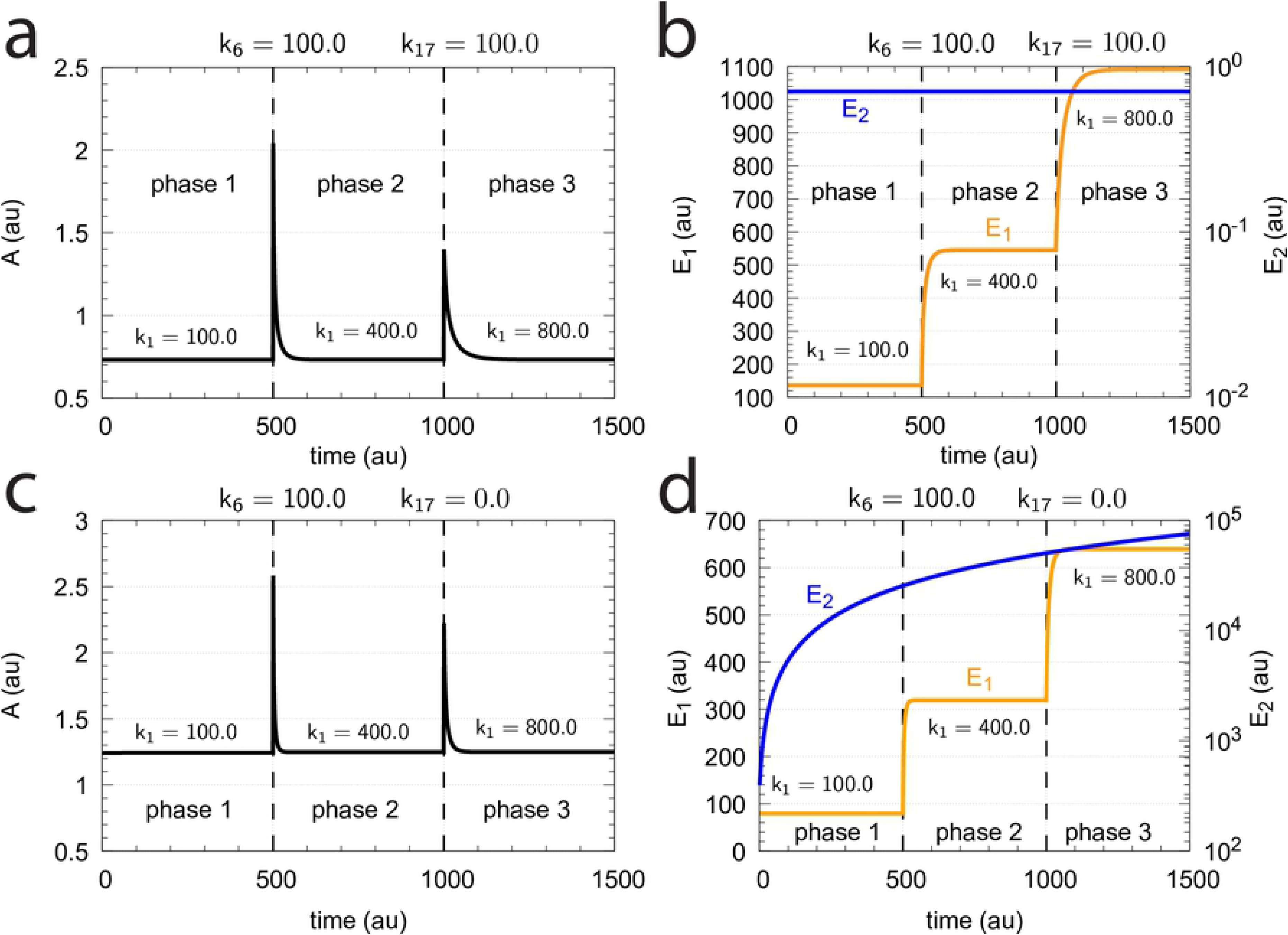
Demonstration of homeostatic behavior in *A* (Fig 45) when *k*_6_=10.0 and *k*_17_=100.0 (panels a and b) or *k*_17_=0.0 (panel c and d). Step-wise perturbations are applied with values *k*_1_=100 (phase 1), *k*_1_=400 (phase 2), and *k*_1_=800 (phase 3). Other rate constants as in Fig 45. Initial concentrations, (a) and (b): A_0_=0.1189, E_1,0_=8.40×10^2^, E_2,0_=5.25×10^−2^, Ez_0_=5.66×10^−11^, (E_1_·Ez)_0_=4.75×10^−8^, (Ez^*^·E_2_)_0_=4.75×10^−8^, and 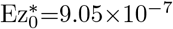. Initial concentrations, (c) and (d): A_0_=0.25, E_1,0_=3.99×10^2^, E_2,0_=1.25×10^−1^, Ez_0_=2.51×10^−10^, (E_1_·Ez)_0_=1.00×10^−7^, (Ez^*^·E_2_)_0_=1.00×10^−7^, and 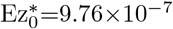.

For *k*_6_=100.0 the controller’s behavior is shown in Fig 47. Also in this case robust homeostasis in *A* is observed due to the high values of *E*_1_ and the constancy in *E*_2_. Due to the large values in both *E*_1_ and *E*_2_ the set-point of the controller is (*k*_17_=0):

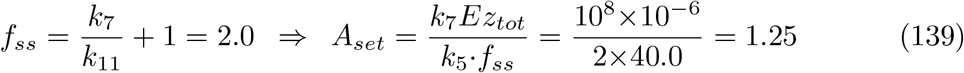

**Fig 47.**
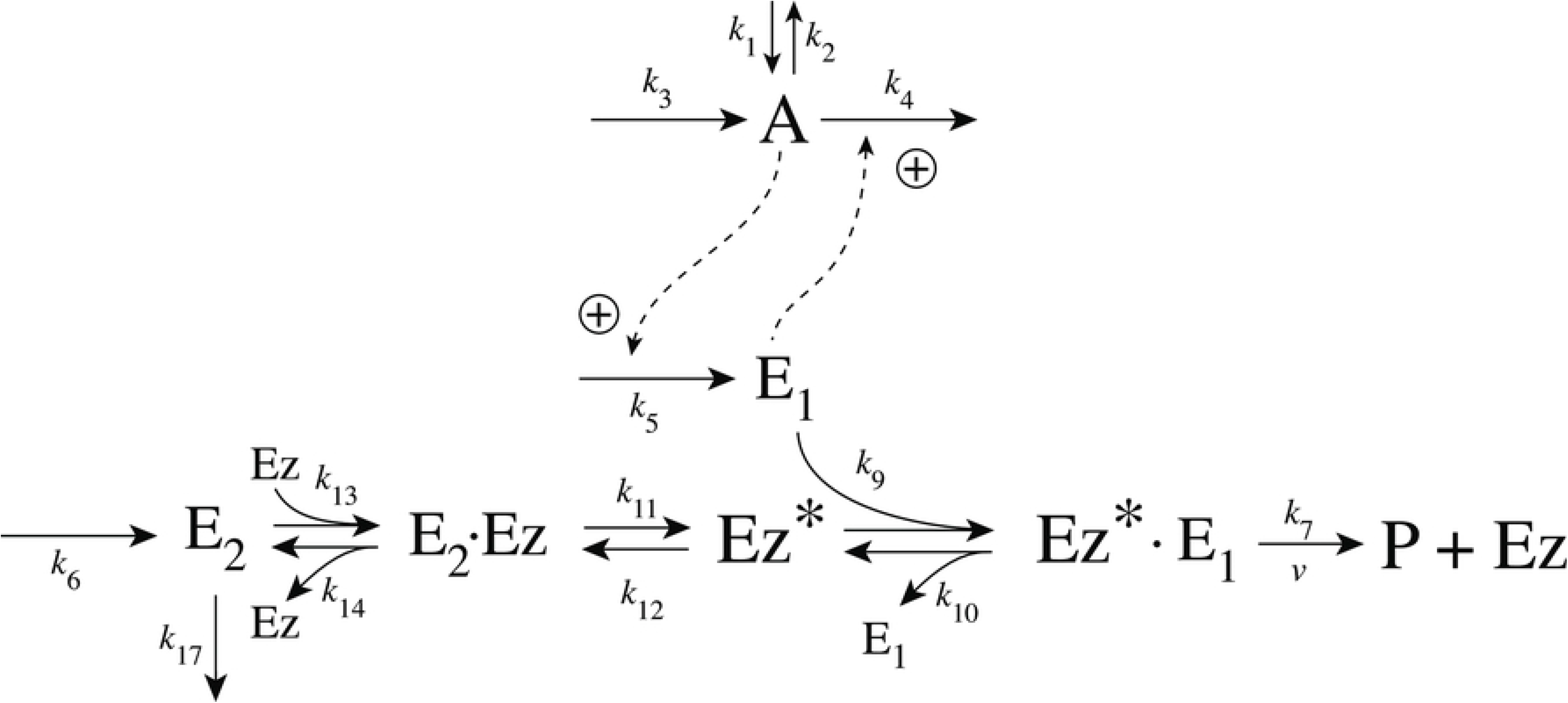
Demonstration of the homeostatic behavior of *A_ss_* in Fig 45 when *k*_6_=100.0 and *k*_17_=100.0 (panels a and b) or *k*_17_=0.0 (panel c and d). Step-wise perturbations are applied with values *k*_1_=100 (phase 1), *k*_1_=400 (phase 2), and *k*_1_=800 (phase 3). Other rate constants as in Fig 45. The linear increase of *E*_2_ is seen as a concave line due to the logarithmic scale of the *E*_2_-axis. Initial concentrations, (a) and (b): A_0_=0.7309, E_1,0_=1.36×10^2^, E_2,0_=7.08×10^−1^, Ez_0_=2.15×10^−9^, (E_1_·Ez)_0_=2.95×10^−7^, (Ez^*^·E_2_)_0_=2.93×10^−7^, and 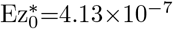. Initial concentrations, (c) and (d): A_0_=1.24, E_1,0_=7.96×10^1^, E_2,0_=3.50×10^−2^, Ez_0_=6.23×10^−9^, (E_1_·Ez)_0_=4.96×10^−7^, (Ez^*^·E_2_)_0_=4.96×10^−7^, and 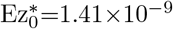.

#### Motif 5 dual-E controller removing *E*_1_ and *E*_2_ by a ping-pong mechanism with *E*_2_ binding first to *Ez*

Fig 48 shows the scheme of this mechanism.

**Fig 48.**
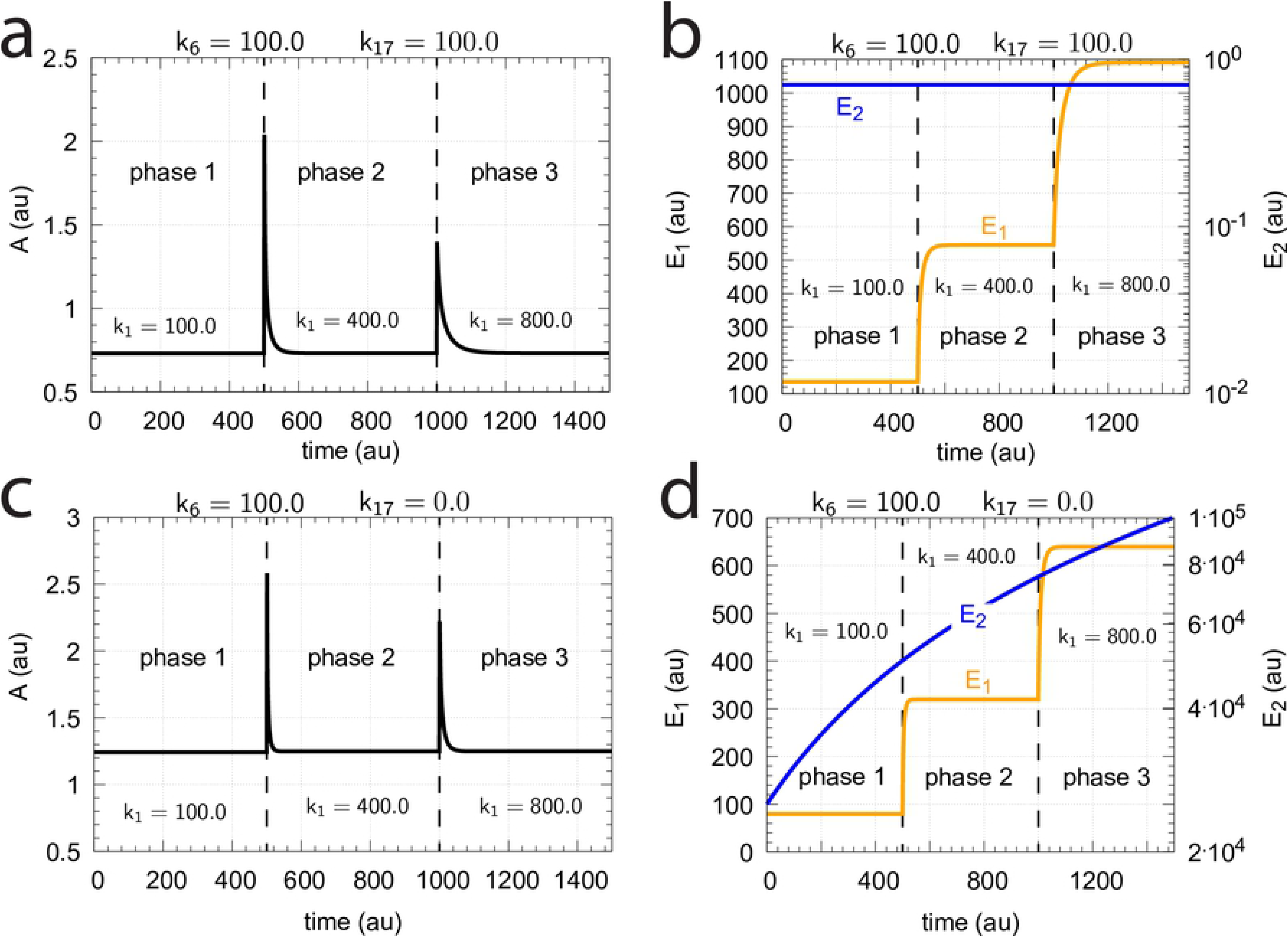
Scheme of the m5 controller when *E*_1_ and *E*_2_ are removed by a ping-pong mechanism with *E*_2_ binding first to the free enzyme *Ez*.

The rate equations are:

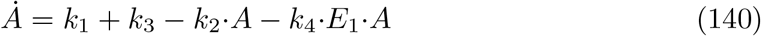

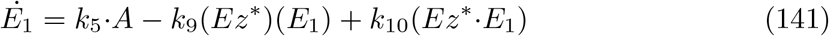

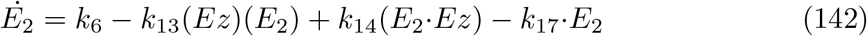

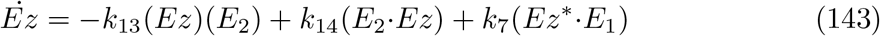

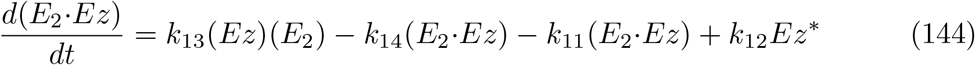

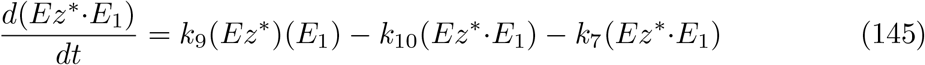

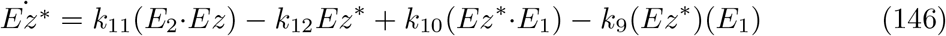

The numerically calculated velocity is calculated as

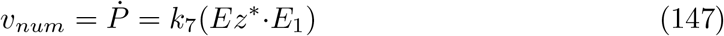

which has been found to be in excellent agreement with the steady-state expression

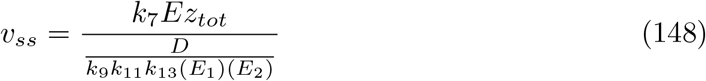

with D being

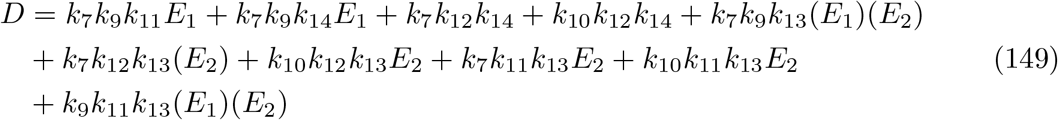

Eq 148 is derived along the same lines as for Eq 134, i.e. by using the King-Altman method (S1 Text).

The set-point *A_set_* for the controller is determined by its steady-state, *A_ss_*, due to the condition

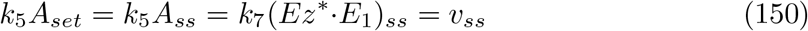

where *v_ss_* is independent of *k*_1_. The maximum velocity is reached for zero-order conditions with respect to *E*_1_ and *E*_2_, and is given by

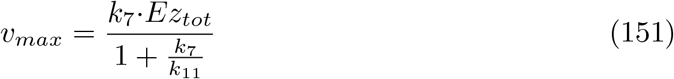

analogous to the condition by Eq 139.

The relationship between *A_set_* and (*Ez*^*^·*E*_1_)_*ss*_ is also independent of *k*_6_. Like for the m5 ping-pong based controller when *E*_1_ binds first to *Ez*, *E*_2_ has also here no control function and remains constant as long as the inflow of *k*_6_ can be compensated by *v*=*Ṗ*. However, when *k*_6_>*v_max_*, and for example *k*_17_=0, *E*_2_ will increase linearly. Fig 49 shows the controller’s behavior for three different *k*_1_ perturbation values analogous as in Fig 47 when *k*_6_=100.0. For this high value of *k*_6_ (> *v_max_* and *k*_17_=0) *E*_2_ shows a (linear) increase.

**Fig 49.**
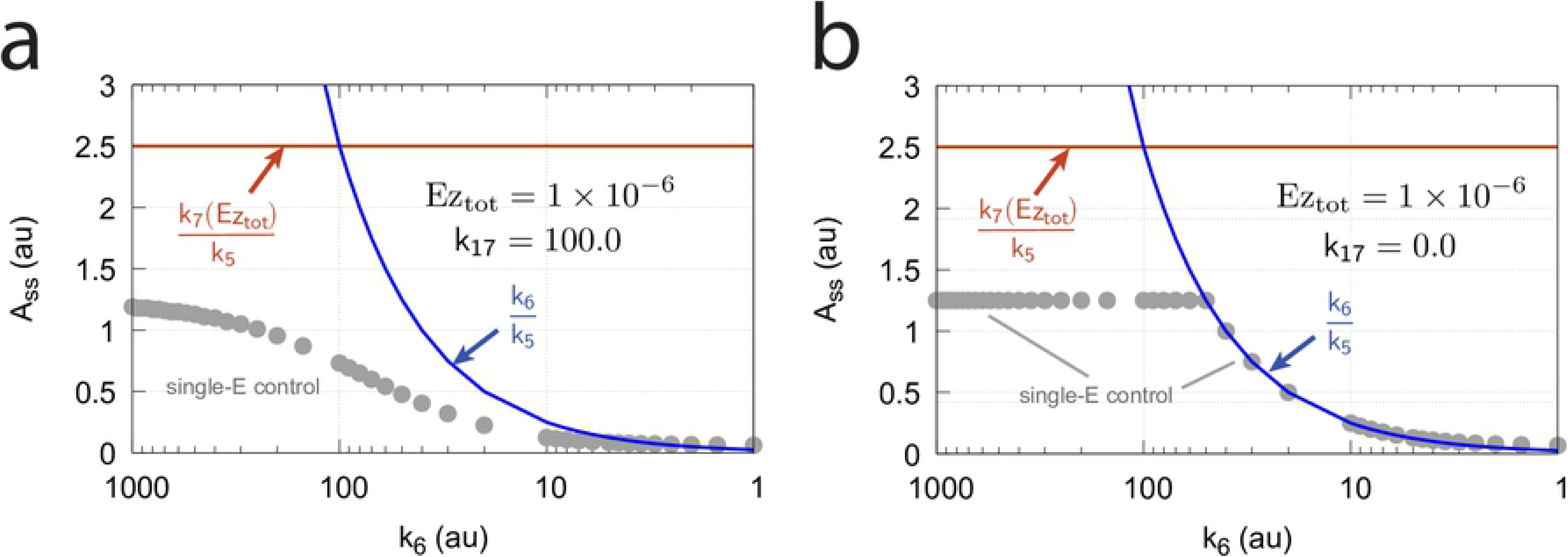
Demonstration of the homeostatic behavior of *A_ss_* (scheme Fig 48) when *k*_6_=100.0 and *k*_17_=100.0 (panels a and b) or *k*_17_=0.0 (panel c and d). The behavior is analogous to that shown in Fig 47. Step-wise perturbations are applied with values *k*_1_=100 (phase 1), *k*_1_=400 (phase 2), and *k*_1_=800 (phase 3). Other rate constants as in Fig 45. The linear increase of *E*_2_ is seen as a concave line due to the logarithmic scale of the *E*_2_-axis. *v_max_*=50 (Eq 151). Initial concentrations, (a) and (b): A_0_=0.7309, E_1,0_=1.36×10^2^, E_2,0_=7.08×10^−1^, Ez_0_=2.15×10^−9^, (E_2_·Ez)_0_=2.924×10^−7^, (Ez^*^·E_1_)_0_=2.924×10^−7^, and 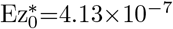. Initial concentrations, (c) and (d): A_0_=1.242, E_1,0_=7.95×10^1^, E_2,0_=2.52×10^4^, Ez_0_=1.97×10^−11^, (E_2_·Ez)_0_=4.97×10^−7^, (Ez^*^·E_1_)_0_=4.97×10^−7^, and 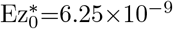.

Fig 50 shows how *A_ss_* (which defines the set-point *A_set_*) depends on *k*_6_. Only when *k*_17_=0 and *k*_6_ < *v_max_* the set-point is defined by *k*_6_/*k*_5_. Despite the formal agreement with the set-point of a dual-E control when comparison is made to the above ternary-complex motif 5 mechanisms, the ping-pong mechanism shows robust single-E control conducted by *E*_1_.

**Fig 50.**
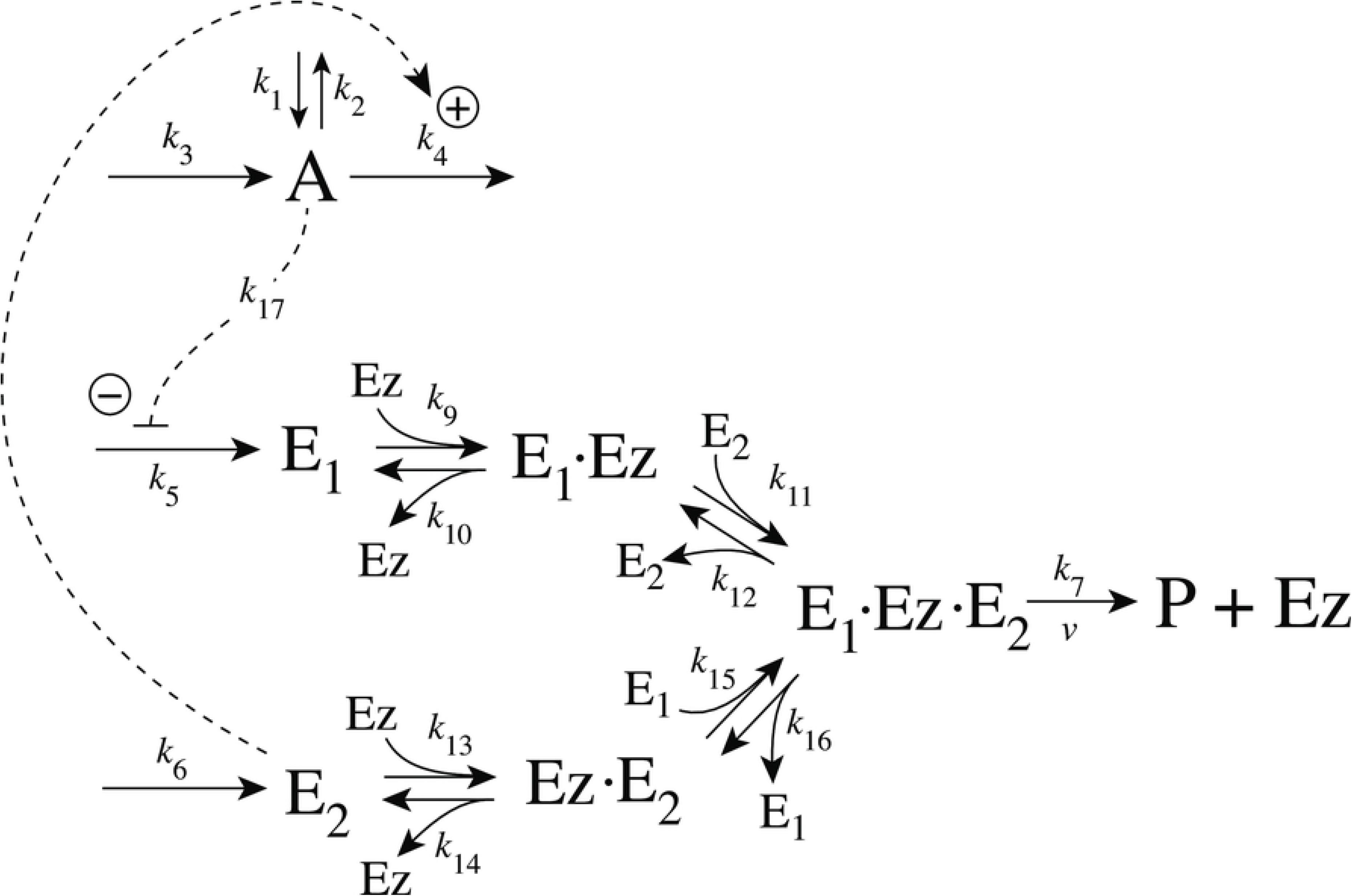
*A_ss_* (=*A_set_*) as a function of *k*_6_ when (a) *k*_17_=100.0 and (b) *k*_17_=0.0. Gray solid points show the numerically calculated values of *A_ss_*, while red and blue curves show the values of *k*_7_(*Ez_tot_*)/*k*_5_ and *k*_6_/*k*_5_, respectively. Other rate constant values: *k*_1_=800.0, *k*_2_=1.0, *k*_3_=0.0, *k*_4_=1.0, *k*_5_=40.0, *k*_7_=1×10^8^, *k*_9_=*k*_11_=*k*_13_=1×10^8^, and *k*_10_=*k*_12_=*k*_14_=1×10^3^. Initial concentrations: A_0_=1.0, E_1,0_=9.9×10^1^, E_2,0_=5.04×10^−1^, Ez_0_=6.03×10^−9^, (E_2_·Ez)_0_=4.97×10^−7^, (Ez^*^·E_1_)_0_=1.0×10^−7^, and 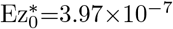. Simulation time: 3000 time units, step-length: 0.01 time units.

### Controllers based on motif 7

#### Motif 7 dual-E controller removing *E*_1_ and *E*_2_ by a random-order ternary-complex mechanism

Fig 51 shows the reaction scheme when in a m7 controller configuration *E*_1_ and *E*_2_ are removed by an enzymatic random-order ternary-complex mechanism.

**Fig 51.**
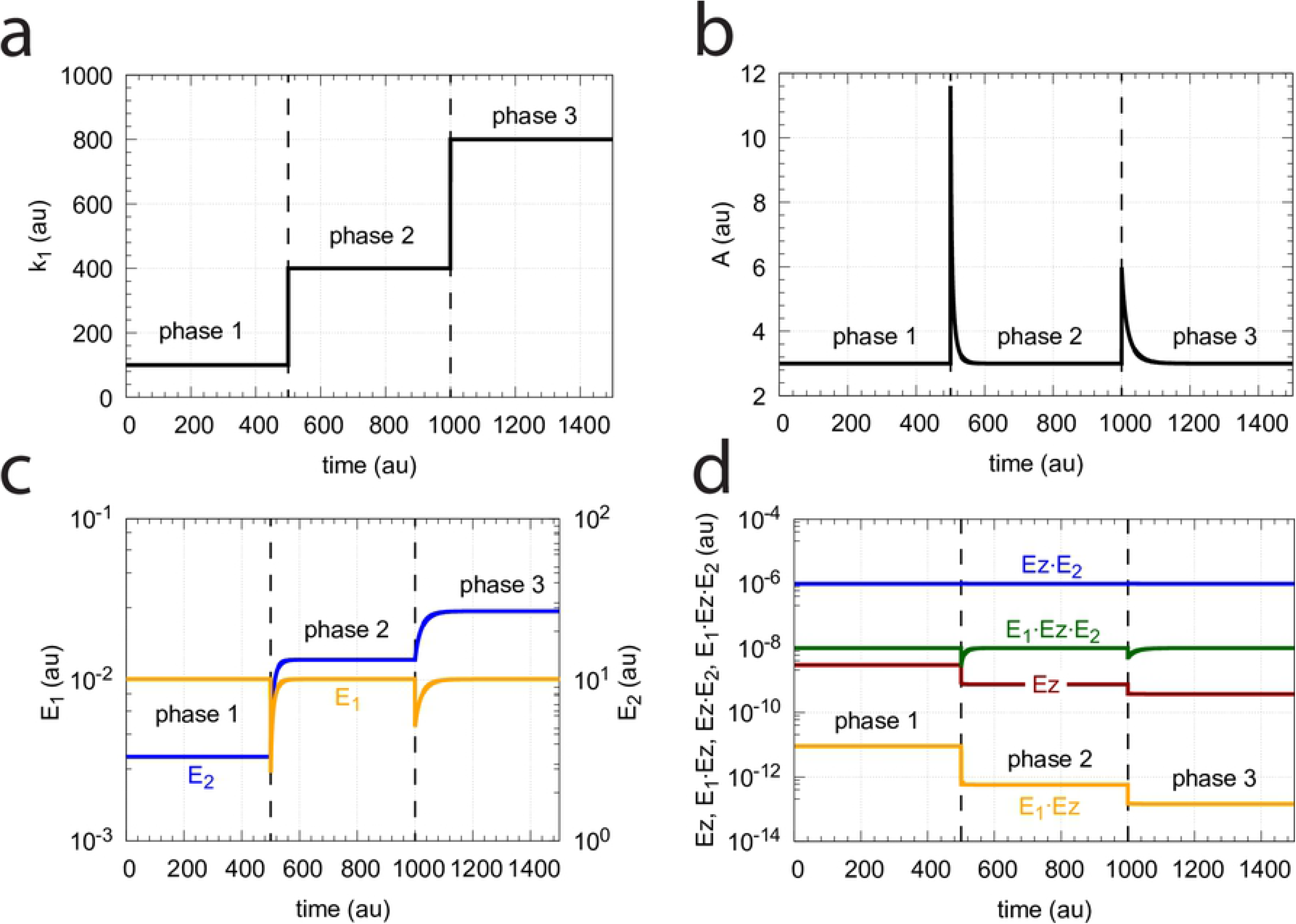
Reaction scheme of the m7-type of controller when *E*_1_ and *E*_2_ are removed by enzyme *Ez* with a random-order ternary-complex mechanism.

The rate equations are:

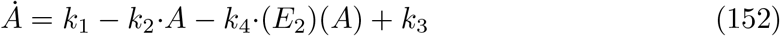

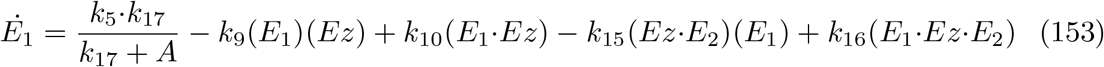

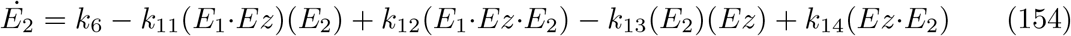

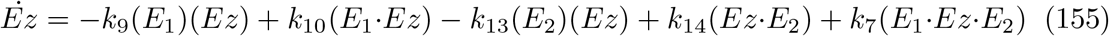

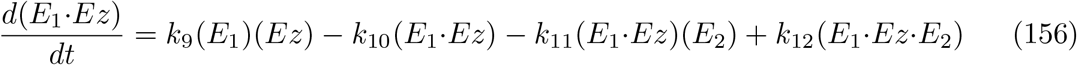

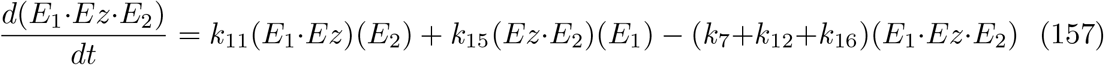

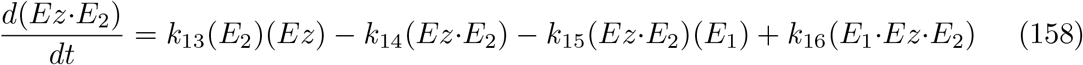

Under dual-E conditions the steady state concentration of *A* is determined by setting inflow rates *k*_6_ and *j*_5_=*k*_5_*k*_17_/(*k*_17_ + *A_ss_*) equal to the outflow rate *v*=*k*_7_(*E*_1_·*Ez*·*E*_2_), i.e.,

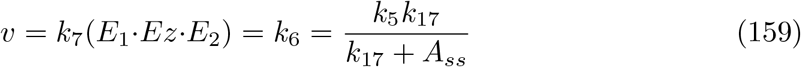

Solving for *A_ss_*, which is equal to *A_set_*, gives

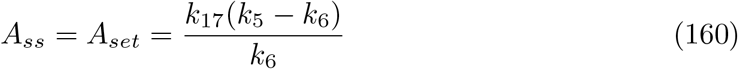

Fig 52 illustrates the controller’s homeostatic behavior, following Eq 160, for step-wise perturbations in *k*_1_. The controller operates by increasing the controller variable *E*_2_, which activates the compensatory flux *k*_4_·(*E*_2_)(*A*). Although *E*_1_ undergoes an excursion during the perturbation, its steady-state value remains unchanged at the different *k*_1_ values.

**Fig 52.**
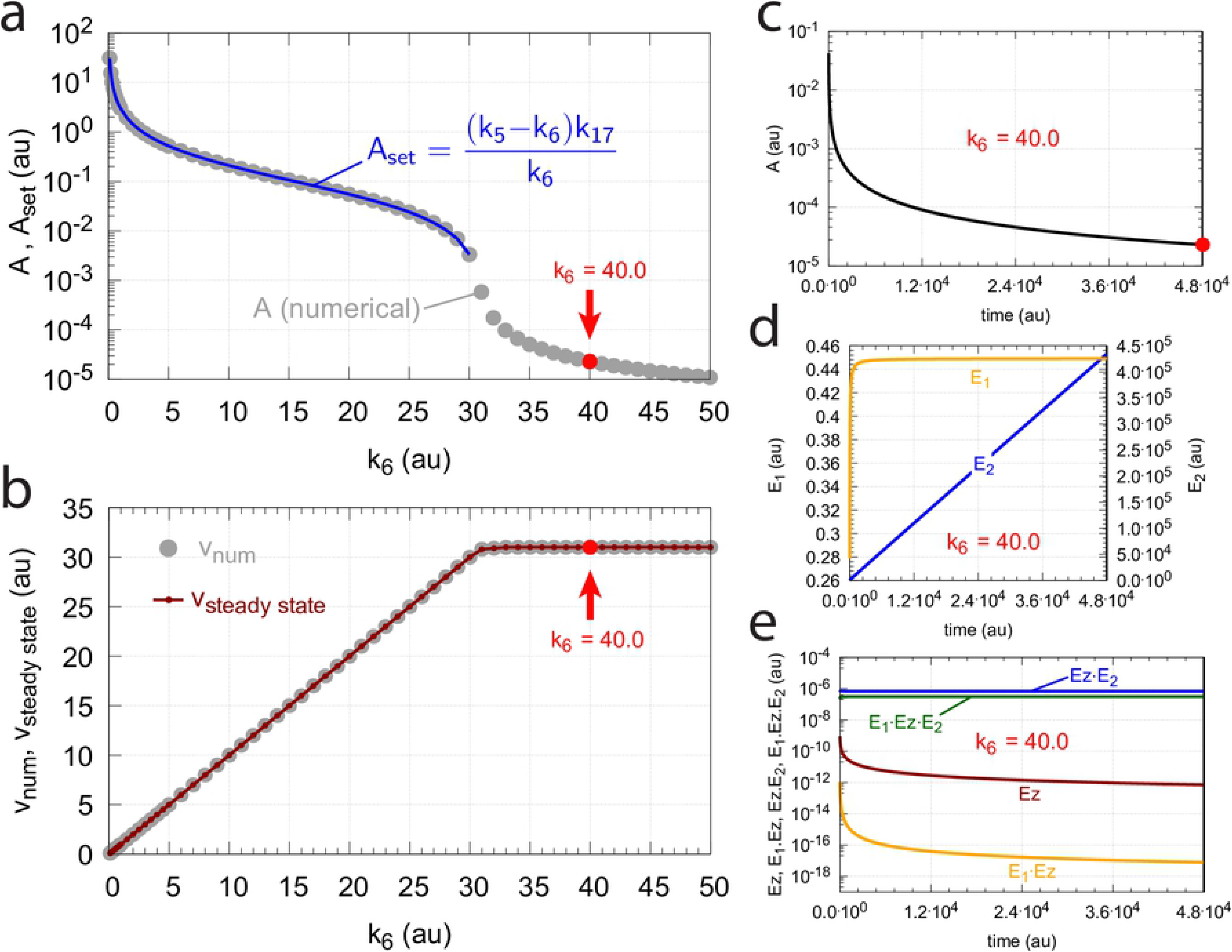
Homeostatic behavior towards step-wise perturbations of *k*_1_ in the scheme of Fig 51. (a) stepwise changes of *k*_1_, (b) homeostatic control of *A*, (c) Variation of controller variables *E*_1_ and *E* − 2, (d) changes in the enzymatic species *Ez*, *E*_1_·*Ez*, *Ez*·*E*_2_, and *E*_1_·*Ez*·*E*_2_. Rate constants: *k*_1_=100.0 in phase 1, 400.0 in phase 2, and 800.0 in phase 3. *k*_2_=*k*_3_=0.0, *k*_4_=1.01×10^1^, *k*_5_=31.0, *k*_6_=1.0, *k*_7_=1×10^8^, *k*_8_ not used, *k*_9_=*k*_11_=*k*_13_=*k*_15_=1×10^8^, *k*_10_=*k*_12_=*k*_14_=*k*_16_=1×10^3^, *k*_17_=0.1. Initial concentrations: A_0_=3.000, E_1,0_=1.01×10^−2^, E_2,0_=3.333, Ez_0_=2.994×10^−9^, (E_1_·Ez)_0_=9.102×10^−12^, (E_1_·Ez·E_2_)_0_=1.0×10^−8^, (EzE_2_)_0_=9.871×10^−7^, *Ez_tot_*=1.0×10^−6^.

#### Operational range and irreversibility of the controller

Dual-E control is enabled as long as the condition by Eq 159 is obeyed, i.e., *k*_6_ values need to be lower than *k*_5_, together with *k*_5_<*k*_7_·*Ez_tot_*. For these conditions the rates *v*, *k*_6_, and *j*_5_=*k*_5_*k*_17_/(*k*_17_ + *A_ss_*) are equal. However, when *k*_6_ → *k*_5_, then *A_ss_* → 0, and *j*_5_ → *k*_5_. In the limit, when *k*_5_=*k*_6_, the feedback is broken and *A* does not exert inhibition on *j*_5_.

In the case when *k*_6_>*k*_5_, *E*_2_ will continuously increase, because *v*=*k*_7_(*E*_1_·*Ez*·*E*_2_) balances with *k*_5_, but not with *k*_6_. Due to the continuous increase of *E*_2_ the compensatory flux *k*_4_·(*E*_2_)(*A*) will also increase and drive *A* to zero.

The loss of homeostasis in A when *k*_6_<*k*_5_ is described in Fig 53 where panels (a) and (b) show the numerically calculated *A* and *v* values (gray dots) as a function of *k*_6_ after a simulation time of 48000 time units. In these calculations *k*_5_=31.0 and *k*_17_=0.1. The blue line in panel (a) shows the calculated *A_set_* values by Eq 160. When *k*_6_≥31.0 *A_set_* becomes (formally) negative. In this case *A* is found to decrease as a function of time due to the continuous increase of *E*_2_ as a result of the negative feedback loss. The changes of *A* and *E*_2_ concentrations are shown in panels (c) and (d) when *k*_6_=40.0 (indicated by the red dot and red arrow). Panel (e) shows the concentrations of the different enzymatic species. Indicated in panel (b) is the loss of the negative feedback loop when *k*_6_≥31.0 leading to *A* → 0 and a constant *v_num_*=*k*_5_=31.0.

**Fig 53.**
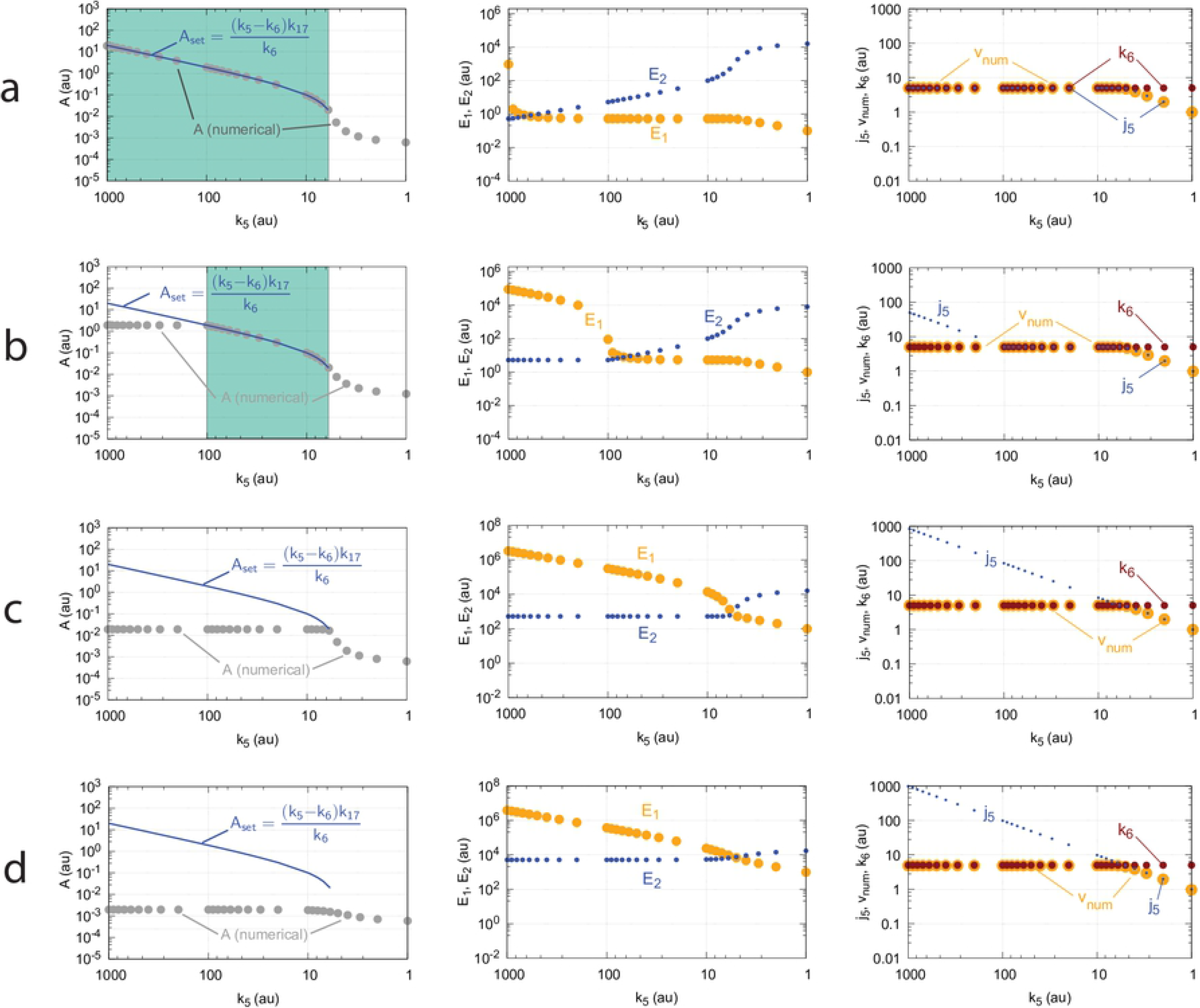
Loss of *A*-homeostasis in the m7 controller with a random-order ternary-complex mechanism (Fig 51) when *k*_6_>*k*_5_. (a) *A_ss_* as a function of *k*_6_. (b) *v_num_* (gray dots) and *v_steady state_* (King-Altman) (red line and small red dots) as a function of *k*_6_. (c) Decrease of *A* as a function of time when *k*_6_=40.0. (d) Steady state in *E*_1_ and wind-up in *E*_2_ when *k*_6_=40.0. (e) Time profiles of the different enzyme species. Rate constants: *k*_1_=100.0, *k*_2_=1.0, *k*_3_=0.0, *k*_4_=1×10^1^, *k*_5_=31.0, *k*_7_=1×10^8^, *k*_8_=0.1, *k*_9_=*k*_11_=*k*_13_=*k*_15_=1×10^8^, *k*_10_=*k*_12_=*k*_14_=*k*_16_=1×10^3^, *k*_17_=0.1. Initial concentrations: A_0_=2.5, E_1,0_=5.5, E_2,0_=52.1, Ez_0_=1×10^−6^, (E_1_·Ez)_0_=0.0, (E_1_·Ez·E_2_)_0_=0.0, (EzE_2_)_0_=0.0, *Ez_tot_*=1.0×10^−6^. Steady state values are determined after a simulation time of 48000 time units.

As mentioned before a necessary condition to obtain robust control is the presence of a sufficient irreversible flux within the controller. This is indicated in Figs 54a-c by using different values of the forward enzymatic rate constants *k*_9_, *k*_11_, *k*_13_, and *k*_15_, while the corresponding reverse rate constants *k*_10_, *k*_12_, *k*_14_, and *k*_16_ are kept constant (1×10^3^). In panel d the enzymatic process is entirely irreversible (*k*_10_, *k*_12_, *k*_14_, and *k*_16_ are all zero), but due to the low value of the forward enzymatic rate constants *k*_10_, *k*_12_, *k*_14_, and *k*_16_ (all 1×10^3^), the controller does not show homeostasis at all, despite being completely irreversible.

**Fig 54.**
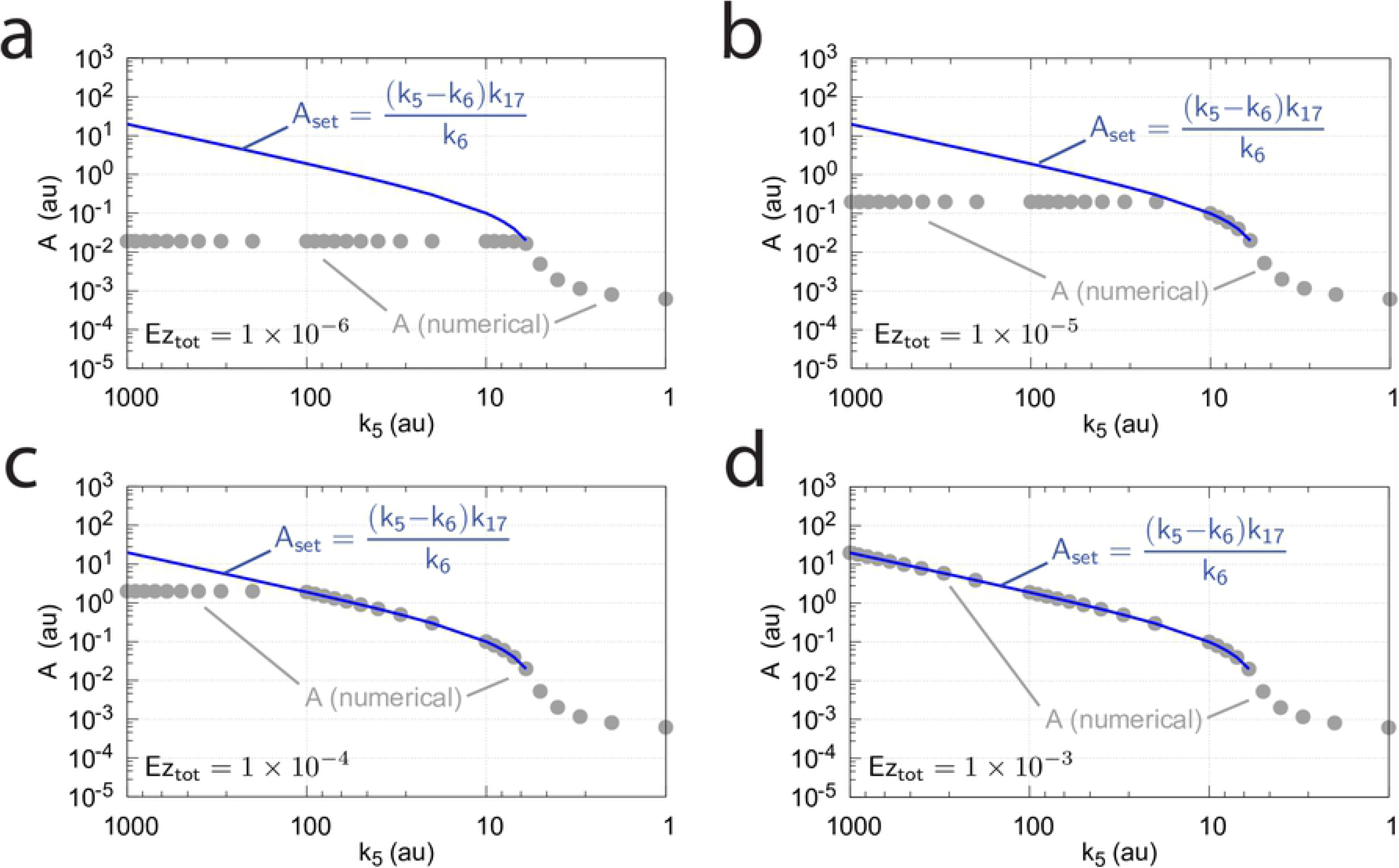
Decrease in the operational range of the enzymatic controller of Fig 51 (in dual-E control mode) as a function to decreased values of the forward enzymatic rate constants *k*_9_, *k*_11_, *k*_13_, and *k*_15_. The *k*_6_ range for which homeostasis is observed is outlined as turquoise areas. (a) *k*_9_=*k*_11_=*k*_13_=*k*_15_=1×10^7^. (b) *k*_9_=*k*_11_=*k*_13_=*k*_15_=1×10^6^. (c) *k*_9_=*k*_11_=*k*_13_=*k*_15_=1×10^4^. The reverse rate constants *k*_10_, *k*_12_, *k*_13_, *k*_15_ are in (a)-(c) kept constant at 1×10^3^. (d) *k*_9_=*k*_11_=*k*_13_=*k*_15_=1×10^3^, while *k*_10_=*k*_12_=*k*_14_=*k*_16_=0. Despite the irreversibility of the system the *k*_9_, *k*_11_, *k*_13_, and *k*_15_ values are too low to enable homeostasis. Other rate constants (a)-(d): *k*_1_=100, *k*_2_=*k*_3_=0, *k*_6_=5.0, *k*_7_=1×10^8^, *k*_17_=0.1. *Ez_tot_*=1×10^−6^. Initial concentrations (a)-(d): A_0_=3.000, E_1,0_=1.01×10^−2^, E_2,0_=3.333, Ez_0_=2.994×10^−9^, (E_1_·Ez)_0_=9.102×10^−12^, (E_1_·Ez·E_2_)_0_=1.0×10^−8^, (EzE_2_)_0_=9.871×10^−7^, *Ez_tot_*=1.0×10^−6^.

The reason behind this failure to show homeostasis at high *k*_5_ values is the incapability of the enzymatic system to absorb the high *j*_5_ inflows. As a result the enzymatic system becomes saturated and *E*_1_ increases continuously.

#### Effect of enzyme concentration

The above incapability of a saturated enzymatic system to maintain homeostatic behavior at large *j*_5_ values can be counteracted by increasing the total enzyme concentration. This is shown in Fig 55, where total enzyme concentration changes from 1×10^−6^ to 1×10^−3^. Clearly, the total amount of the enzyme plays an important role in the performance of catalyzed homeostatic controllers.

**Fig 55.**
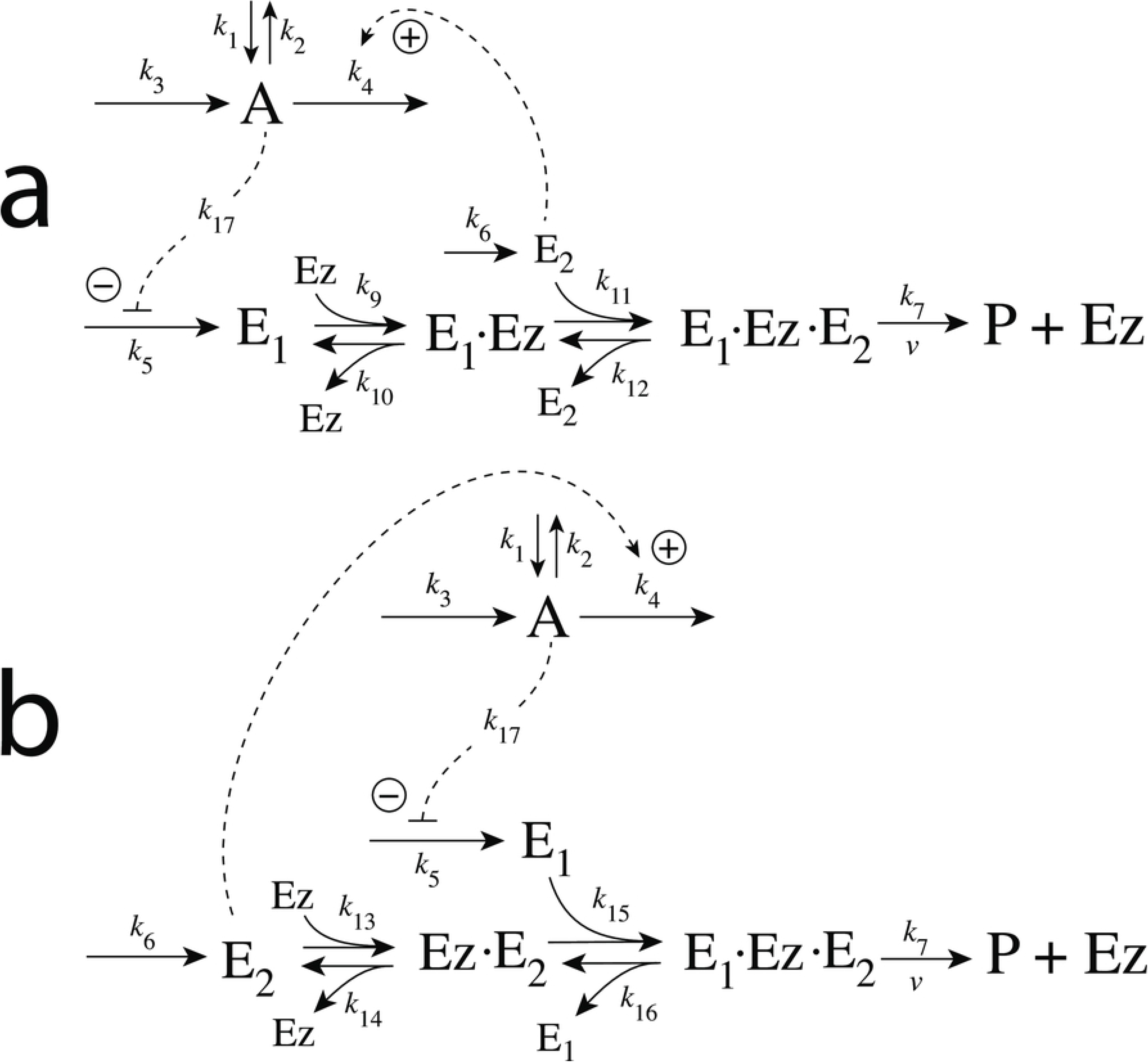
Influence of total enzyme concentration *Ez_tot_* on the performance of the system in Fig 54c. (a) *Ez_tot_*=1.0×10^−6^; (b) *Ez_tot_*=1.0×10^−5^; (c) *Ez_tot_*=1.0×10^−4^; (d) *Ez_tot_*=1.0×10^−3^. Rate constant values as for Fig 54c. Initial concentrations: (a) A_0_=3.000, E_1,0_=1.01×10^−2^, E_2,0_=3.333, Ez_0_=1×10^−6^, (E_1_·Ez)_0_=0.0, (E_1_·Ez·E_2_)_0_=0.0, (EzE_2_)_0_=0.0. (b) A_0_=3.000, E_1,0_=1.01×10^−2^, E_2,0_=3.333, Ez_0_=1×10^−5^, (E_1_·Ez)_0_=0.0, (E_1_·Ez·E_2_)_0_=0.0, (EzE_2_)_0_=0.0. (c) A_0_=3.000, E_1,0_=1.01×10^−2^, E_2,0_=3.333, Ez_0_=1×10^−4^, (E_1_·Ez)_0_=0.0, (E_1_·Ez·E_2_)_0_=0.0, (EzE_2_)_0_=0.0. (d) A_0_=3.000, E_1,0_=1.01×10^−2^, E_2,0_=3.333, Ez_0_=1×10^−3^, (E_1_·Ez)_0_=0.0, (E_1_·Ez·E_2_)_0_=0.0, (EzE_2_)_0_=0.0.

#### Motif 7 controller using compulsory-order ternary-complex mechanisms

As for the other ternary-complex controller motifs there are two compulsory-order mechanisms, one in which *E*_1_ binds first to free *Ez* (Fig 56a), while in the other one (Fig 56b) *E*_2_ binds first to *Ez*.

**Fig 56.**
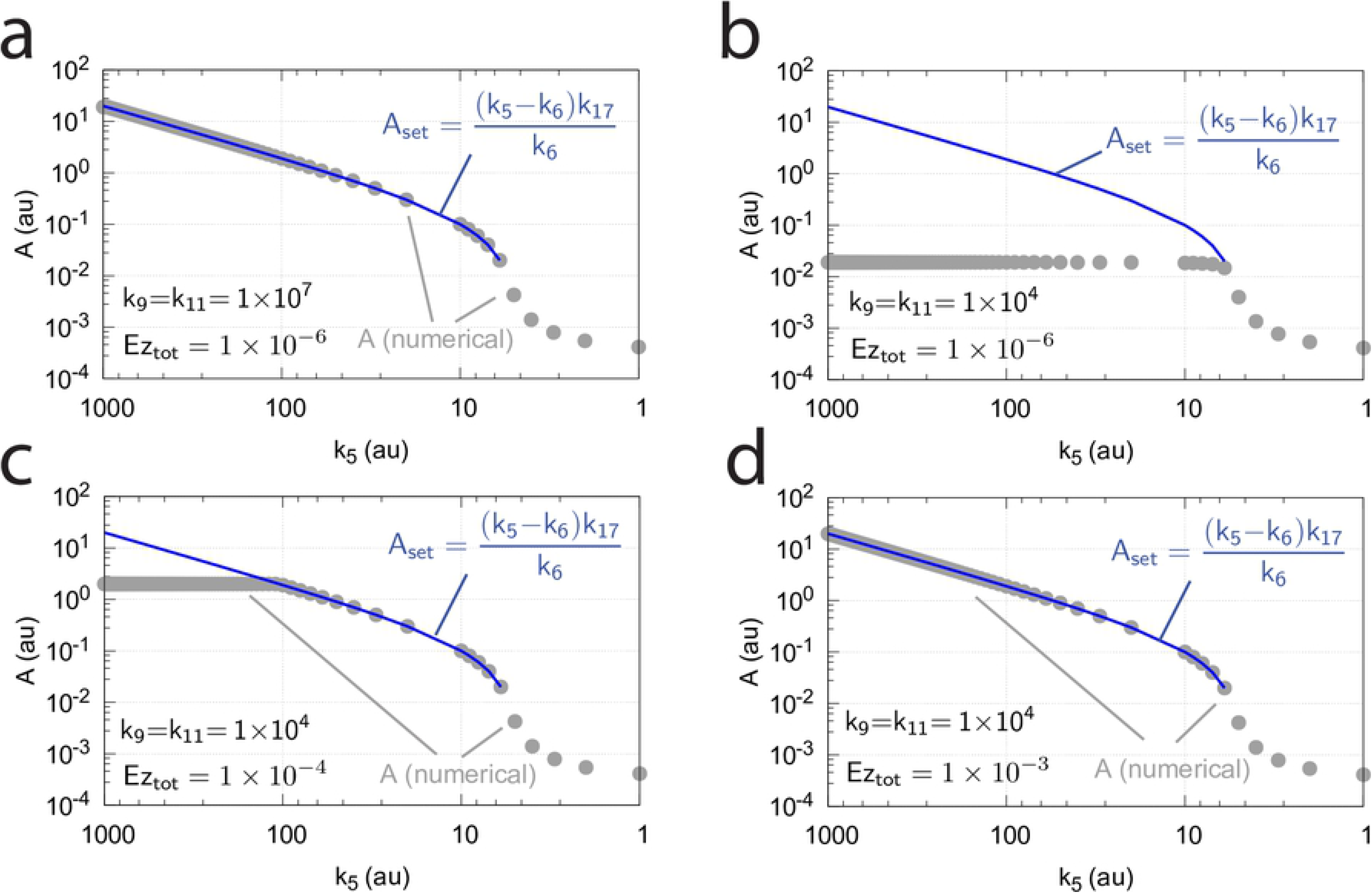
The two compulsory-order ternary-complex mechanisms with feedback motif m7. In (a) *E*_1_ binds first to the free enzyme *Ez*, while in (b) *E*_2_ binds first.

The two compulsory-order mechanisms behave quite similar compared with the random-order mechanism. In the case when *E*_1_ is binding first to the free enzyme *Ez* (Fig 56a), the rate equations are:

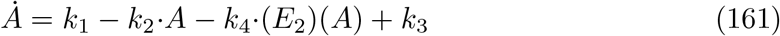

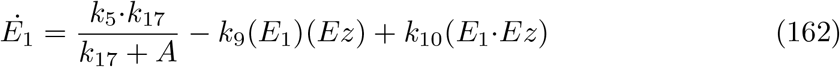

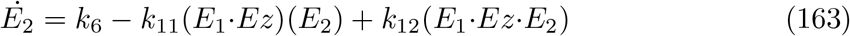

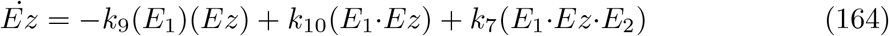

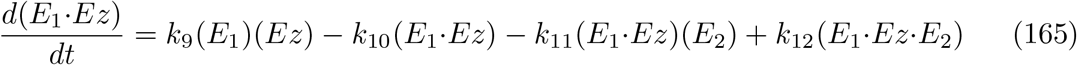

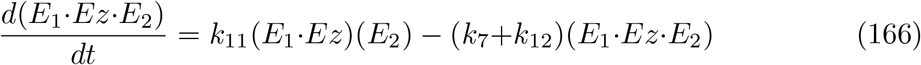

The set-point for the controller in dual-E mode is derived in an analogous way as for the random-order mechanism, i.e. the condition for the operative controller is given by Eq 159 and Eq 160 for the set-point. Also here we have explored how the controller’s performance changes in response to rate constant *k*_5_ and find identical behaviors in response to the enzyme system’s behavior to “absorb” the flux *j*_5_=*k*_5_*k*_17_/(*k*_17_+*A*). High values of *k*_9_ and *k*_11_, as seen in Fig 57a, promote the functionality of the controller, while low *k*_9_ and *k*_11_ values (Fig 57b) lead to a breakdown. As in the random-order case (Fig 55), an increase of the total enzyme concentration leads to an improvement of the controller’s homeostatic behavior. In Figs 57c-d the total enzyme concentrations are increased from 1×10^−6^ to 1×10^−4^ and 1×10^−3^. This allows the controller to maintain the homeostatic *A_set_* for larger *k*_5_ values.

**Fig 57.**
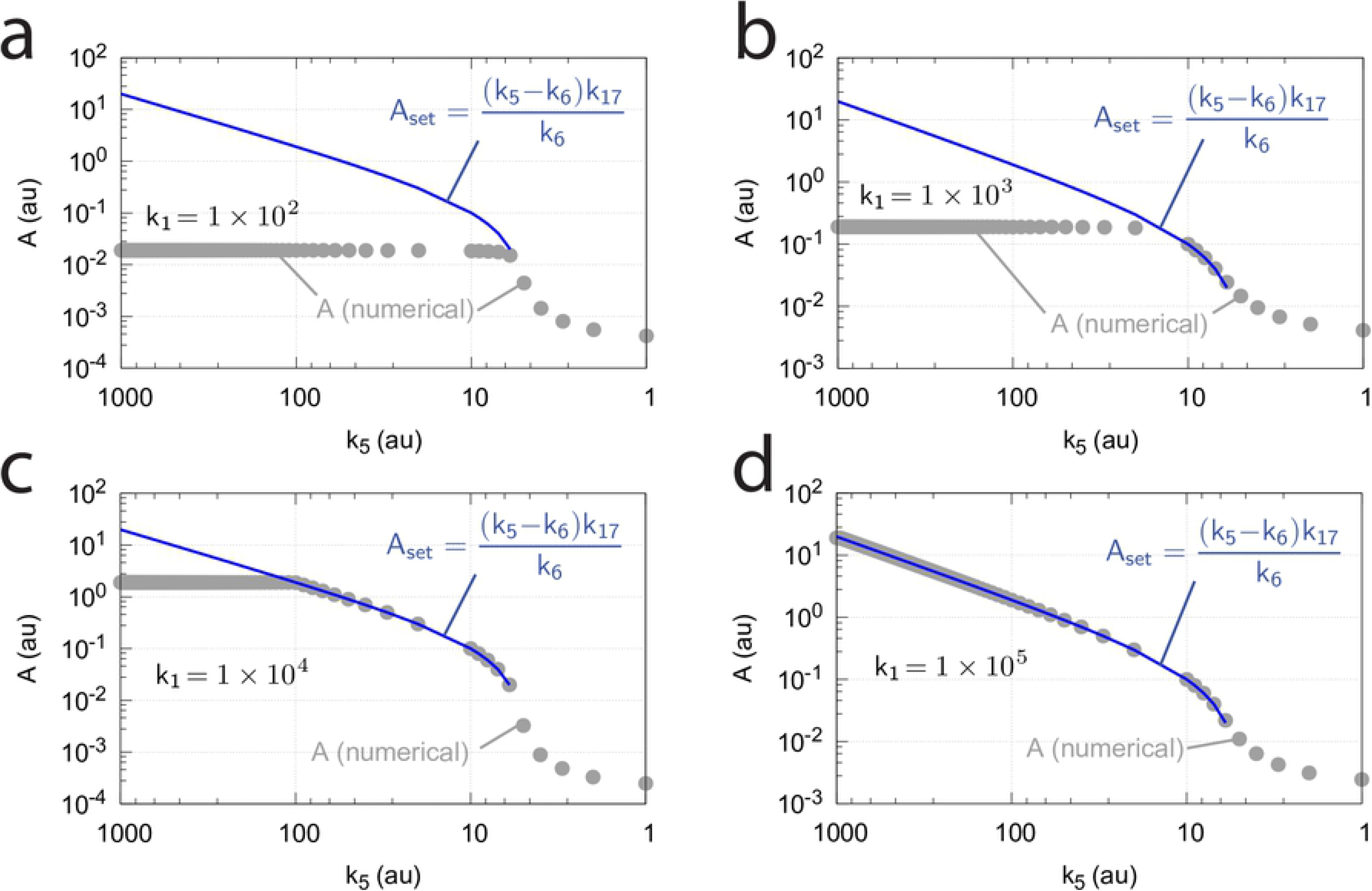
Influence of *k*_9_, *k*_11_, and Ez_tot_ on the operational range of the controller from Fig 56a. (a) Optimum controller behavior for large *k*_9_ and *k*_11_ values (both 1×10^7^) at *Ez_tot_*=1×10^−6^. (b) Reducing *k*_9_ and *k*_11_ to 1×10^4^ leads to a complete loss of the controller’s homeostatic behavior. Although the steady state values of *A* (gray circles) are independent and constant for *k*_5_>*k*_6_, they depend on the perturbation *k*_1_, which will be illustrated below for scheme Fig 56b. (c) Increasing the total enzyme concentration to 1×10^−4^ partially improves the controller’s performance. (d) Increasing the total *Ez* concentration to 1×10^−3^ restores the homeostatic behavior as the increased *k*_9_ and *k*_11_ values in (a) at low *Ez_tot_*. Other rate constants (a)-(d): *k*_1_=100, *k*_2_=*k*_3_=0, *k*_4_=10.0, *k*_6_=5.0, *k*_7_=1×10^8^, *k*_9_=*k*_11_=*k*_13_=*k*_15_=1×10^8^, *k*_10_=*k*_12_=*k*_14_=*k*_16_=1×10^3^, *k*_17_=0.1. Initial concentrations (a)-(d): A_0_=0.08, E_1,0_=5.27×10^−2^, E_2,0_=125.0; (a)-(b) Ez_0_=1.0×10^−6^, (E_1_·Ez)_0_=(E_1_·Ez·E_2_)_0_=0.0; (c) Ez_0_=1.0×10^−4^, (E_1_·Ez)_0_=(E_1_·Ez·E_2_)_0_=0.0; (d) Ez_0_=1.0×10^−3^, (E_1_·Ez)_0_=(E_1_·Ez·E_2_)_0_=0.0. The steady state values of *A* were determined after 6000 time units.

In the case *E*_2_ binds first to the free enzyme *Ez* (Fig 56b), the rate equations become:

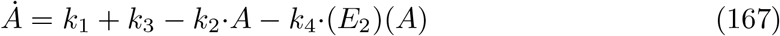

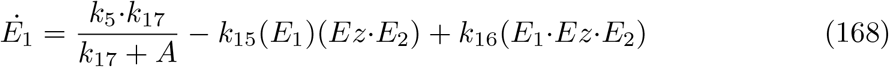

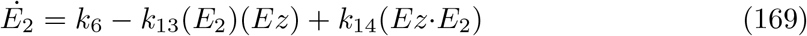

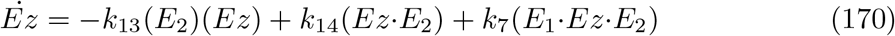

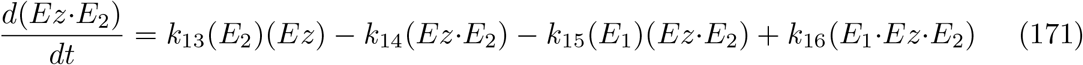

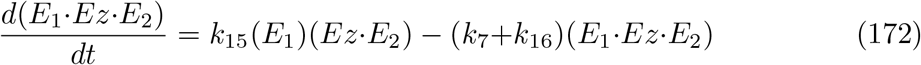

As for the other m7 ternary-complex mechanisms the set-point of *A* is determined by the balance between reaction rates *k*_6_, *j*_5_=*k*_5_*k*_17_/(*k*_17_+*A*), and *v*=*k*_7_(*E*_1_·*Ez*·*E*_2_) (see Eqs 159 and 160). With respect to varying values of *k*_1_ and *k*_5_ the controller’s steady state values in *A* behave precisely as shown in Fig 57 for the other ternary-complex mechanisms, i.e., the loss of homeostasis by low forward rate constants *k*_13_ and *k*_15_ can be counteracted by an increase in the total enzyme concentration.

As we already saw from the previous controllers (see for example the m5 controller, Fig 38) an increase in the perturbation strength (here *k*_1_) will drive the steady-state of the regulated variable *A* towards its theoretical set-point *A_set_*. This is also observed for the m7-type of controllers for dual-E control. As seen in Fig 58, increasing *k*_1_ values extend the homeostatic region of the controller. This same pattern of *A* as a function of *k*_5_ for different *k*_1_ values is also observed for the other ternary-complex mechanisms (Fig 56a and Fig 51).

**Fig 58.**
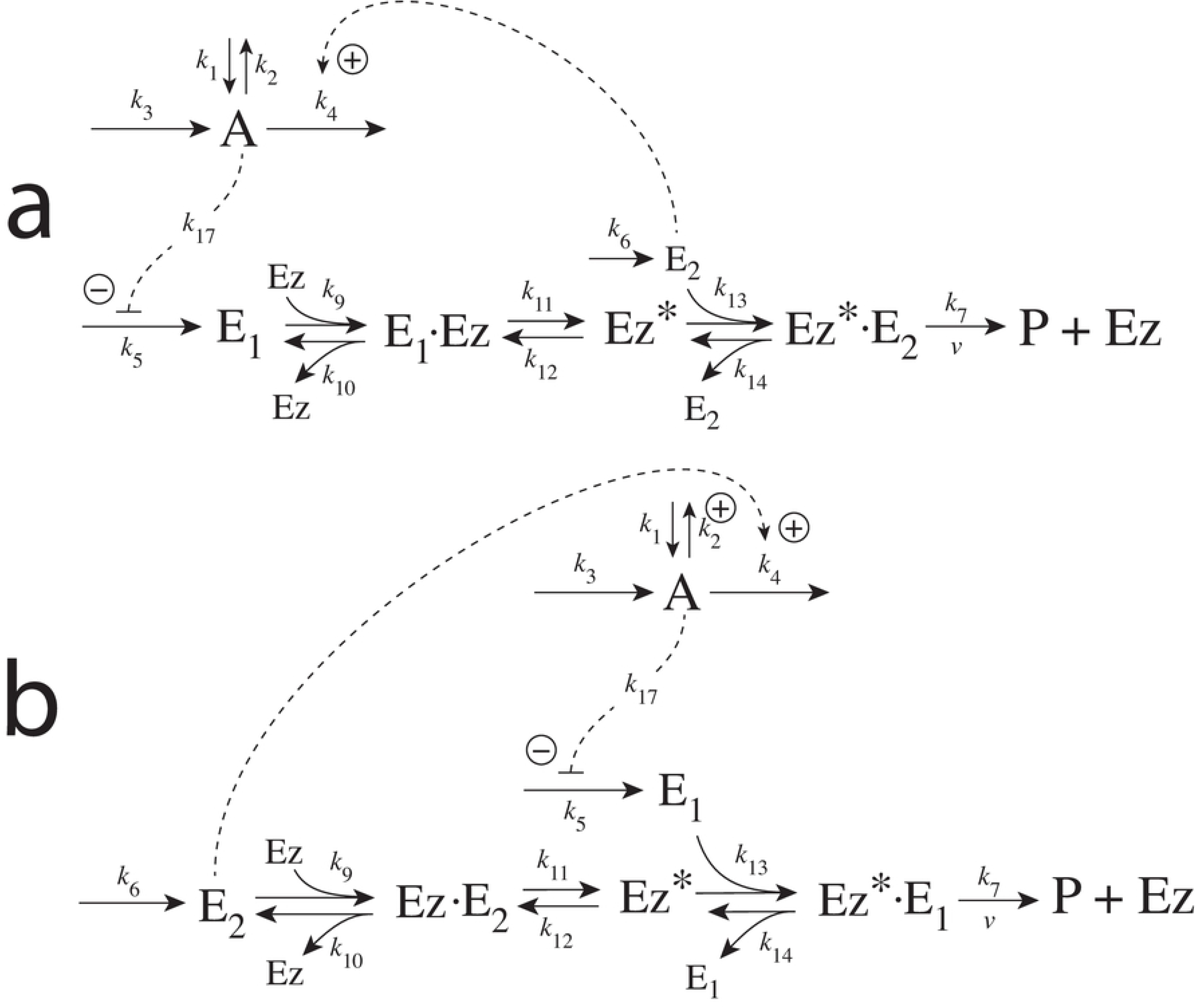
Influence of *k*_1_ on the operational range of the m7 ternary-complex controllers. The results using the scheme of Fig 56b are shown. (a) *k*_1_=1×10^2^. (b) *k*_1_=1×10^3^. (c) *k*_1_=1×10^4^. (d) *k*_1_=1×10^5^. Other rate constants (a)-(d): *k*_2_=*k*_3_=0, *k*_4_=10.0, *k*_6_=5.0, *k*_7_=1×10^8^, *k*_13_=*k*_15_=1×10^4^, *k*_14_=*k*_16_= *k*_17_=1×10^3^. Initial concentrations (a)-(d): A_0_=1.88, E_1,0_=5.39×10^2^, E_2,0_=5.315, Ez_0_=1.0×10^−6^, (E_1_·Ez)_0_=(E_1_·Ez·E_2_)_0_=0.0. Due to a slow response (large response time) of the controller at higher *k*_1_, steady state values of *A* were determined after 1×10^6^ time units.

#### Motif 7 dual-E controller removing *E*_1_ and *E*_2_ by ping-pong mechanisms

Fig 59 shows the reaction scheme when in a m7 controller configuration *E*_1_ and *E*_2_ are removed by the two enzymatic ping-pong mechanisms when *E*_1_ binds first to *Ez* (Fig 59a) or when *E*_2_ binds first to *Ez* (Fig 59b).

**Fig 59.**
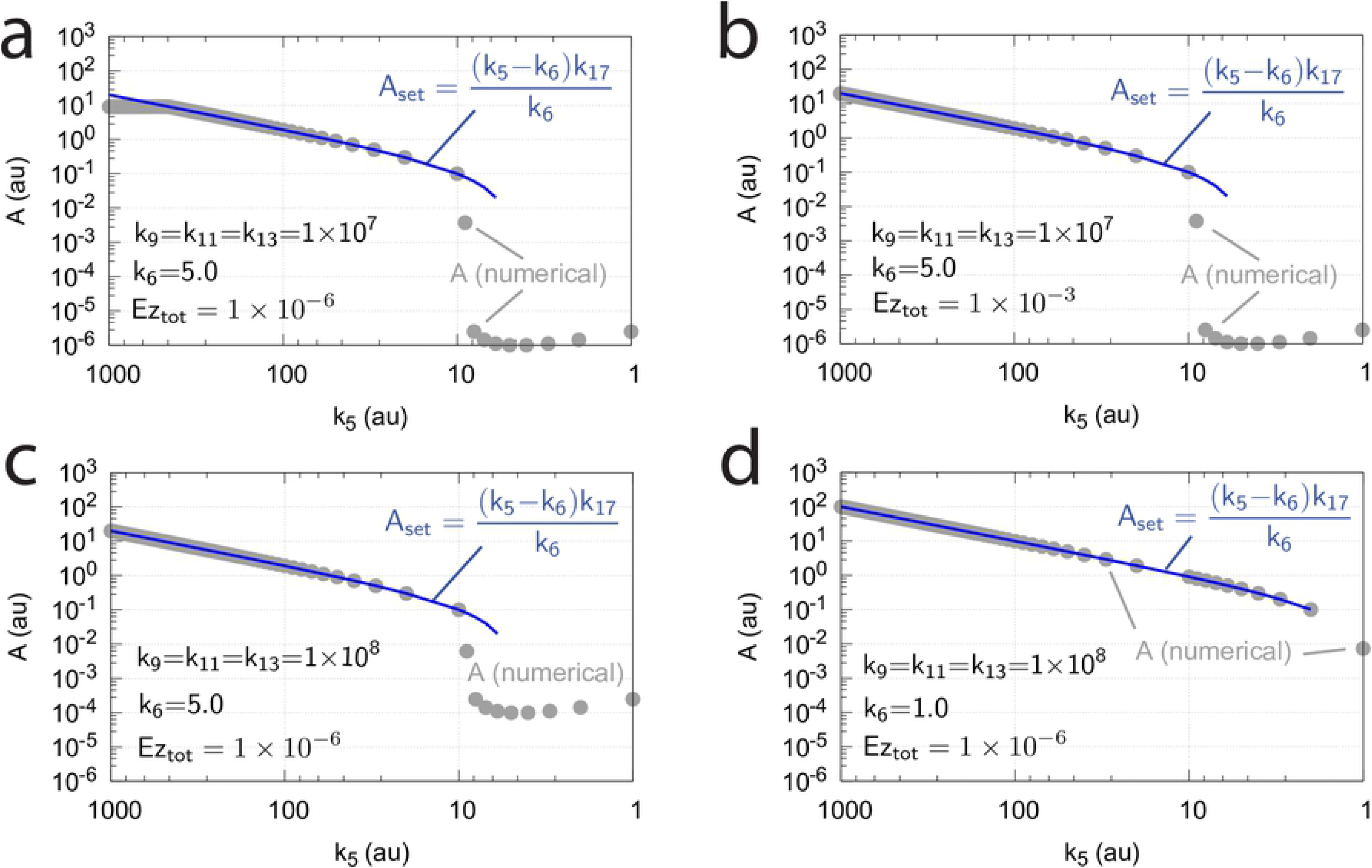
Reaction schemes of the two m7-type of controllers when *E*_1_ and *E*_2_ (Fig 3) are removed by enzyme *Ez* using ping-pong mechanisms with (a) *E*_1_ binding first, or (b) when *E*_2_ binds first.

In the case *E*_1_ binds first to *Ez* (Fig 59a) the rate equations are:

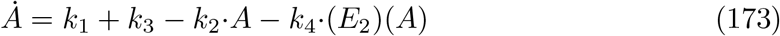

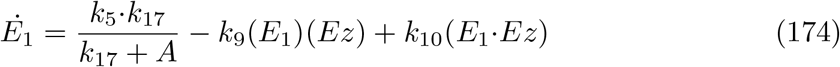

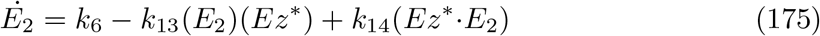

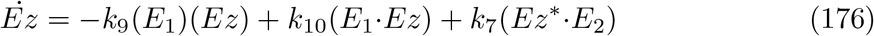

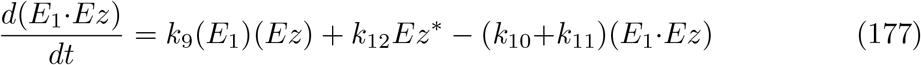

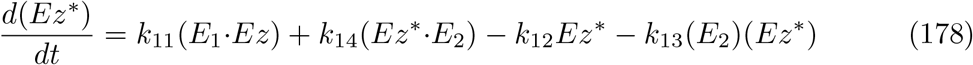

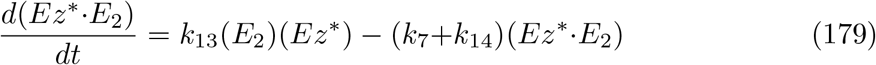

#### Minor differences between the m7 ping-pong and ternary-complex mechanisms

The dynamic behaviors of the ping pong-mechanisms are very similar to the (m7) ternary-complex mechanisms. Also here *A_set_* is determined by the balancing of the three fluxes *j*_5_=*k*_5_*k*_17_/(*k*_17_+*A*), the inflow described by *k*_6_, and the rate *v*=*k*_7_(*Ez*^*^·*E*_2_) making *P*. Accordingly, *A_set_* is described by Eq 160. Also, an increase of total enzyme concentration and an increase of the forward enzymatic rate constants *k*_9_, *k*_11_,… will improve the homeostatic performance of the ping-pong controllers.

However, since the ping-pong mechanisms have a slightly longer enzymatic reaction chain in comparison with the ternary-complex mechanisms, in the ping-pong case larger forward enzymatic rate constants values are needed together with lower *k*_6_ values to match the fluxes *j*_5_, *k*_6_, and *v* to achieve moving *A_ss_* to its set-point. The influences of the forward enzymatic rate constants and the total *Ez* concentration are illustrated in Fig 60 where numerically calculated *A* values are compared with the theoretical set-point *A_set_*. In comparison with the ternary-complex mechanism results from Fig 57a Fig 60a show the behavior of the ping-pong mechanism when *Ez_tot_*=1×10^−6^, and *k*_9_=*k*_11_=*k*_13_=1×10^7^. Unlike in the ternary-complex mechanism, in the ping-pong case deviations between the numerically calculated A values and *A_set_* are observed at the higher (*k*_5_>460) and lower *k*_5_<10 ends of the *k*_5_ scale. When in Fig 60a *k*_5_ gets higher than 460 the enzymatic system cannot absorb the inflow flux *j*_5_=*k*_5_*k*_17_/(*k*_17_+*A*). As a result, *E*_1_ shows a linear increase in time, with a slope which is dependent on the value of *k*_5_, but where *A* becomes constant and independent of *k*_5_. At the lower end of the *k*_5_ scale (*k*_5_<10) the value of *k*_6_ is too high to get absorbed by *Ez*^*^·*E*_2_. This has the result that *E*_2_ shows a liner increase in time and an increasing compensatory flux *k*_4_·*A*·*E*_2_ with *A* decreasing continuously without reaching a steady state. The loss of homeostasis at high *k*_5_ values can be overcome by either increasing the total amount of enzyme (Fig 60b) or by increasing the values of the forward rate constants *k*_9_, *k*_11_, and/or *k*_13_ (Fig 60c).

**Fig 60.**
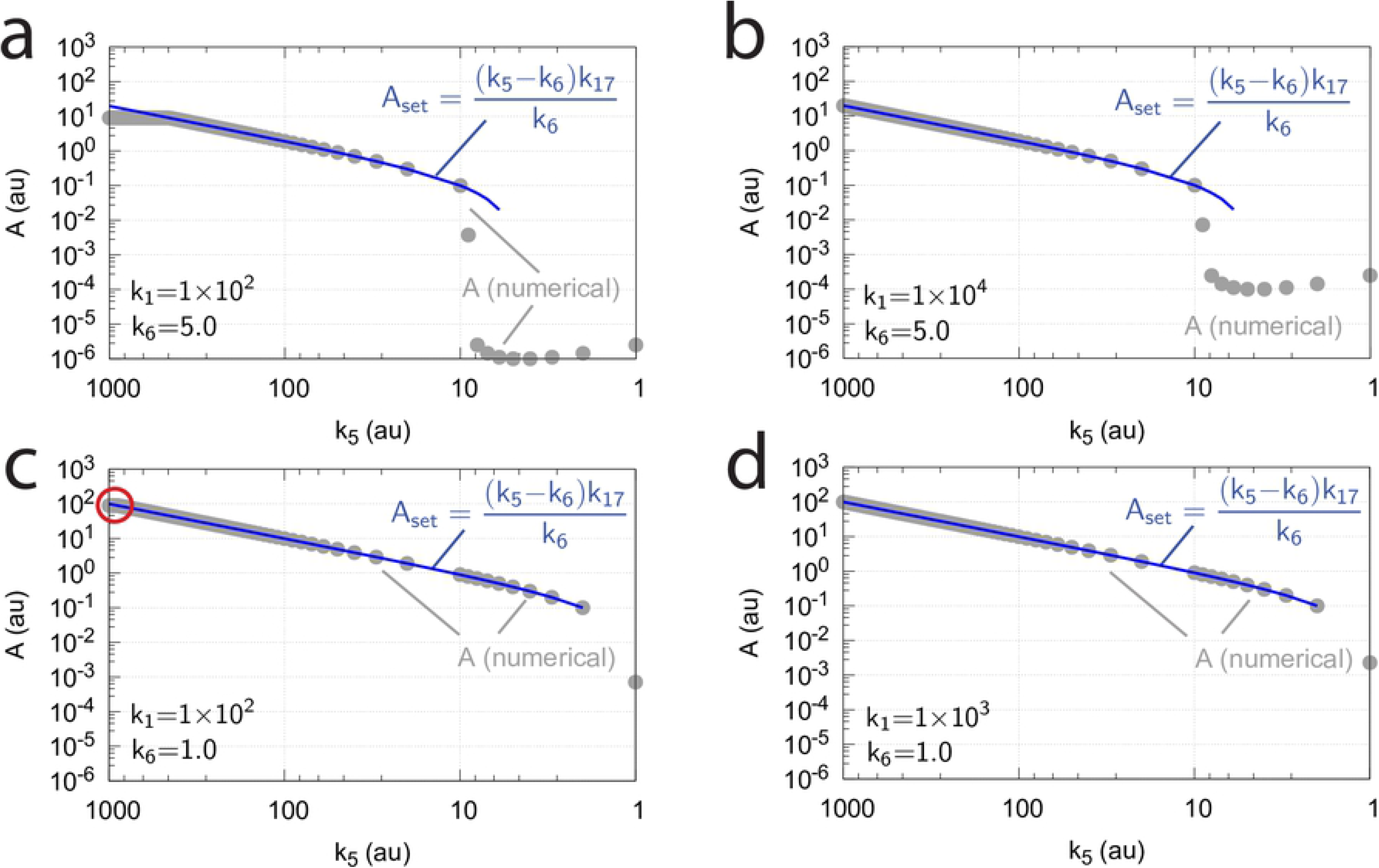
Influence of *k*_6_, *k*_9_, *k*_11_, *k*_13_ and *Ez_tot_* on the homeostatic behavior of the m7 ping-pong controller when *E*_1_ binds first to free enzyme *Ez*. Numerical *A* values calculated after 10^4^ time units are compared with corresponding analytical expressions of *A_set_* as a function of *k*_5_. (a) *k*_6_=5.0, *k*_9_=*k*_11_=*k*_13_=1×10^7^, and *Ez_tot_*=1×10^−6^. (b) *k*_6_=5.0, *k*_9_=*k*_11_=*k*_13_=1×10^7^, and *Ez_tot_*=1×10^−3^. (c) *k*_6_=5.0, *k*_9_=*k*_11_=*k*_13_=1×10^8^, and *Ez_tot_*=1×10^−6^. (d) *k*_6_=1.0, *k*_9_=*k*_11_=*k*_13_=1×10^8^, and *Ez_tot_*=1×10^−6^. Other rate constants (a)-(d): *k*_1_=100.0, *k*_2_=*k*_3_=0, *k*_4_=10.0, *k*_7_=1×10^8^, *k*_12_=*k*_14_=1×10^3^. Initial concentrations (a), (c), and (d): A_0_=3.0, E_1,0_=1.0×10^−2^, E_2,0_=3.0×10^2^, Ez_0_=1.0×10^−6^, 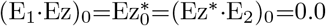. Initial concentrations (b): A_0_=3.0, E_1,0_=1.0×10^−2^, E_2,0_=3.0×10^2^, Ez_0_=1.0×10^−3^, 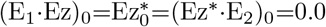.

An increase of the perturbation *k*_1_ leads also in the ping-pong controllers to an improvement in the homeostatic accuracy as for example observed in the m5 controllers, but not at low *k*_5_ values. This is indicated in Fig 61. In panel a we have rate constant values for *k*_1_, *k*_6_, *Ez_tot_*, and the forward enzymatic rate constants (*k*_9_, *k*_11_, and *k*_13_) as in Fig 60a and as in the ternary-complex mechanisms of Fig 57a. An increase of *k*_1_ from 1×10^2^ to 1×10^4^ in the ping-pong mechanism (Fig 61b) does improve the homeostatic response of the controller at high *k*_5_ values, but not at low *k*_5_, where the high inflow rate by *k*_6_ cannot be absorbed. In fact, a decrease of *k*_6_ from 5.0 to 1.0 leads to homeostasis for all *k*_5_ values with *A_set_>*0 (Fig 61c).

**Fig 61.**
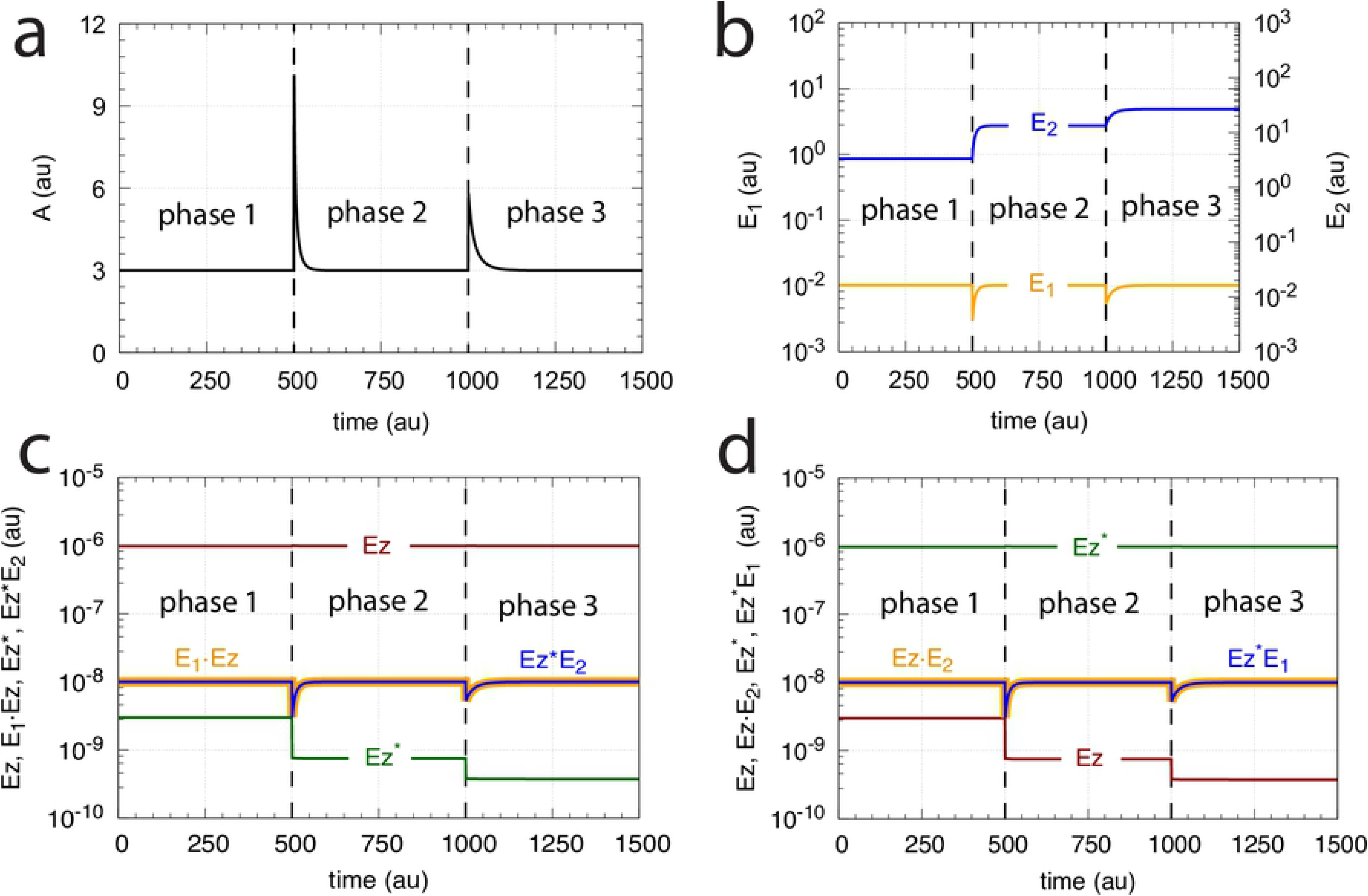
Influence of *k*_1_ on the homeostatic behavior of the m7 ping-pong controller when *E*_1_ binds first to free enzyme *Ez*. Since the response of the controller at higher *k*_1_ values becomes significantly slower the numerical *A* values are calculated after 10^6^ time units and compared with positive *A_set_* (blue lines) as a function of *k*_5_. (a) *k*_1_=100.0, *k*_6_=5.0. The controller looses homeostatic control in the *k*_5_ range from 460-1000. (b) Increasing *k*_1_ from 100.0 to 10000.0 moves *A_ss_* to *A_set_* for the higher *k*_5_ values, but not for the lower *k*_5_ values. (c) A decrease of *k*_6_ from 5.0 to 1.0 while *k*_1_ is kept at 100.0 gives a general improvement of the homeostatic performance of the ping-pong controller, except for the higher end *k*_5_ range between 900-1000, where *A_ss_* becomes constant (indicated by the red circle). (d) Low *k*_6_ (1.0) and higher *k*_1_ (1000.0) shows *A_ss_* values that match *A_set_*. Other rate constants (a)-(d): *k*_2_=*k*_3_=0, *k*_4_=10.0, *k*_7_=1×10^8^, *k*_9_=*k*_11_=*k*_13_=1×10^7^, *k*_12_=*k*_14_=1×10^3^, *k*_17_=0.1. Initial concentrations (a)-(d): A_0_=3.0, E_1,0_=1.0×10^−2^, E_2,0_=3.0×10^2^, Ez_0_=1.0×10^−6^, 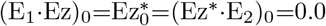.

An analysis of the two m7 ping-pong mechanisms in Fig 59 shows that their responses in *A*, *E*_1_, and *E*_2_ are, for different *k*_1_ perturbation strengths, identical (see Figs 62a and b). Both use *E*_2_ as the variable which controls the compensatory flux *j_comp_*=*k*_4_·*A·E*_2_. The roles of the enzymatic species *Ez*, *Ez*^*^, *E*_1_·*Ez* and *Ez*^*^*E*_2_ in the mechanism of Fig 59a are in the mechanism of Fig 59b replaced by the respective species *Ez*^*^, *Ez*, *Ez·E*_2_, and *Ez*^*^*E*_1_; see Figs 62c and d. While the concentrations in *A*, *E*_1_, *E*_2_, *E*_1_·*Ez*, *Ez*^*^*E*_2_, *Ez·E*_2_, and *Ez*^*^*E*_1_ are identical as a function of *k*_5_, the concentrations of *Ez* and *Ez*^*^ are different, but interchange in dependence whether *E*_1_ or *E*_2_ binds first to free *Ez*. The numerical results are shown in Fig 63 for the individual reaction species.

**Fig 62.**
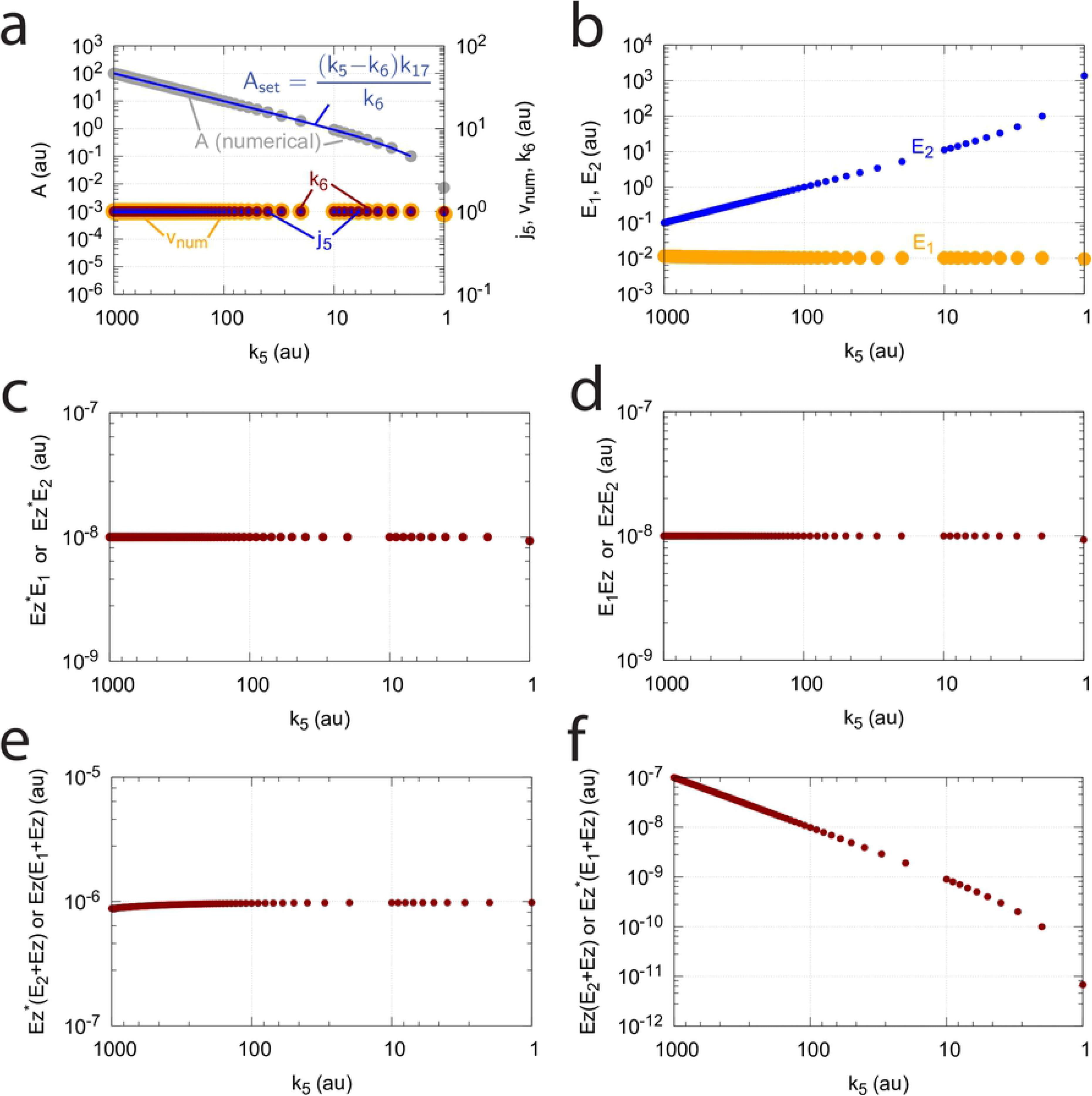
The m7 ping-pong mechanisms (Fig 59) show identical homeostatic responses for step-wise changes in *k*_1_. Phase 1: *k*_1_=100.0, phase 2: *k*_1_=400.0, phase 3: *k*_1_=800.0. (a) *A* as a function of the step-wise changes in *k*_1_ for both controllers. (b) Concentration profiles of *E*_1_ and *E*_2_ for both controllers. (c) Concentration profiles of the enzyme species for the mechanism in Fig 59a. (d) Concentration profiles of the enzyme species for the mechanism in Fig 59b. Other rate constants: *k*_2_=*k*_3_=0, *k*_4_=10.0, *k*_5_=31.0, *k*_6_=1.0, *k*_7_=1×10^8^, *k*_9_=*k*_11_=*k*_13_=1×10^8^, *k*_12_=*k*_14_=1×10^3^, *k*_17_=0.1. Initial concentrations for the controller of Fig 59a: A_0_=3.0, E_1,0_=1.0×10^−2^, E_2,0_=3.33, Ez_0_=9.77×10^−7^, (E_1_·Ez)_0_=1.0×10^−8^, 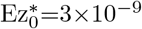, (Ez^*^·E_2_)_0_=1.0×10^−8^. Initial concentrations for the controller of Fig 59b: A_0_=3.0, E_1,0_=1.0×10^−2^, E_2,0_=3.33, Ez_0_=3×10^−9^, (Ez·E_2_)_0_=1.0×10^−8^, 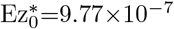, (Ez^*^·E_1_)_0_=1.0×10^−8^.

**Fig 63.**
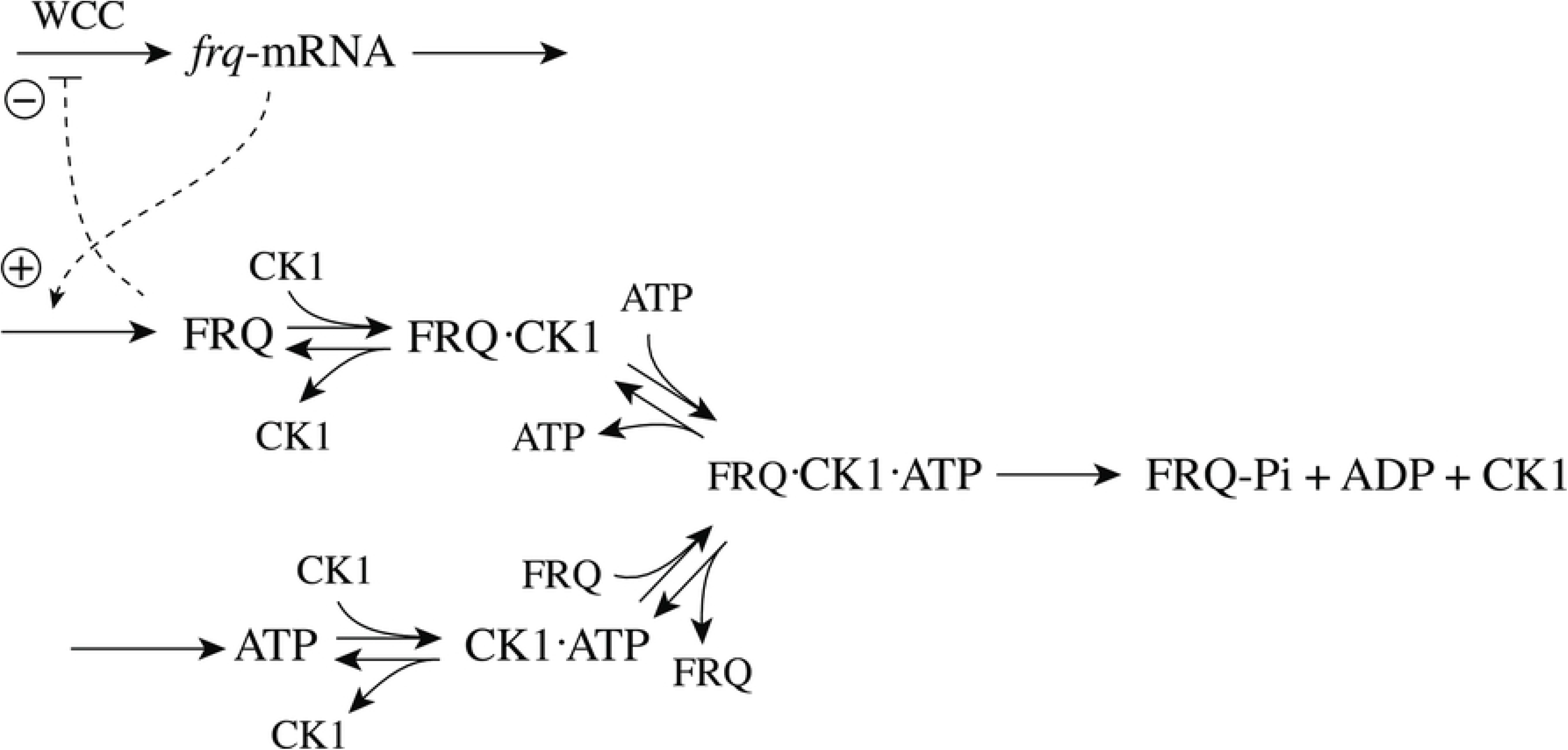
Concentration profiles of reaction species of the two m7 ping-pong mechanisms (Fig 59) as a function of *k*_5_. (a) Left ordinate: Numerical steady state values of *A* (gray dots) in comparison with the theoretical set-point *A_set_* (Eq 160, blue line). Ordinate to the right: *k*_6_ (red dots), steady state values of *j*_5_=*k*_5_*k*_17_/(*k*_17_+*A*) (blue dots), and numerically calculated reaction rate *v_num_*=*dP/dt* (orange dots). (b) Steady state values of *E*_1_ (orange dots) and *E*_2_ (blue dots). (c) Steady state profiles of *Ez*^*^*E*_2_ (Fig 59a) or *Ez*^*^*E*_1_ (Fig 59b). (d) Steady state profiles of *E*_1_*Ez* (Fig 59a) or *EzE*_2_ (Fig 59b). (e) Steady state profile of *Ez* when *E*_1_ binds first to it (Fig 59a) or profile for *Ez*^*^ when *E*_2_ binds first to *Ez* (Fig 59b). (f) Steady state profile of *Ez* when *E*_2_ binds first to *Ez* (Fig 59b) or profile for *Ez*^*^ when *E*_1_ binds first to *Ez* (Fig 59a). Rate constants: *k*_1_=100.0, other rate constants as in Fig 62. Initial concentrations (a)-(f): A_0_=3.0, E_1,0_=1.0×10^−2^, E_2,0_=3.0×10^2^, Ez_0_=1.0×10^−6^, 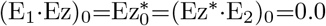.

## Discussion

There are presently three kinetic approaches how error integration (Fig 1) can be achieved leading to perfect adaptation or homeostasis. One approach is the use of applying zero-order kinetics in the removal of the controller variable *E* [1, 2, 4–6, 15, 26]. A second approach [8, 27, 28] is based on a first-order autocatalytic production of *E* combined with its first-order removal. Finally, a third approach is based on antithetic control (described here also as dual-E control) [7, 8, 10, 11], where one of the controller variables (for example *E*_1_) participates in a negative feedback and reacts with a second controller variable *E*_2_, for example as described by Eq 3.

The advantage of the antithetic approach is that the removal of *E*_1_ and *E*_2_ does not necessarily need to be precisely a second-order process as formulated by Eq 3, but can in principle be of any type of kinetics. As practically all biochemical processes are catalyzed by enzymes, we have focussed here on mechanisms which remove *E*_1_ and *E*_2_ by classical two-substrate enzyme kinetics [12, 13]. In addition, taking a previously suggested basic set of negative feedback loops (controller motifs m1-m8) as a starting point, we have extended in Fig 3 this set for dual-E/antithetic control. Using enzyme kinetics with *E*_1_ and *E*_2_ as substrates allows for a large variety of processes as candidates for robust homeostasis. In the following we discuss a few examples where robust regulation appears associated to enzymatic dual-E/antithetic control.

### Protein phosphorylation

Regulation by phosphorylating enzymes is observed in practically all aspects of life [29]. The enzymes, protein kinases, use as substrate a target protein and MgATP. A general feature of protein kinases is that they follow compulsory-order or random-order ternary-complex mechanisms [30]. In the following we give two examples that describe m2 control where ATP and the target protein are processed by a kinase using a ternary-complex mechanism.

### Circadian rhythms

Circadian rhythms play an important part in the adaptation of organisms to their environment, in particular to the day/night and seasonal changes on earth. The molecular bases of circadian rhythms are transcriptional-translational negative feedback loops which oscillate with a period of circa 24 hours [31]. Using the model organism *Neurospora crassa* phosphorylation was found to serve two functions: firstly, to close the negative feedback loop by phosphorylating the transcription factor WCC (White Collar Complex). The WCC phosphorylation leads to its inhibition by its gene product, the protein FREQUENCY (FRQ) [32, 33], Secondly, FRQ, which is central to the *Neurospora* circadian pacemaker [34] is phosphorylated by CK1 with the result that phosphorylated FRQ is no longer able to inhibit WCC. Fig 64 indicates the central negative feedback loop in the *Neurospora* circadian clock describing the phosphorylation of FRQ by CK1 as a random-order ternary-complex mechanism, thereby moving FRQ out of the negative feedback loop. FRQ is phosphorylated at multiple sites [35] and hyper-phosphorylated FRQ is finally degraded. It should be noted that analogous feedback loops with post-translational phosphorylation have also been observed for the *Drosophila* circadian clock [36, 37].

**Fig 64.**
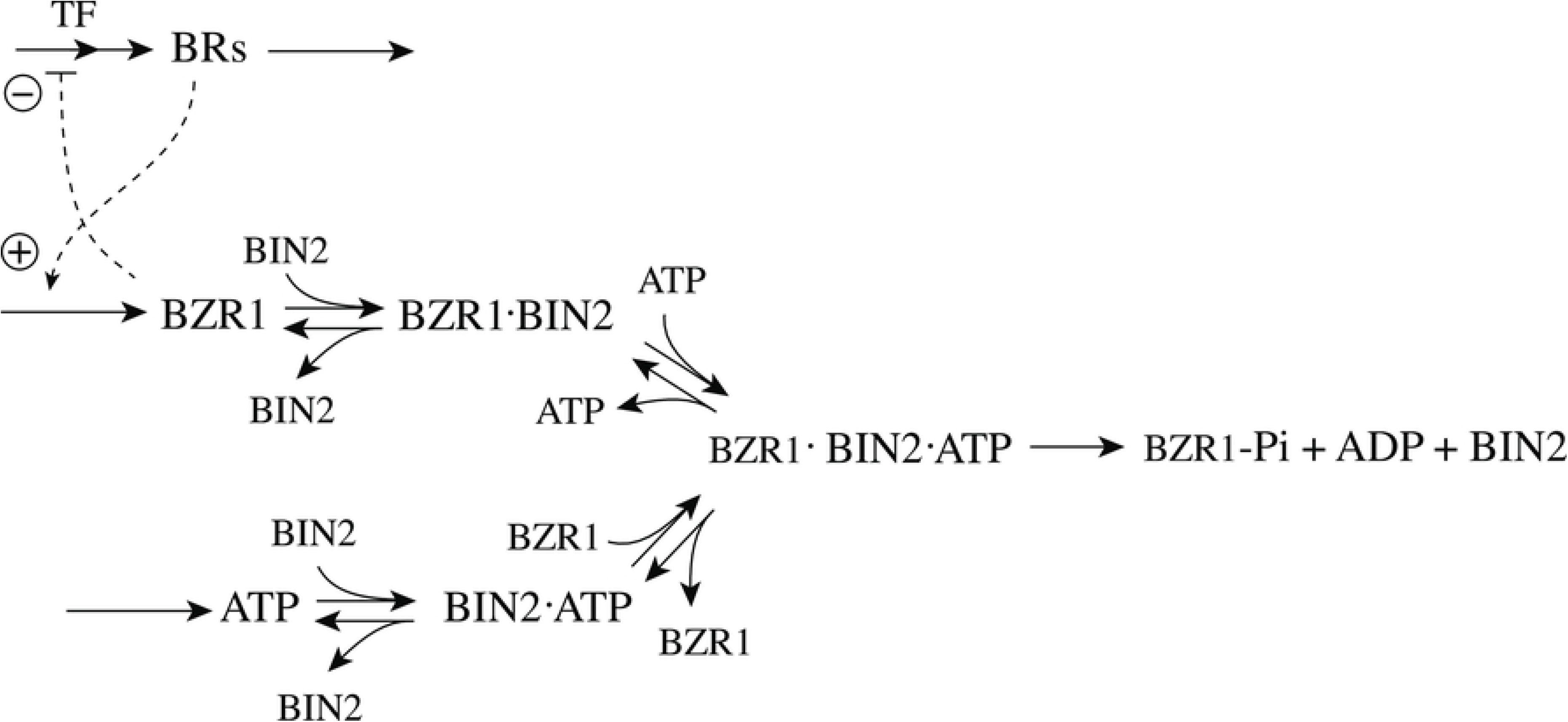
Central transcriptional-translational negative feedback loop of the *Neurospora* circadian clock. In the presence of FRQ the transcription factor White Collar Complex (WCC) is phosphorylated, which leads to its inhibition by FRQ and thereby suppressing FRQ synthesis. FRQ on its side is phosphorylated, which moves the inhibitory FRQ form out of the loop and leads to its eventually to its degradation. The dual-E controller suggests that *frq*-mRNA is under homeostatic control with respect to variable *frq*-mRNA degradation.

In comparison with our m2 calculations above, Fig 64 predicts that *frq*-mRNA appears to be under homeostatic regulation with respect to its degradation. This prediction is indeed in agreement with experimental findings by Liu et al. [38]. Their results indicate that the level of *frq*-mRNA, although changing on a circadian time scale, is on average not altered at different temperatures. Since, furthermore, the circadian period is compensated towards variations in temperature (temperature-compensation) [39–41], it will be interesting to investigate how FRQ phosphorylation by CK1, leading to putative *frq*-mRNA homeostasis, also contributes to temperature compensation in the *Neurospora* circadian clock as indicated by recent experiments [42].

### Brassinosteroid homeostasis

Brassinosteroids (BRs) are plant hormones which have influence on plant growth and development, and adapt plants to environmental stresses. Plants lacking BRs show dwarf growth and abnormal organs [43]. BRs bound to their receptor BZR1 produce BZ1, which inhibits the transcription of the BR genes by binding to promoter regions of different genes in the BR synthesis pathway [44, 45]. The GSK3-like kinase BIN2 phosphorylates BZR1, which then leads to its proteasomal degradation [46]. Fig 65 shows the removal of BZR1 out of the negative feedback loop by BIN2 phosphorylation using a random-order ternary-complex mechanism [30].

**Fig 65.**
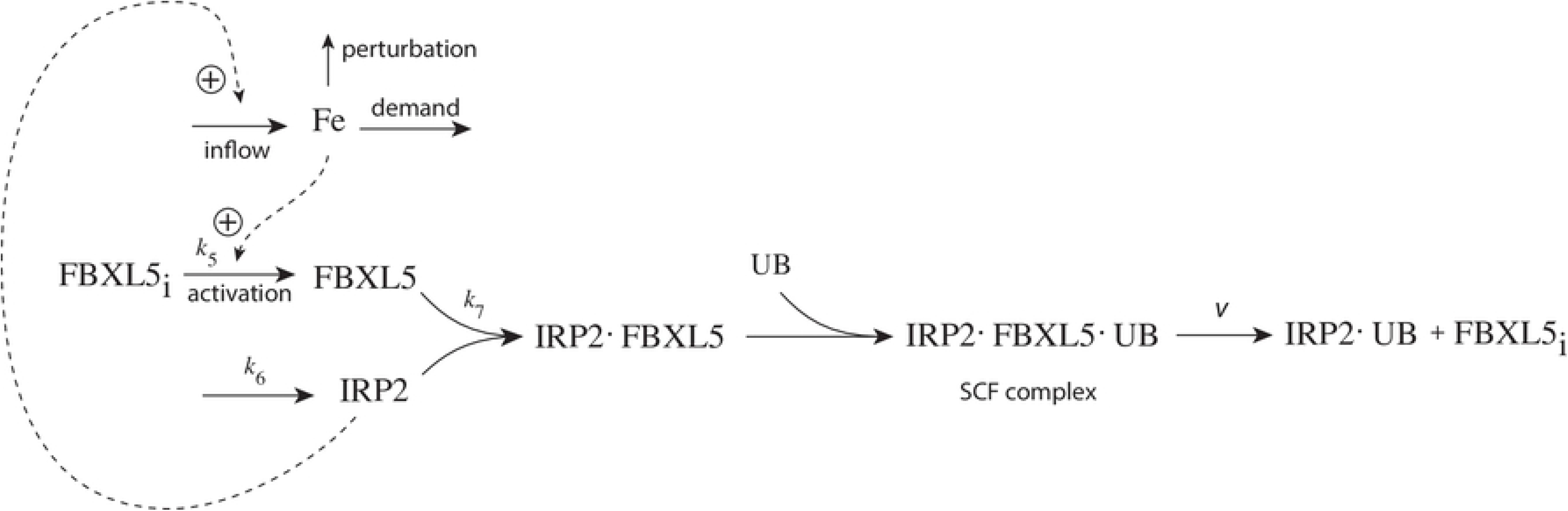
M2 dual-E control loop of Brassinosteroid homeostasis. Brassinosteroid genes are transcribed where TF indicate a set of transcription factors. When BRs bind to their receptors unphosphorylated BZ1 is produced which binds to the transcription factor and thereby inhibits Brassinosteroid transcription. The GSK3-like kinase BIN2 phosphorylates BZR1 and removes it from the negative feedback loop. Phosphorylated BZR1 is finally degraded by the proteasome.

### Ubiquitination and proteasomal degradation

In metal-ion homeostasis many of the controller molecules are subject to proteasomal degradation in a metal-ion dependent fashion (for a summary, see the Supporting Material of Ref [6]). In proteasomal degradation, ubiquitin, a small protein, is moved through a cascade of three ligases (E1-E3; not to be confused with the controllers *E*_1_ and *E*_2_ above) and then added on to the target protein [47]. Repeated ubiquitin ligation of the target protein leads then finally to its degradation by the proteasome.

A relatively well understood example is mammalian iron homeostasis. At low iron levels IRP2 together with IRP1 promote the inflow of iron by stabilizing mRNAs which code for proteins that are necessary for iron supply. Results by Vashisht et al. [48] indicate that IRP2 is degraded in an iron-dependent manner where the F-box protein FBXL5 catalyzes IRP2 ligation with ubiquitin. While in this case three substrates are involved (iron, IRP2 and ubiquitin), dual-E control as described above cannot directly applied. However, the indication by Vashisht et al. [48] that iron stabilizes/activates FBXL5 leads to the following m1 dual-E mechanism (Fig 66) where iron activates FBXL5 from a pool of inactive enzyme. This allows the binding of IRP2 forming a SCF complex [48, 49]. For simplicity, the other components of the SCF complex are not shown and the pool of inactive enzyme (FBXL5_i_) is considered to be constant.

**Fig 66.**
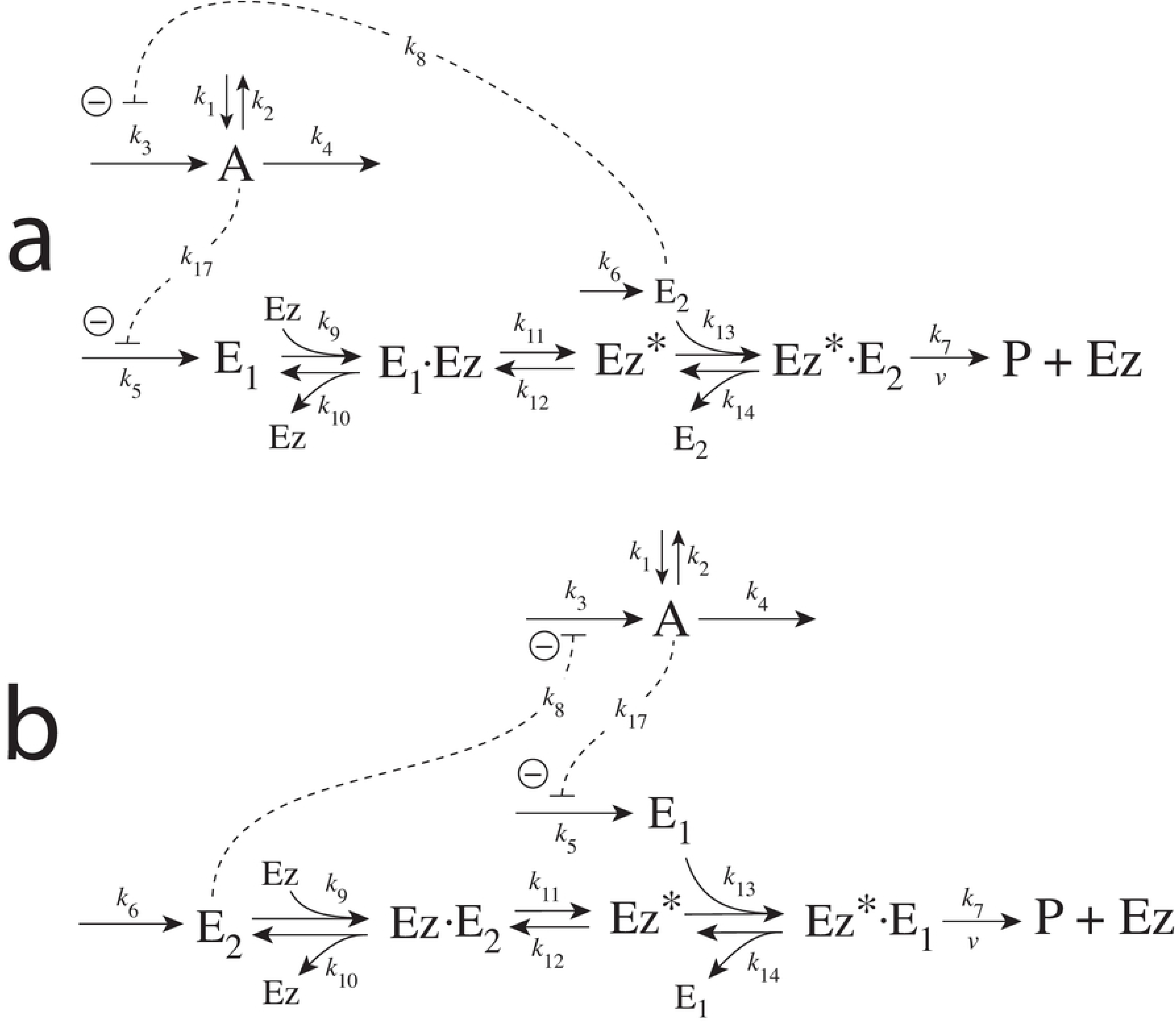
Suggested mechanism for the inflow control regulation of iron in mammalian cells. IRP2 activtes and stabilizes reactions promoting the inflow of iron into the cell. Iron activates the enzyme FBXL5 (FBXL5_i_ is an inactive form) which enables the binding of IRP2 and UB leading to ubiquitinated IRP2. In this way IRP2 is moved out of the negative feedback loop.

Under these assumptions iron is homeostatically controlled with set-point *Fe_set_*, which is determined by the condition

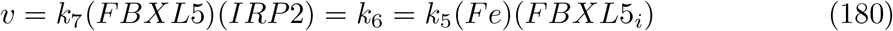

resulting in

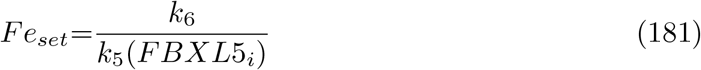

Thus, the level of iron under iron-deficient conditions is given by the ratio between the rate of IRP2 generation and the rate of FBXL5 activation.

Iron and zinc homeostasis in yeast follow analogous strategies (see Supporting Material in Ref [6]).

## Conclusion

We showed that antithetic/dual-E control can be incorporated into eight basic negative feedback motifs m1-m8. For four of them we have explicitly shown that robust antithetic control is possible when the removal of the two controller molecules *E*_1_ and *E*_2_ is catalyzed by an enzyme. Antithetic control has the advantage that it does not require specific kinetics, like zero-order kinetics is required for robust single-E control. Enzymatic dual-E control allows for the possibility that many enzymatic processes which take part in feedback regulations (like phosphorylation) may be better understood in terms of their contributions to obtain robust control. Although dual-E controllers based on ternary-complex or ping-pong mechanisms have similar (and often identical) dynamics with respect to the controlled variable, the kinetics of the participating enzymatic species are generally different for the different mechanisms. Low enzyme concentrations may limit robust homeostatic performance of catalyzed dual-E (and single-E) controllers. Transition between dual-E and single-E control may occur, but robust homeostasis for the resulting single-E controller is generally bound to zero-order kinetics. Single-E control within a dual-E network may show metastability, i.e. single-E control will switch spontaneously to dual-E control and “critical slowing down” may be observed.

Irreversibility of catalyzed (or uncatalyzed) controllers is one of the necessary conditions to obtain robust homeostasis. The work by Prigogine and coworkers [19] showed that organisms, as dissipative structures, exist as steady states far from chemical equilibrium [50]. In view of Cannon’s definition [18, 51] homeostasis preserves these steady states and thereby contributes to the stability of organisms and cells.

## Supporting information

**S1 Text. Steady state (King-Altman) expressions for enzyme-catalyzed ternary-complex and ping-pong reactions using E_1_ and E_2_ as substrates**.

## References

1. Yi TM, Huang Y, Simon MI, Doyle J. Robust perfect adaptation in bacterial chemotaxis through integral feedback control. PNAS. 2000;97(9):4649–53.

2. El-Samad H, Goff JP, Khammash M. Calcium homeostasis and parturient hypocalcemia: an integral feedback perspective. J Theor Biol. 2002;214(1):17–29.

3. Wilkie J, Johnson M, Reza K. Control Engineering. An Introductory Course. New York: Palgrave; 2002.

4. Ni XY, Drengstig T, Ruoff P. The control of the controller: Molecular mechanisms for robust perfect adaptation and temperature compensation. Biophys J. 2009;97(5):1244–53.

5. Ang J, Bagh S, Ingalls BP, McMillen DR. Considerations for using integral feedback control to construct a perfectly adapting synthetic gene network. J Theor Biol. 2010;266(4):723–738.

6. Drengstig T, Jolma I, Ni X, Thorsen K, Xu X, Ruoff P. A basic set of homeostatic controller motifs. Biophys J. 2012;103(9):2000–2010.

7. Briat C, Gupta A, Khammash M. Antithetic integral feedback ensures robust perfect adaptation in noisy biomolecular networks. Cell Systems. 2016;2(1):15–26.

8. Briat C, Zechner C, Khammash M. Design of a Synthetic Integral Feedback Circuit: Dynamic Analysis and DNA Implementation. ACS Synth Biol. 2016;5(10):1108–1116. doi:10.1021/acssynbio.6b00014.

9. Krishnan J, Floros I. Adaptive information processing of network modules to dynamic and spatial stimuli. BMC Systems Biology. 2019;13(1):32.

10. Aoki SK, Lillacci G, Gupta A, Baumschlager A, Schweingruber D, Khammash M. A universal biomolecular integral feedback controller for robust perfect adaptation. Nature. 2019; p. 1.

11. Khammash MH. Perfect adaptation in biology. Cell Systems. 2021;12(6):509–521.

12. Segel IH. Enzyme Kinetics: Behavior and Analysis of Rapid Equilibrium and Steady State Enzyme Systems. New York: Wiley; 1975.

13. Cornish-Bowden A. Fundamentals of Enzyme Kinetics. Fourth Edition. John Wiley & Sons; 2012.

14. Radhakrishnan K, Hindmarsh AC. Description and Use of LSODE, the Livermore Solver for Ordinary Differential Equations. NASA Reference Publication 1327, Lawrence Livermore National Laboratory Report UCRL-ID-113855. Cleveland, OH 44135-3191: National Aeronautics and Space Administration, Lewis Research Center; 1993.

15. Ang J, McMillen DR. Physical constraints on biological integral control design for homeostasis and sensory adaptation. Biophys J. 2013;104(2):505–515.

16. Cleland WW. The kinetics of enzyme-catalyzed reactions with two or more substrates or products: I. Nomenclature and rate equations. Biochimica et Biophysica Acta (BBA)-Specialized Section on Enzymological Subjects. 1963;67:104–137.

17. Lotka AJ. Elements of Physical Biology. Baltimore: Williams & Wilkins Company; 1925.

18. Langley, LL, editor. Homeostasis. Origins of the Concept. Stroudsbourg, Pennsylvania: Dowden, Hutchinson & Ross, Inc.; 1973.

19. Nicolis G, Prigogine I. Self-organization in Nonequilibrium Systems. From Dissipative Structures to Order through Fluctuations. John Wiley & Sons; 1977.

20. Gánti T. The Principles of Life. Oxford University Press; 2003.

21. Capra F, Luisi PL. The Systems View of Life: A Unifying Vision. Cambridge: Cambridge University Press; 2014.

22. King EL, Altman C. A schematic method of deriving the rate laws for enzyme-catalyzed reactions. J Phys Chem. 1956;60(10):1375–1378.

23. Fjeld G, Thorsen K, Drengstig T, Ruoff P. The performance of homeostatic controller motifs dealing with perturbations of rapid growth and depletion. J Phys Chem B. 2017;121:6097–6107.

24. Ruoff P, Agafonov O, Tveit DM, Thorsen K, Drengstig T. Homeostatic controllers compensating for growth and perturbations. PLoS One. 2019;14(8):e0207831.

25. Ganapathisubramanian N, Showalter K. Critical slowing down in the bistable iodate-arsenic (III) reaction. The Journal of Physical Chemistry. 1983;87(7):1098–1099.

26. Kleppe R, Waheed Q, Ruoff P. DOPA Homeostasis by Dopamine: A Control-Theoretic View. International Journal of Molecular Sciences. 2021;22(23). doi:10.3390/ijms222312862.

27. Shoval O, Goentoro L, Hart Y, Mayo A, Sontag E, Alon U. Fold-change detection and scalar symmetry of sensory input fields. PNAS. 2010; p. 201002352.

28. Drengstig T, Ni X, Thorsen K, Jolma I, Ruoff P. Robust adaptation and homeostasis by autocatalysis. J Phys Chem B. 2012;116(18):5355–5363.

29. Cohen P. The origins of protein phosphorylation. Nature Cell Biology. 2002;4(5):E127–E130.

30. Wang Z, Cole PA. Catalytic mechanisms and regulation of protein kinases. Methods in Enzymology. 2014;548:1–21.

31. Dunlap J. Molecular bases for circadian clocks. Cell. 1999;96(2):271–290.

32. Huang G, Chen S, Li S, Cha J, Long C, Li L, et al. Protein kinase A and casein kinases mediate sequential phosphorylation events in the circadian negative feedback loop. Genes & Development. 2007;21(24):3283–3295.

33. Wang B, Kettenbach AN, Zhou X, Loros JJ, Dunlap JC. The phospho-code determining circadian feedback loop closure and output in *Neurospora*. Molecular Cell. 2019;74(4):771–784.

34. Baker CL, Loros JJ, Dunlap JC. The circadian clock of *Neurospora crassa*. FEMS Microbiology Reviews. 2012;36(1):95–110.

35. Baker CL, Kettenbach AN, Loros JJ, Gerber SA, Dunlap JC. Quantitative proteomics reveals a dynamic interactome and phase-specific phosphorylation in the *Neurospora* circadian clock. Molecular Cell. 2009;34(3):354–363.

36. Edery I, Zwiebel LJ, Dembinska ME, Rosbash M. Temporal phosphorylation of the *Drosophila* period protein. PNAS. 1994;91(6):2260–2264.

37. Rosbash M, Bradley S, Kadener S, Li Y, Luo W, Menet J, et al. Transcriptional feedback and definition of the circadian pacemaker in *Drosophila* and animals. In: Cold Spring Harbor Symposia on Quantitative Biology. vol. 72. Cold Spring Harbor Laboratory Press; 2007. p. 75–83.

38. Liu Y, Merrow M, Loros JJ, Dunlap JC. How temperature changes reset a circadian oscillator. Science. 1998;281(5378):825–829.

39. Rensing L, Mohsenzadeh S, Ruoff P, Meyer U. Temperature Compensation of the Circadian Period Length - A Special Case Among General Homeostatic Mechanisms of Gene Expression? Chronobiology International. 1997;14(5):481–498.

40. Rensing L, Ruoff P. Temperature effect on entrainment, phase shifting, and amplitude of circadian clocks and its molecular bases. Chronobiology International. 2002;19(5):807–864.

41. Ruoff P, Loros JJ, Dunlap JC. The relationship between FRQ-protein stability and temperature compensation in the Neurospora circadian clock. PNAS. 2005;102(49):17681–17686.

42. Hu Y, Liu X, Lu Q, Yang Y, He Q, Liu Y, et al. FRQ-CK1 Interaction Underlies Temperature Compensation of the Neurospora Circadian Clock. mBio. 2021;12(3):e01425–21.

43. Tanaka K, Asami T, Yoshida S, Nakamura Y, Matsuo T, Okamoto S. Brassinosteroid homeostasis in Arabidopsis is ensured by feedback expressions of multiple genes involved in its metabolism. Plant Physiology. 2005;138(2):1117–1125.

44. He JX, Gendron JM, Sun Y, Gampala SS, Gendron N, Sun CQ, et al. BZR1 is a transcriptional repressor with dual roles in brassinosteroid homeostasis and growth responses. Science. 2005;307(5715):1634–1638.

45. Wei Z, Li J. Regulation of brassinosteroid homeostasis in higher plants. Frontiers in Plant Science. 2020;11:1480.

46. He JX, Gendron JM, Yang Y, Li J, Wang ZY. The GSK3-like kinase BIN2 phosphorylates and destabilizes BZR1, a positive regulator of the brassinosteroid signaling pathway in Arabidopsis. PNAS. 2002;99(15):10185–10190.

47. Ciechanover A. The ubiquitin-proteasome proteolytic pathway. Cell. 1994;79(1):13–21.

48. Vashisht AA, Zumbrennen KB, Huang X, Powers DN, Durazo A, Sun D, et al. Control of iron homeostasis by an iron-regulated ubiquitin ligase. Science. 2009;326(5953):718–721.

49. Zheng N, Schulman BA, Song L, Miller JJ, Jeffrey PD, Wang P, et al. Structure of the Cul1–Rbx1–Skp1–F box Skp2 SCF ubiquitin ligase complex. Nature. 2002;416(6882):703–709.

50. Karsenti E. Self-organization in cell biology: A brief history. Nature Reviews Molecular Cell Biology. 2008;9(3):255–262.

51. Cannon W. Organization for physiological homeostatics. Physiol Rev. 1929;9:399–431.

